# Potent *in vitro* anti-SARS-CoV-2 activity by gallinamide A and analogues via inhibition of cathepsin L

**DOI:** 10.1101/2020.12.23.424111

**Authors:** Anneliese S. Ashhurst, Arthur H. Tang, Pavla Fajtová, Michael Yoon, Anupriya Aggarwal, Alexander Stoye, Mark Larance, Laura Beretta, Aleksandra Drelich, Danielle Skinner, Linfeng Li, Thomas D. Meek, James H. McKerrow, Vivian Hook, Chien-Te K. Tseng, Stuart Turville, William H. Gerwick, Anthony J. O’Donoghue, Richard J. Payne

## Abstract

The emergence of SARS-CoV-2 in late 2019, and the subsequent COVID-19 pandemic, has led to substantial mortality, together with mass global disruption. There is an urgent need for novel antiviral drugs for therapeutic or prophylactic application. Cathepsin L is a key host cysteine protease utilized by coronaviruses for cell entry and is recognized as a promising drug target. The marine natural product, gallinamide A and several synthetic analogues, were identified as potent inhibitors of cathepsin L activity with IC_50_ values in the picomolar range. Lead molecules possessed selectivity over cathepsin B and other related human cathepsin proteases and did not exhibit inhibitory activity against viral proteases Mpro and PLpro. We demonstrate that gallinamide A and two lead analogues potently inhibit SARS-CoV-2 infection *in vitro*, with EC_50_ values in the nanomolar range, thus further highlighting the potential of cathepsin L as a COVID-19 antiviral drug target.

## Introduction

The novel severe acute respiratory syndrome (SARS)-like coronavirus-2 (SARS-CoV-2) is a beta-coronavirus responsible for COVID-19, a respiratory disease that emerged from the Hubei province in China in December 2019 and was declared a global pandemic on the 11^th^ of March 2020 (*1*). Alarmingly, in less than 12 months, COVID-19 has been reported in nearly every country, with over 77 million confirmed COVID-19 cases and more than 1.7 million deaths globally (at the time of writing). The COVID-19 pandemic has had a devastating effect on both global health and the functioning of society, and prophylactic and therapeutic intervention strategies are urgently needed. To date, there has been an intense global effort directed towards the development of an effective COVID-19 vaccine, with several candidates, (*2*) including two mRNA vaccines and a chimpanzee adenovirus-vectored vaccine recently completing phase III human clinical trials (*3–5*). In addition to an effective prophylactic vaccine, the control of COVID-19, and potentially future SARS coronavirus zoonoses, also requires efficacious antiviral therapeutics. While antiviral drug discovery for SARS-CoV-2 has been the subject of significant effort, the majority of molecules currently in clinical trials are repurposed from other indications for which they were originally approved. Remdesivir is currently the only antiviral drug to be approved by the U.S Food and Drug Administration (FDA) for the treatment of COVID-19. Whilst this molecule has been reported to show some efficacy during early infection, the drug has performed poorly in a number of trials where it was deemed ineffective, (*6, 7*) including a recent report from the World Health Organization that suggested remdesivir provided little to no effect in the outcome of COVID-19 infections in hospitalized patients (*8*). Examples of other repurposing approaches that have been explored, include the use of the hydroxychloroquine, (*9–11*) the HIV therapy Lopinavir-ritonavir, (*12, 13*) as well as type I interferon regimens; (*14, 15*) however, these too have shown no improvement over standard care in hospitalized COVID-19 patients. At the present time, one of the most effective means of improving COVID-19 patient outcomes has been through the use of the glucocorticoid dexamethasone which serves to reduce inflammation-mediated lung injury (*16*). Taken together, despite significant effort from the global research community, there is still an urgent need to discover effective antivirals for COVID-19 infection that operate through novel mechanisms of action and that are distinctive to the molecules currently in use. While there are obvious benefits to post-exposure drug therapies, the development of a pre-exposure prophylaxis approach, such as what has been employed for HIV, (*17*) would also be transformative for protecting vulnerable communities.

SARS-CoV-2 shares 82% genome identity to the SARS-CoV that emerged in Guangdong province, China in 2002. It is now accepted that the two coronaviruses share similar molecular mechanisms of host cell recognition, entry and replication (*18–20*). Recognition and entry of SARS-CoV-2 into target cells relies on binding between the receptor-binding domain (RBD) of an envelope homotrimeric spike glycoprotein (S) and the host cellular receptor, angiotensin-converting enzyme 2 (ACE2) (**Figure 1A**). Recently, a second receptor protein, a transmembrane glycoprotein of the immunoglobulin superfamily, known as CD147, has been identified as mediating spike protein interaction and viral uptake via endocytosis (*21*). Each monomeric unit of the S protein contains an S1 and S2 subunit that mediate attachment and membrane fusion with host cells, respectively. Host cell entry requires priming of the S protein by cleavage at the S1/S2 and the S2ʹ site that enables fusion of viral and cellular membranes. This cleavage has been shown to be carried out primarily by the membrane-bound serine protease TMPRSS2, (*19*) but can also be performed by the cysteine protease, cathepsin L (CatL) (*22, 23*). Following entry of the virus into the cell via an endosomal pathway, CatL is responsible for S1 cleavage at acidic pH, conditions where TMPRSS2 is not catalytically functional. Overexpression of CatL in human cell lines enhances SARS-CoV-2 spike-mediated viral entry, while circulating CatL is elevated during COVID-19 disease and correlates with progression and severity (*24*). Finally, it is known that expression of CatL, but not cathepsin B (CatB), is up-regulated by interleukin-6 (IL-6) (*25*). This is of importance, as there is early evidence that IL-6 is a surrogate inflammatory marker for severe COVID-19 disease with poor prognosis (*26*). This would imply that CatL is upregulated under these conditions and would therefore be a potentially important target for controlling excessive pathology.

**Figure 1.**
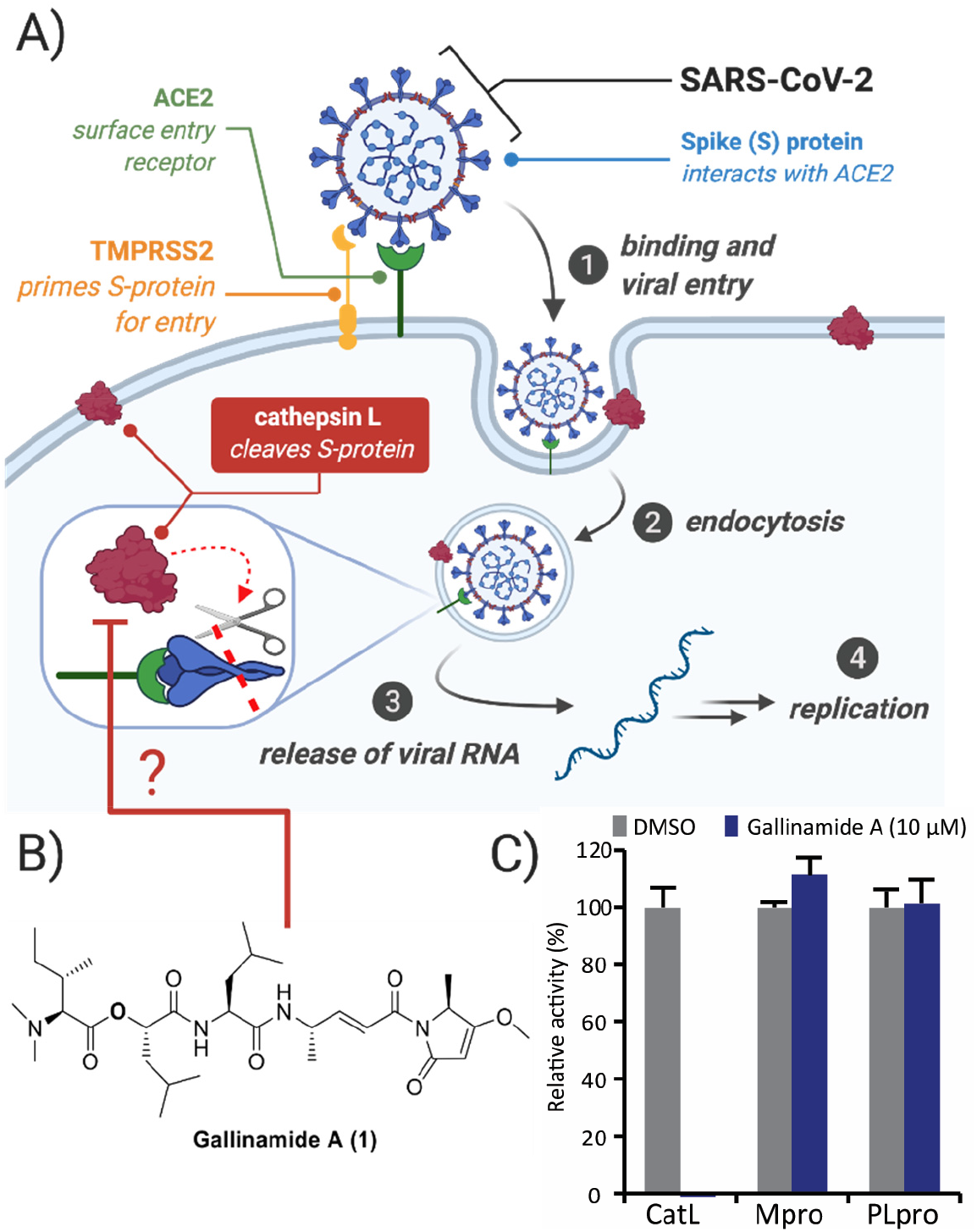
Targeting cathepsin L with gallinamide A to inhibit SARS-CoV-2. (**A**) Role of host proteases in cell entry mechanisms for SARS-CoV-2 (endocytic entry route shown). (**B**) Structure of the natural product depsipeptide CatL inhibitor gallinamide A (**1**). (**C**) Enzymatic activity of CatL, Mpro and PLpro following incubation with 10 μM gallinamide A. Assays were performed in triplicate wells and relative activity was compared to reactions containing 0.2% DMSO.

This dual role of CatL in the establishment and progression of COVID-19 pathology has therefore reinforced the enzyme as a key drug target for SARS-CoV-2 (*27*). The importance of both TMPRSS2 and CatL for facilitating viral entry and replication is highlighted by the effectiveness of TMPRSS2 inhibitors, camostat mesylate (*19*) and nafamostat mesylate (*28*) and the pan-cysteine protease inhibitors E64d (*19*) and K777 (*23*) to reduce virus infection levels of SARS-CoV-2 (and SARS-CoV) in a range of human cell lines. Following uncoating and release of the viral RNA from the endosome, translation of the two large viral reading frames gives rise to the polyproteins pp1a and pp1ab that are subsequently processed by two viral proteases: the main protease (Mpro) and the papain-like protease (PLpro) (*18*). This gives rise to a number of non-structural proteins (nsps) that subsequently orchestrate viral replication and release from infected cells, to infect new cells. As such, both Mpro and PLpro are also promising antiviral targets for SARS-CoV-2 (*29–32*).

Gallinamide A **1** (also known as symplostatin 4) is a modified depsipeptide natural product that was independently discovered from marine cyanobacteria of the *Schizothrix* genus in Panama (*33*) and *Symploca* in Florida (*34*). The natural product has several unusual structural features including a pyrrolinone derived from L-alanine and an α,β-unsaturated imide moiety (**Figure 1B**). Notably, gallinamide A has been demonstrated to be a potent covalent inhibitor of several parasite-derived cysteine proteases (*35–37*), as well as human CatL (*38*), with several synthetic analogues of the natural product also shown to have potent *in vivo* antimalarial activity in a murine model (*39*). Given the importance of CatL for cellular entry of SARS-CoV-2, we sought to investigate whether the CatL inhibitory activity of gallinamide A could be leveraged for SARS-CoV-2 antiviral activity. Towards this end, in May 2020 we assembled an international consortium to investigate gallinamide A, together with 32 synthetic natural product analogues, as novel inhibitors of SARS-CoV-2 entry. We report herein that several analogues exhibit potent inhibitory activity against CatL with IC_50_ values in the low nanomolar to picomolar range. Several of these gallinamide A-inspired molecules also possess selectivity over CatB and other related cathepsin proteases and do not inhibit the two viral proteases Mpro and PLpro. Finally, we demonstrate that gallinamide A and two of the most active CatL-inhibiting natural product analogues potently inhibit SARS-CoV-2 infection *in vitro*, with EC_50_ values in the nanomolar range, thus further highlighting the potential of CatL as a COVID-19 antiviral target.

## Results

### Gallinamide A inhibits SARS-CoV-2 infection *in vitro* via the inhibition of CatL

Gallinamide A (**1**) was initially screened in a SARS-CoV-2 viral infection assay in Vero 76 clone E6 cells (VeroE6) at the National Institute of Allergy and Infectious Disease (NIAID) to gauge whether the natural product exhibited antiviral activity. Pleasingly, gallinamide A was shown to decrease viral load with an IC_90_ of 88 nM. Following this, we further validated the activity of the natural product by performing the *in vitro* SARS-CoV-2 infectivity assay in our own laboratories using VeroE6 cells and adenocarcinomic human alveolar basal epithelial cells overexpressing ACE2 (A549/ACE2). Briefly gallinamide A (**1**) was pre-incubated with 4000 VeroE6 cells in a 384-well plate for 30 minutes and then challenged with 0.5 MOI SARS-CoV-2. In the absence of treatment, these conditions lead to viral cytopathic effects (CPE) and cell loss that is correlated to the level of available infectious virus in the initial inoculum after 72 hours. Addition of inhibitors that block viral entry reduces CPE/cell loss in a dose-dependent manner. To obtain a quantitative measure of CPE, the nuclei of live cells were stained, imaged and enumerated using high-content image analysis software. Using this method, we could readily generate dose-dependent sigmoidal inhibition curves and calculated an EC_50_ of 28 ± 12 nM for gallinamide A from four independent experiments (**Figure S1).** In parallel, 500 SARS-CoV-2 viral particles were used to infect confluent monolayers of A549/ACE2 cells grown in 96-well plates that had been incubated for 1 hr with serially diluted gallinamide A, followed by assessment of the formation of CPE 96 hrs later under an inverted microscope. This showed that gallinamide A at a concentration as low as 625 nM was capable of completely preventing virus-induced CPE.

Recent studies have shown that peptidic inhibitors of CatL and other related cysteine proteases also possess inhibitory activity against SARS-CoV-2 Mpro (*40, 41*). In order to assess the comparative activity of gallinamide A against human CatL and the viral proteases Mpro and PLpro, we incubated these proteases with 10 μM of the natural product, and quantified the remaining activity using fluorogenic substrates. CatL activity was completely inhibited by gallinamide A at this concentration, while no inhibition of Mpro or PLpro was observed (**Figure 1C**).

We next sought to determine the abundance of CatL in the VeroE6 and A549 cell lines. A comprehensive bottom-up proteomic analysis was utilized to assess the likelihood that this protease serves as the physiological target of gallinamide A in cell-based assays (**Figure S2, Data File S1**). In VeroE6 cells, CatL was among the top 300 most abundant proteins and in comparison, its closely related homologue cathepsin B (CatB), was 10-fold lower in abundance. In contrast, A549 cells had 10-fold higher levels of CatB compared to CatL. In addition to determining the abundance of CatL by proteomics, we confirmed that CatL was catalytically active in VeroE6 cells by incubating lysates with the fluorogenic substrate Z-Phe-Arg-AMC. This activity was completely inhibited by 20 μM of Gallinamide A. CatB can also cleave Z-Phe-Arg-AMC, however when a CatB-specific inhibitor, CA-074(*42*) was added to the lysate, no inhibitory activity was observed (**Figure S3A**). These data reveal that there is very little active CatB in VeroE6 lysates relative to CatL. For A549 cell extracts, protease activity using Z-Phe-Arg-AMC was also inhibited by gallinamide A, however most of the activity was also sensitive to CA-074 indicating that CatB is the dominant cysteine cathepsin in these cells (**Figure S3B**). This activity data is supported by our proteomics analysis of A549 cells where CatB was found to be more abundant than CatL (**Figure S2**). Taken together, these data suggest that the antiviral activity of gallinamide A is not due to inhibition of viral (MPro and PLpro) protease targets but likely due to inhibition of host-derived CatL. The level of CatL differs substantially in these cell lines which supports published data that some cell lines are more sensitive to cysteine protease inhibition than others (*23*).

### Assessment of CatL inhibitory activity of synthetic gallinamide A analogues

Three series of synthetic gallinamide A analogues have been developed by our laboratories to date. Series 1 consist of depsipeptide compounds **2-10**, with various aliphatic residues incorporated at the pseudo-*N*-terminus, the α,β-unsaturated moiety and on the pyrrolinone ring (**Figure 2**) (*37*). Series 2 is comprised of indolylpyrrolinone analogues **11-23** that have varied functionality at the pseudo-*N*-terminus and the three amino acid residues within the linear chain (*39*). Finally, series 3 analogues (**24-33**) possess a number of biaryl-moieties appended to the pyrrolinone unit, and dimethylvaline or methylpiperidine functionalities at the pseudo-*N*-terminus. Each of these analogues was subjected to a preliminary inhibitory screen against human CatL, Mpro and PLpro (**Figure 3**). For the CatL assays, compounds were incubated at 416 nM; the majority of the analogues showed potent inhibition at this concentration, with 21 of the compounds reducing enzymatic activity by more than 90% (**Figure 3A**). Based on the lack of potency of the parent natural product **1** against Mpro and PLpro, compounds were screened at a higher concentration (10 μM) against these enzymes. As with gallinamide A, none of these analogues showed appreciable inhibition of SARS-CoV-2 Mpro or PLpro under these conditions (**Figure 3B**). A follow-up screen against CatL at 13 nM identified gallinamide A (**1**), analogue **3** from series 1, and **17**, **19**, **20** and **23** from series 2 as being the most potent inhibitors. These were selected for more detailed inhibitory assays against human CatL as described below.

**Figure 2:**
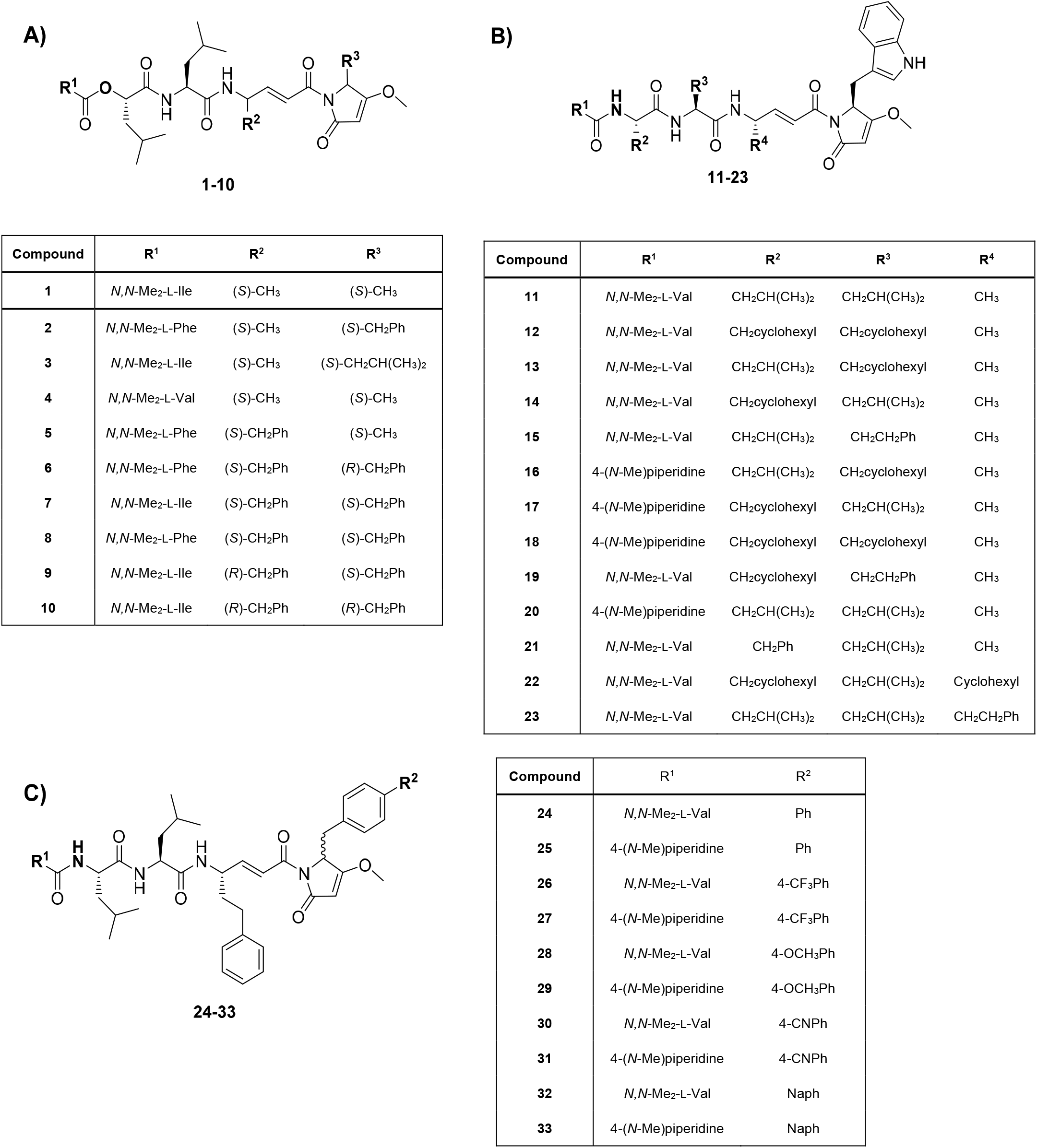
Structures of gallinamide A natural product analogues. (**A**) Gallinamide A **1** and depsipeptide gallinamide A analogues **2-10** with variation at the pseudo-*N*-terminus and the α,β-unsaturated and pyrrolinone units; (**B**) Indolylpyrrolinone gallinamide A analogues **11-23**; (**C**) Biaryl-functionalized pyrrolinone analogues **24-33**.

**Figure 3.**
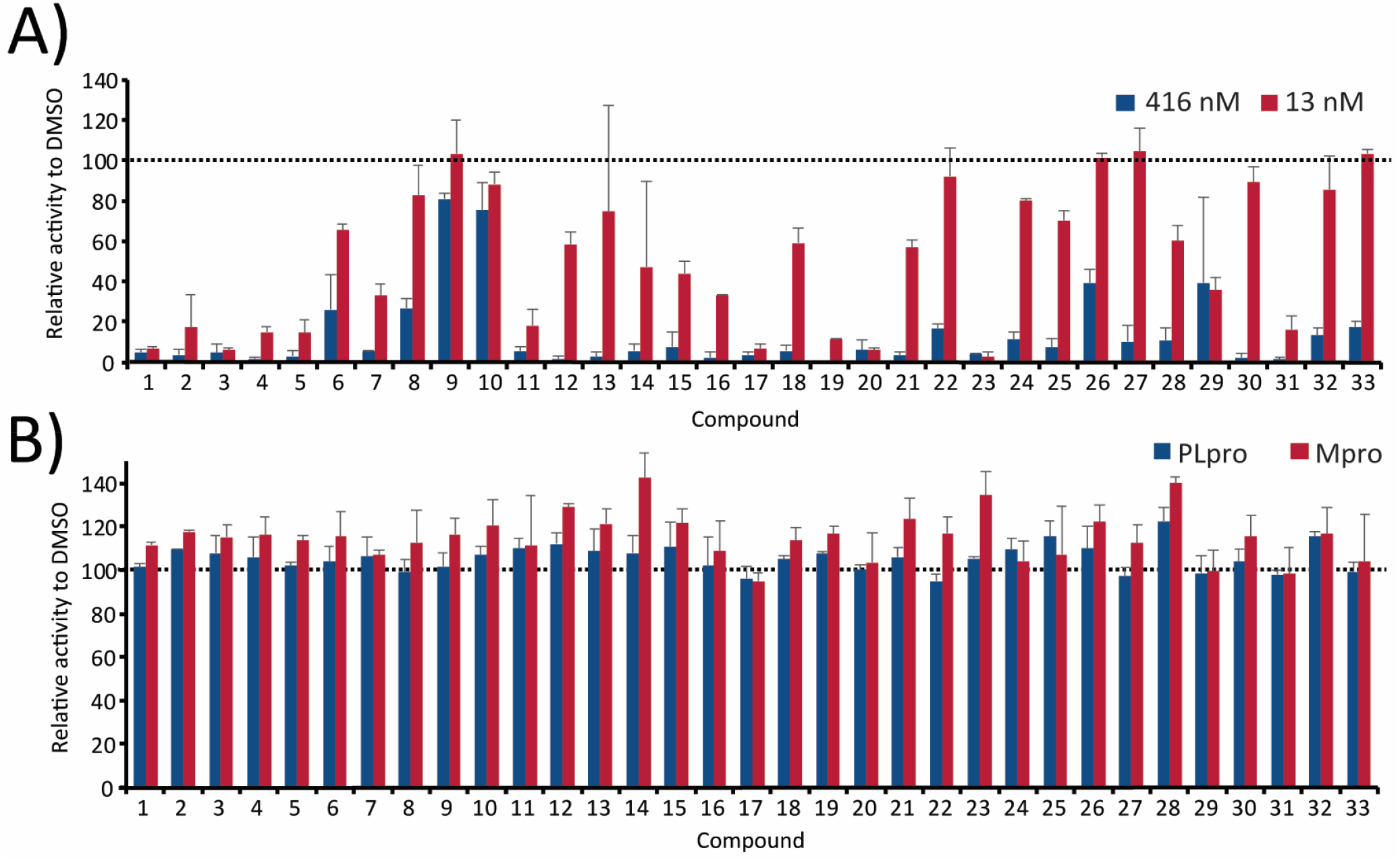
Activity of gallinamide A and analogues against cellular and viral proteases. (**A**) Screening of gallinamide A **1** and analogues **2-33** at 13 nM and 416 nM for inhibition of human CatL; (**B**) Screening of gallinamide A and analogues at 10 μM for inhibition against SARS-CoV-2 Mpro and PLpro. Assays were performed in triplicate wells and data are the means +/− SD. Activity was compared to reactions containing 0.2% DMSO in assay buffer.

The inhibitory potency of gallinamide A (**1**) and analogues **17**, **19**, **20** and **23** was next evaluated using dose-response assays against CatL for the determination of IC_50_ values (**Figure 4**). An insufficient amount of analogue **3** was available for these experiments, and therefore no further studies were performed with this compound. Gallinamide A (**1**) exhibited an IC_50_ of 17.6 pM against CatL (**Figure 4A**), while the four analogues possessed IC_50_ values ranging from 6 to 17 pM (**Figure 4B-E**). Each of the compounds were counter-screened against related human cysteine cathepsins, including CatB, cathepsin V (CatV), cathepsin K (CatK) and cathepsin S (CatS) (**Figure 4A-E**). While each of the compounds showed inhibitory activity against all cathepsins tested, the IC_50_ values were generally between 1 and 4 orders of magnitude less potent than for CatL, thus indicating selectivity for CatL over the other cathepsins. In addition to determining the IC_50_ values, we also determined k_inact_/K_i_ values for gallinamide A and the lead analogues for CatL **(Figure 4F**, **Figure S4**). Based on the k_inact_/K_i_ constants, we determined that analogue **23** was 1.4-fold more potent than the parent natural product **1**, while **17**, **19** and **20** were between 2.1- and 3.2-fold less potent than **1**.

**Figure 4.**
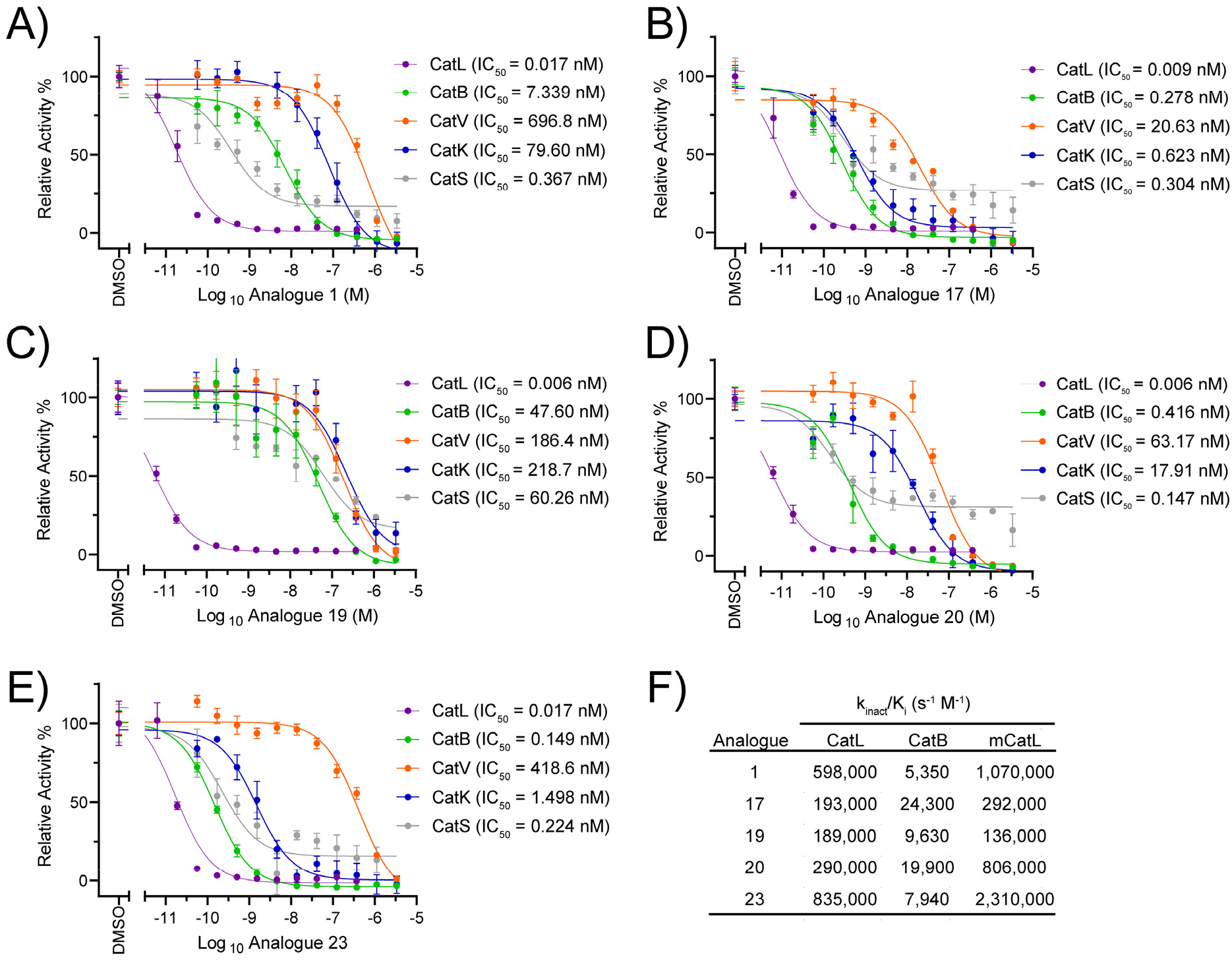
Dose response curves and corresponding IC_50_ values against CatL, CatB, CatV, CatK and CatS for (**A**) gallinamide A (**1**) and synthetic analogues (**B**) **17**; (**C**) **19**; (**D**) **20** and (**E**) **23**. Activity was normalized to a vehicle control and all data are the means ± SD for three technical replicates. (**F**) k_inact_/K_i_ (s^−1^ M^−1^) values for the inhibition of human CatL, human CatB and mouse CatL (mCatL) by gallinamide A (**1**) and analogues **18**, **20**, **21** and **23**.

Given that CatB was detected in our proteomics studies using VeroE6 and A549 lysates (**Figure S2**), and it has also been recently reported that a broad-spectrum activity-based probe for cysteine proteases was active against both proteases in cell lysates, (*23*) we also calculated the k_inact_/K_i_ values for gallinamide A and the selected lead analogues against CatB. This revealed selectivity values ranging from 8-fold for compound **17** to 112-fold for gallinamide A (**Figure 4F, Figure S5**) for CatL over CatB. In addition, we evaluated the molecules against mouse CatL (mCatL) to give an initial indication of the future applicability in mouse models of SARS-CoV-2 infection. With the exception of **19**, the compounds possessed increased potency against mCatL relative to human CatL (**Figure 4F, Figure S6**).

### Inhibition of SARS-CoV-2 Entry into Cells

Having established the potent inhibition of CatL and SARS-CoV-2 entry by gallinamide A, and having also demonstrated that synthetic analogues **17**, **19**, **20** and **23** were also potent inhibitors of the protease, we next assessed these lead analogues against SARS-CoV-2 entry and infectivity in VeroE6 cells. Before embarking on the antiviral assays, each of the analogues were first assessed for cytotoxicity on VeroE6 cells. Pleasingly, none of the compounds exhibited cytotoxicity after 24 hours at up to a concentration of 100 μM in this cell line (**Figure S7**). Antiviral assays were next performed in VeroE6 cells on the lead analogues using the same high content fluorescence microscopy method described for gallinamide A above. Analogues **19** and **23** both exhibited potent antiviral activity with EC_50_ values of 168 nM and 920 nM, respectively (**Table 1, Figure S8**). Based on these data, we further assessed the activity of the compounds against SARS-CoV-2-induced CPE in A549/ACE2 cells. In these experiments complete inhibition of CPE was observed at concentrations of 310 nM for **19** and **23** (**Figure S9**), making them more potent than gallinamide A in these cells. Interestingly, in both cell lines, **17** and **20** exhibited poor antiviral activity with EC_50_ values greater than 5 μM, despite showing potent inhibition of CatL *in vitro* (IC_50_ of 9 and 6 pM, respectively) and complete inactivation of CatL in VeroE6 and A549 lysates at 10 μM (**Figure S3**). The reduced activity of **17** and **20** compared to **1**, **19** and **23** in the antiviral assay may be owing to poor cell permeability of these compounds.

**Table 1.** EC_50_ values for Gallinamide A (1) and analogues 17, 19, 20 and 23 against SARS-CoV-2 infection in VeroE6 cells. The percentage inhibition of CPE was determined by high content fluorescence microscopy. Data are the mean ± SD of 3-4 independent biological replicates (see **Figure S1** and **Figure S9** for data).

| Compound | EC<sub>50</sub> SARS-CoV-2 (μM) |
| --- | --- |
| Gallinamide A (1) | 0.028 ± 0.012 |
| **17** | 6.07 ± 2.4 |
| **19** | 0.168 ± 0.067 |
| **20** | > 5 |
| **23** | 0.920 ± 0.43 |

While both CatL and TMPRSS2 can mediate SARS-CoV-2 entry into target cells by facilitating S protein priming, (*22*) evidence suggests that the predominant mechanism of entry is dependent on the relative levels of the two proteins in a given cell line (*19, 28, 41*). To assess the effect of the lead compounds **1** and **19** in a TMPRSS2 overexpressing cell line, we used MRC-5/ACE2/TMPRSS2 cells which we also verified to possess CatL, albeit in lower abundance than in VeroE6 and A549 cells (**Figure S2**). Gratifyingly, both **1** and **19** were able to completely neutralize SARS-CoV-2 entry into this cell line at a concentration of 32 nM (**Figure S10**), thus suggesting that CatL is still important for viral entry in these TMPRSS2 overexpressing cells. Interestingly, while the TMPRSS2 inhibitor Nafamostat mesylate was able to block infection in MRC-5/ACE2/TMPRSS2 (albeit with neutralization at 4 μM concentrations), it did not exhibit notable antiviral activity on VeroE6 cells **(Figure S10)**. These data suggest that both TMPRSS2 and CatL protease pathways are operational and important for infectivity in MRC-5/ACE2/TMPRSS2 cells, but that CatL is the predominate pathway in VeroE6 cells, consistent with the levels of each protein in these cells (**Figure S2**).

## Discussion

Pathogenic coronaviruses are now well recognized for their potential to induce substantial human morbidity and mortality. To bring the current global pandemic of SARS-CoV-2 under control will ultimately require mass implementation of a safe and effective vaccine. Whilst recent vaccine results show promise, challenges remain with respect to protecting unvaccinated individuals or vulnerable populations. In previous epidemics, where vaccines were limited, either by efficacy or reach within the community, antivirals served as the cornerstone for fighting disease. This has been recently observed for two major viral pathogens, HCV, which is now readily curable with a short course of antivirals, and HIV. Whilst HIV cannot be cured by therapeutic approaches, it can be readily managed through pre-exposure prohylaxis (PreP) or treatment post-exposure with antivirals. While it is acknowledged that respiratory diseases like SARS-CoV-2 present many unique challenges with respect to antiviral strategies, it is feasible that equivalent success in PreP development could in turn lead to the protection of vulnerable communities, such as healthcare workers or the elderly within care facilities. In addition, PreP-based approaches may reduce the viral reproductive rate in communities with high SARS-CoV-2 prevalence. Even if PreP antiviral strategies were limited to lowering viral load, this would improve not only prognosis but also limit the onward spread to the next host.

Early in pandemics, clinically ready drugs provide the fastest means for effective treatment. However, to date, many repurposed drugs have not provided any evidence of clinical benefit for SARS-CoV-2, with the exception of dexamethasone, a steroid rather than an antiviral agent, for late-stage disease. With the virus entrenched within the global population, and new more infectious strains emerging (*43*), we now require compounds that target the virus with greater potency and breadth. Previous studies have indicated that CatL inhibitors have the potential to serve as antivirals for a range of coronaviruses, including HCoV-229E (*44*), MERS-CoV (*45*) and SARS-CoV-1 (*46*). SARS-CoV-2 spike-mediated cell entry is enhanced by overexpression of CatL in human cell lines, and recent evidence suggests elevated levels of circulating CatL is also correlated with COVID-19 disease progression and severity (*24*). Importantly, targeting a host protease, as opposed to a viral protease specific to one virus or viral strain, may provide the best opportunity for broad spectrum antiviral activity that is less susceptible to rapidly progressing viral mutations. CatL is therefore a promising drug target for SARS-CoV-2, and also for other current or future coronavirus strains, as well as other virus families including the *Filoviridae*, such as the Ebola virus (*45*).

Historically, nature has been the single most successful source for the discovery of new pharmaceutical lead compounds, especially in the areas of anti-infective and anticancer agents (*47*). In this work, we demonstrate that gallinamide A, a peptide-based natural product isolated from a marine cyanobacterium, and two synthetic analogues of the natural product, possess potent activity against SARS-CoV-2 infectivity in monkey (VeroE6) and human (A549) cells. Specifically, the parent natural product exhibited the most potent inhibition of SARS-CoV-2 in VeroE6 cells, with an EC_50_ of 28 nM, whereas analogues **19** and **23** were the most potent in A549/ACE2 cells (complete inhibition of CPE at 310 nM). We show that these molecules were able to inhibit CatL in both cell lines, and could selectively target recombinant CatL over other related cysteine cathepsins *in vitro,* with no host cell cytotoxicity up to 100 μM. Based on this potent CatL inhibitory activity of gallinamide A, and the structural analogues, it is tempting to speculate that the evolution of this cyanobacterially-derived molecule has been guided and optimized by activity at a cysteine protease target related to CatL.

CatL is a lysosomal enzyme that plays a key role in intracellular protein degradation in our cells. Expression of this enzyme is dysregulated in several human diseases, including cancer (*48*), arthritis (*49*) and diabetic nephropathy (*50*), with therapeutics in development for these ailments (*51*). Mice deficient for this gene exhibit hair loss (*52*), bone and heart defects (*53, 54*), and enhanced susceptibility to bacterial infection (*55*). Therefore, targeting of this host enzyme for treatment of SARS-CoV-2 infections may pose a risk for side-effects. However, treatment for coronavirus infection would likely be short-term, aiming to reduce viral entry and early replication until host innate and adaptive responses can be developed. There are already a number of FDA-approved drugs that possess inhibitory activity against CatL (*27*). Furthermore, recent studies have shown that K777, a potent irreversible inhibitor of CatL consisting of a dipeptide-vinyl sulfone, was found to be well tolerated in rodents, dogs and nonhuman primates (*56*), and an IND has been opened with the FDA for its use as a therapeutic treatment of COVID-19 infection (*23*). Importantly, synthetic gallinamide A analogues have already been safely examined *in vivo* for other infectious indications (*39*). Our studies showed that gallinamide A and the most promising synthetic analogues maintain potency against mouse CatL and therefore future work in our laboratories will investigate *in vivo* efficacy in SARS-CoV-2 infection models.

While CatL shows clear promise as a suitable drug target, the entry pathways for SARS-CoV-2 are complex and can be endosomal or at the cell membrane, with the latter driven by the serine protease TMPRSS2 (*19*). Tropism of the virus is equally complex. Oropharyngeal tissue is well known for its ability to facilitate transmission and contributes to the early stages of disease, whilst once established, infection can proceed across many organ types outside of the respiratory tract, including kidneys, liver, heart, brain, and blood (*57*). CatL and TMPRSS2 are both highly expressed in lung tissue (*58*), however treatment of infection across multiple different cell types and organ systems may require preferential inhibition at the membrane (TMPRSS2) or in the endosome (CatL). Future studies will therefore also seek to incorporate inhibitors of the major fusogenic entry pathways for the virus. This combination approach may have an added benefit, in that it may allow lower doses of the individual drugs to be utilized due to synergism in the mechanisms of action.

In summary, we demonstrate here that the marine natural product, gallinamide A and several synthetic analogues, are potent inhibitors of CatL, a key host cysteine protease involved in the pathogenesis of SARS-CoV-2. Lead molecules possessed selectivity over CatB and other related human cathepsin proteases and did not exhibit inhibitory activity against viral proteases Mpro and PLpro. Gallinamide A and two lead analogues potently inhibited SARS-CoV-2 infection *in vitro*, which now serve as leads for the future development of therapeutic or prophylactic drug candidates. Importantly, this work also highlights the potential of CatL as a bone fide target for the development of novel antivirals for SARS-CoV-2 or other pathogenic coronaviruses.

## Materials and Methods

### Protease screening of gallinamide A (1) and analogues 2-33

Recombinant SARS-CoV-2 Mpro was expressed and purified as described previously.(*23*) 200 nM of this enzyme was incubated with 10 μM of each compound at 25 °C in 20 mM Tris-HCl pH 7.5, 150 mM NaCl, 1 mM DTT, 5% glycerol, 0.01% Tween-20 for 15 minutes. An equal volume of 200 μM Mu-HSSKLQ-AMC (Sigma-Aldrich, SCP0224) in the same buffer was then added and protease activity was quantified at 25 °C. Recombinant SARS-CoV-2 PLpro was purchased from Acro Biosystems, PAE-C5184. 49 nM of this enzyme was incubated with 10 μM of each compound at 25 °C for 15 minutes in 50 mM HEPES, 150 mM NaCl, 1 mM DTT, 0.01% Tween 20; pH 6.5. An equal volume of 50 μM Z-RLRGG-AMC (Bachem, I1690) in the same buffer was then added to start the reaction. Recombinant human Cathepsin L was purchased from R&D Systems (952-CY), and 50 pM of the enzyme was incubated with 416 nM and 13 nM of each compound and separately with 10 μM of gallinamide A at 25 °C for 15 minutes in 50 mM sodium acetate pH 5.5, 5 mM DTT, 0.01% BSA, 1 mM EDTA. An equal volume of 5 μM of Z-FR-AMC (Sigma-Aldrich, C9521) was then added. All assays were performed in triplicate wells and DMSO was used as a vehicle control. The final volume of each reaction was 30 μL, and fluorescence was measured at 360/460 nm (ex/em) using a Biotek® Synergy M2 fluorescence plate reader. Activity was normalized to wells lacking inhibitor but containing 0.01% DMSO in assay buffer.

### Dose-response studies

Recombinant human cathepsin B (953-CY), cathepsin S (1183-CY), cathepsin V (1080-CY) and were purchased from R&D Systems. Human cathepsin K (BML-SE553) was purchased from Enzo. 1.11 nM of cathepsin L, cathepsin B, cathepsin V, cathepsin K and 4.44 nM of cathepsin S were preincubated for 30 minutes at 25 ^o^C with 3333 to 0.006 nM of each compound in 40 mM sodium acetate pH 5.5, 5 mM DTT, 0.001% BSA, 1 mM EDTA, 100 mM NaCl. Remaining protease activity was then detected using a final concentration of 40 μM of Z-FR-AMC (Sigma-Aldrich, C9521). Fluorescent activity was recorded as outlined above. For k_inact_/K_i_ studies, mouse cathepsin L (1515-CY) was purchased from R&D Systems. Human CatL, human CatB and mouse CatL (1 nM) were assayed at 25 °C in 30 μL reaction volumes containing 40 mM sodium acetate pH 5.5, 5 mM DTT, 0.001% BSA, 1 mM EDTA, 100 mM NaCl, 40 μM of Z-FR-AMC (Sigma-Aldrich, C9521) in the presence of 555.56 to 0.01 nM of each compound in quadruplicate wells. To calculate the IC_50_, inhibitor and substrate were added simultaneously to the enzyme and the rate of AMC release was determined from 0 to 30 min and compared to a DMSO control. To calculate K_i_ and k_inact_, the rate of product formation was calculated at 6 min intervals for 1 h and normalized to activity in the control wells containing DMSO. k_obs_ values for each inhibitor concentration were calculated from inactivation curves, and inhibition constants kinact and k_inact_/K_i_ were calculated by nonlinear regression of k_obs_ and inhibitor concentration using GraphPad Prism 9 software. A K_i_ value from the Morrison equation was determined using GraphPad Prism 9 software Y=V_o_*(1-((((E_t_+X+(K_i_*(1+(S/K_m_))))-(((E_t_+X+(K_i_*(1+(S/K_m_))))^2)-4*E_t_*X)^0.5))/(2*E_t_)) using the fixed values of enzyme concentration (Et) = 0.001 μM, substrate concentration (S) = 40 μM, and K_m_ = 38.1 μM for human CatB, and K_m_ = 2.2 μM for human CatL and mouse CatL.(*59*)

### Cytotoxicity assays

Cytotoxicity of lead compounds was determined by an Alamar Blue HS (Invitrogen) cell viability assay as per manufacturer’s instructions. Briefly, VeroE6 (ATCC® CRL-1586™) cells were grown in Dulbecco’s Modified Eagle Medium (DMEM) with 4.5 g/L D-Glucose, L-Glutamine and 110 mg/L sodium pyruvate (Gibco), 10% foetal calf serum (Sigma) and penicillin-streptomycin (100 U/ml, Gibco). 5×10^4^ cells were seeded into wells of a 96-well flat bottom culture plate (Corning) and once adhered, compounds were added at varying concentrations and incubated for 24 hours (37 ^o^C, 5% CO_2_). Alamar Blue HS cell viability reagent (Invitrogen) was added and the cells further incubated for 2-4 hours. Relative fluorescent units (RFU) were determined per well at ex/em 560/590 nm (Tecan Infinite M1000 pro plate reader). Increasing RFU is proportional to cell viability.

### Cell lines for SARS-CoV-2 infection assays

VeroE6 cells (ATCC® CRL-1586™) were maintained in Minimal Essential Media (Invitrogen) supplemented with 10% FBS and subcultured according to the supplier’s instructions. MRC-5 cells stably expressing human ACE2 and TMPRSS2 were generated by transducing MRC-5 cells (ATCC CCL-171) with lentiviral particles. Briefly, the ORFs for hACE2 (Addgene#1786) and hTMPRSS2a (Addgene#53887) were cloned into lentiviral expression vectors pRRLsinPPT.CMV.GFP.WPRE (*60*) and pLVX-IRES-ZsGreen (Clontech) respectively. Lentiviral particles expressing the above proteins were produced by co-transfecting expression plasmids individually with a second generation lentiviral packaging construct psPAX2 (courtesy of Dr Didier Trono through NIH AIDS repository) and VSVG plasmid pMD2.G (Addgene#2259) in HEK293T cells (Life Technologies) by using polyethyleneimine as previously described (*61*). Two successive rounds of lentiviral transductions were then performed on MRC-5 cells to generate MRC-5/ACE2/TMPRSS2. Human A549 cells stably transduced with human ACE2 viral receptor (A549/ACE2) were grown in M-10 medium.

### Proteomics and Activity Assays using Cell Lysates

Cell lysates were digested to peptides and purified using a protocol described previously (*62*). Peptide mixtures were analysed by nanoflow LC-MS/MS as described. Identification and quantification were performed using MaxQuant with a 1% false discovery rate at the protein level using a target-decoy approach. The MRC5 and A549 cells were analysed using the *Homo sapiens* proteome database and the VeroE6 cells using the *Chlorocebus sabaeus* proteome database, both of which were from Uniprot.

VeroE6 and A549 cells were lysed by sonication in 50 mM MES pH 6.0; 100 μM AEBSF, 1 μM pepstatin A, 1 mM EDTA, 1 mM DTT buffer and protein concentration in the cleared supernatant was quantified using the bicinchoninic acid assay with BSA as a protein standard. 225 ng of total protein was incubated with 20 μM of **1**, **17**, **19**, **20**, **23**, CA-074 (MilliporeSigma #205530) and E-64 (Alfa Aesar #J62933) for 30 minutes in 50 mM sodium acetate pH 5.5; 5 mM DTT; 1 mM EDTA; 1 μM pepstatin; 100 μM AEBSF, and then mixed with equal volume of 50 μM of Z-Phe-Arg-AMC in the same buffer. Activity was normalized to wells that lacked inhibitor but contained 0.2% DMSO in assay buffer.

### SARS-CoV-2 virus inhibition assay

A high content fluorescence microscopy approach was used to assess the ability of gallinamide A and the synthetic analogues to inhibit SARS-CoV-2 infection in permissive cells. The compounds were initially diluted in cell culture medium (MEM-2% FCS) to make 2x working stock solutions and then serially diluted further in the above media to achieve a 2-fold dilution series. 20 μL of each dilution was then added to cells seeded a day prior at 3×10^3^ cells in a volume of 40 μL of MEM-2% FCS per well in a 384-well plate (Corning #CLS3985). The plates containing the cells and compounds were incubated for 30 minutes at 37 °C, 5% CO_2_ following which 20 μL of virus solution at 8×10^3^ TCID50/mL (*63*) was added to the wells. Plates were incubated at 37 °C, 5% CO_2_ for a further 72 hours following which the cells were stained with NucBlue dye (Invitrogen, #R37605) according to manufacturer’s instructions. The entire 384-well plate was imaged using the InCell 2500 (Cytiva) high throughput microscope, with a 10× 0.45 NA CFI Plan Apo Lambda air objective. Acquired nuclei were then counted in high content using InCarta high-content image analysis software (Cytiva) to give a quantitative measure of CPE. Virus inhibition/neutralization was calculated as %N= (D-(1-Q))x100/D, where; “Q” is the value of nuclei in test well divided by the average number of nuclei in untreated uninfected controls, and “D”=1-Q for the wells infected with virus but untreated with inhibitors. Thus, the average nuclear counts for the infected and uninfected cell controls get defined as 0% and 100% neutralization respectively. To account for cell death due to drug toxicity, cells treated with a given compound alone and without virus were included in each assay. The % neutralization for each compound concentration in infected wells was normalised to % neutralization in wells with equivalent amount of compound but without the virus to yield the final neutralization values for each condition.

At the University of Texas Medical Branch, A549/ACE2 cells were grown in M-10 media and treated with 500 SARS-CoV-2 viral particles (USA_WA1/2020 isolate) prior to addition of 20 to 0.156 μM of compound. After cultivation at 37 °C for 4 days, individual wells were observed by an inverted microscope for determining the status of virus-induced formation of cytopathic effect (CPE). The efficacy of individual compounds was calculated and expressed as the lowest concentration capable of completely preventing virus-induced CPE in 100% of the wells. All compounds were dissolved in 100% DMSO as 10 mM stock solutions and diluted in culture media and experiments using infectious virus were conducted under BSL-3 conditions.

## General

We thank B. Hurst (NIAID) for initial evaluation of gallinamide A in a SARS-CoV-2 assay. The SARS-CoV-2 Mpro expression plasmid was kindly provided to us from Rolf Hilgenfeld, University of Lübeck, Germany. We appreciate support from the UCSD Chancellor’s office through an initiative to create a campus wide small molecule resource library. We acknowledge the assistance of Jason Hsu, Vivian Tat, and Rayavara Kempaiah for assistance with tissue culture. We would also like to thank Dr Luke Dowman for helpful suggestions.

## Author contributions

A.S.A., W.H.G., A.J.O and R.J.P. initiated and designed the project and managed the consortium. A.S.A. and M.L. performed cell-based assays, cytotoxicity experiments and proteomics and analyzed the data. A.H.T. and A.S. synthesized gallinamide A (**1**) and analogues **11-33** and analyzed the NMR data. P. F., M.Y., L.B., L.L, T.D.M and V.H. performed the enzymatic assays and analyzed the data. D.S., P.F. and J.H.M. cloned, expressed and purified SARS-CoV-2 Mpro. A.A. and S.T. performed the *in vitro* SARS-CoV-2 assays on VeroE6 and MRC5 cells and analyzed the data. A.D. and C-T.T. performed the *in vitro* SARS-CoV-2 assays on A549 cells and analyzed the data. A.S.A., A.T., W.H.G., A.J.O and R.J.P. wrote the manuscript and prepared the figures with input from all authors.

