## Supplementary Materials for "Potent *in vitro* anti-SARS-CoV-2 activity by gallinamide A and analogues via inhibition of cathepsin L"

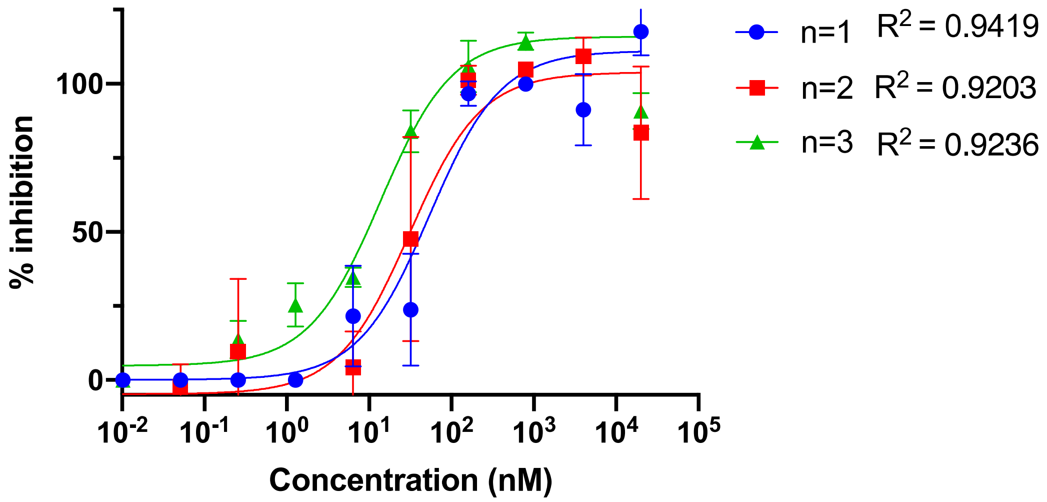


**Figure S1.** Activity of Gallinamide A for inhibition of SARS-CoV-2 infection in VeroE6 cells. Individual data points plus error bars, with lines of best fit, are shown for 3 replicate independent experiments.

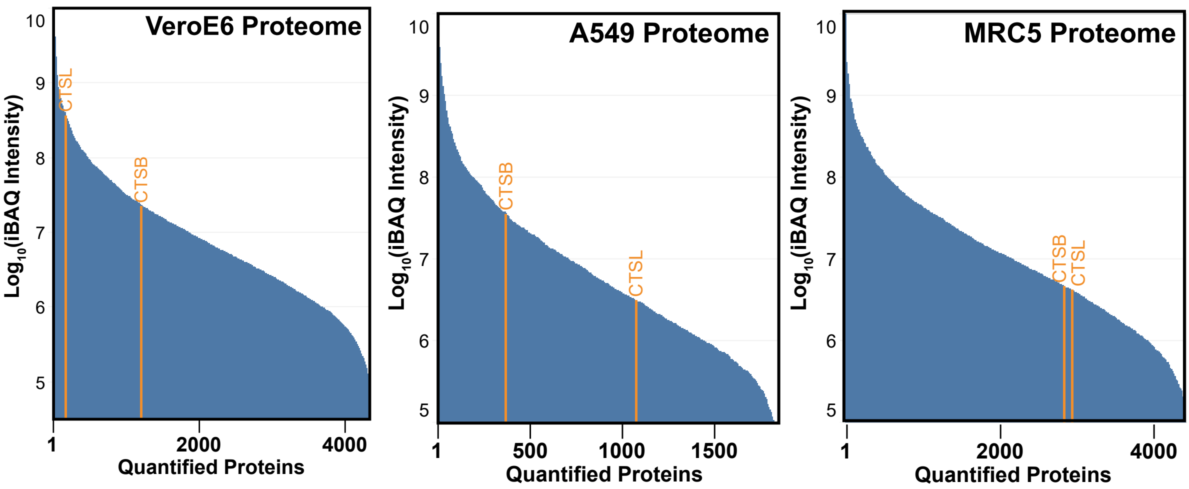


**Figure S2.** Ranked abundance plot of Cathepsin L (CTSL) and Cathepsin B (CTSB) in VeroE6, A549 or MRC5 cells expressing ACE2 and TMPRSS2, from single-shot bottom-up proteomics analysis. Protein rank for each cell line is derived the from their iBAQ intensity (y-axis), which normalises for each protein's ability to generate LC-MS/MS compatible tryptic peptides. Proteins of interest are highlighted in orange.

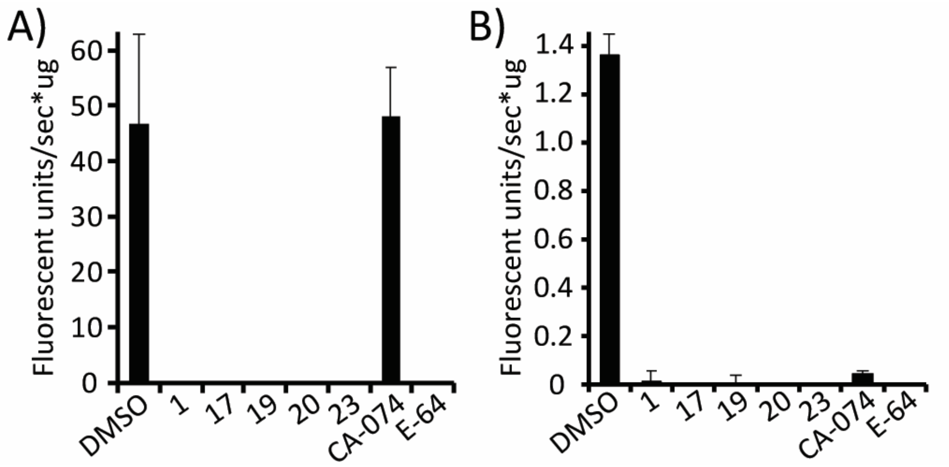


**Figure S3**. Quantitation of protease activity in (**A**) VeroE6 and (**B**) A549 cell lysates using Z-Phe-Arg-AMC. 20 µM of each Cathepsin L inhibiting compound, the selective Cathepsin B inhibitor CA-074, cysteine protease inhibitor E-64, or vehicle control (DMSO), was incubated with the cell lysate for 30 minutes prior to detection of fluorescent activity. Data are the means +/- SD for technical triplicates.

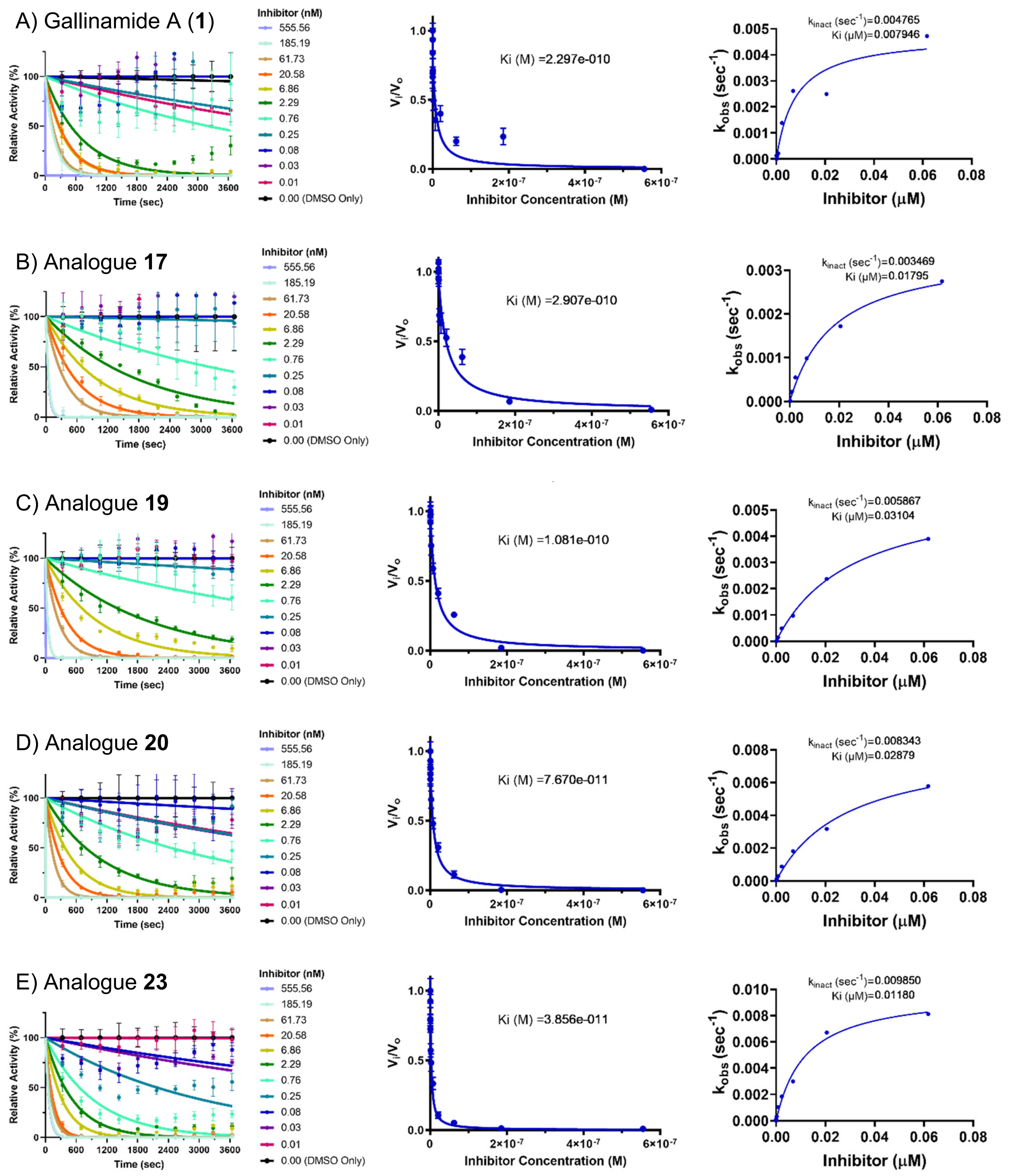


**Figure S4**. Calculation of $K_{obs}$ (left panel), calculation of $K_i$ using the Morrison equation fitting (middle panel) and $K_{inact}/K_i$ (right panel) for human cathepsin L assayed with Gallinamide A (**1**) and analogues **17**, **19**, **20** and **23**.

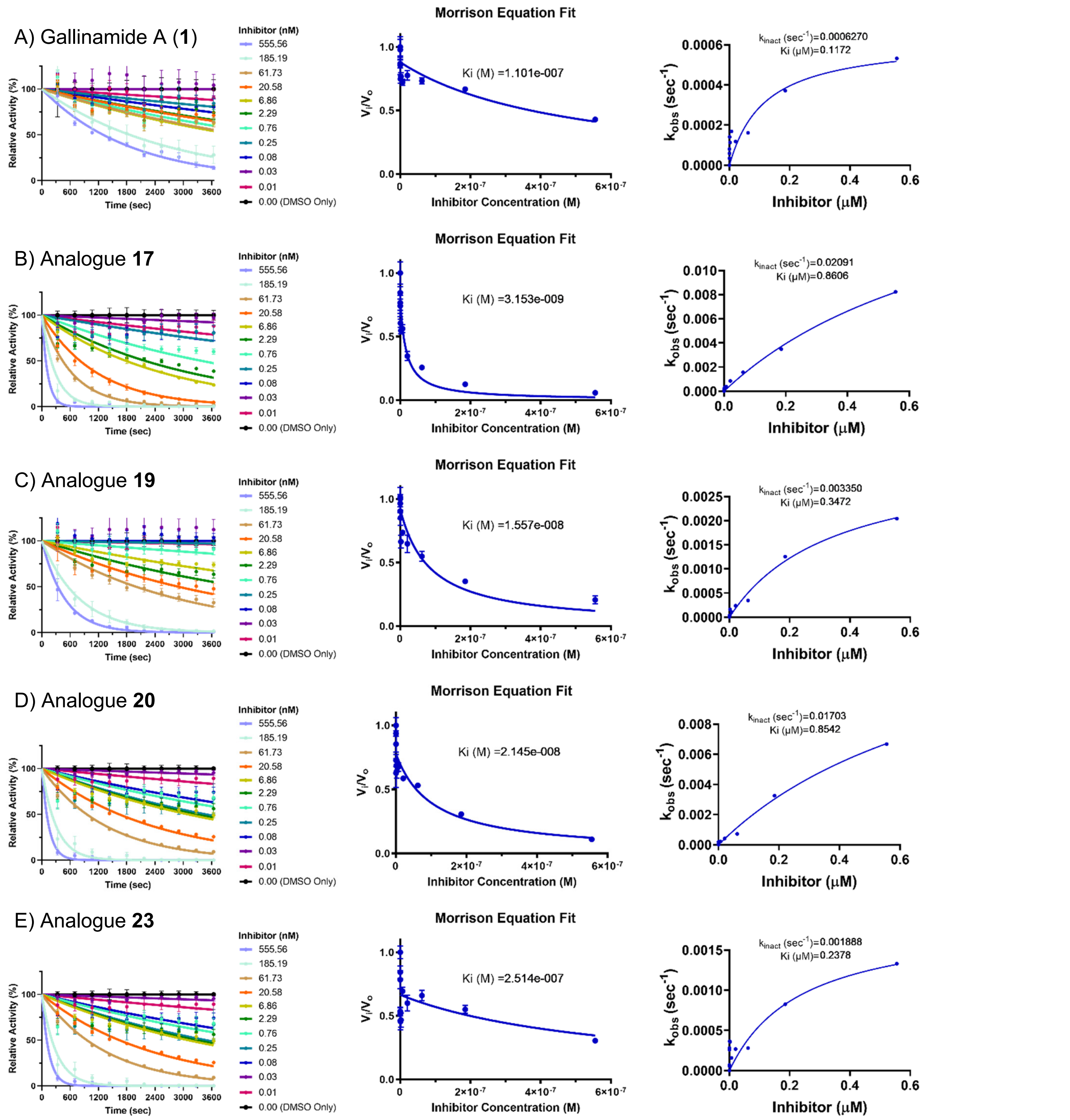


**Figure S5**. Calculation of $K_{obs}$ (left panel), calculation of $K_i$ using the Morrison equation fitting (middle panel) and $K_{inact}/K_i$ (right panel) for human cathepsin B assayed with Gallinamide A (**1**) and analogues **17**, **19**, **20** and **23**.

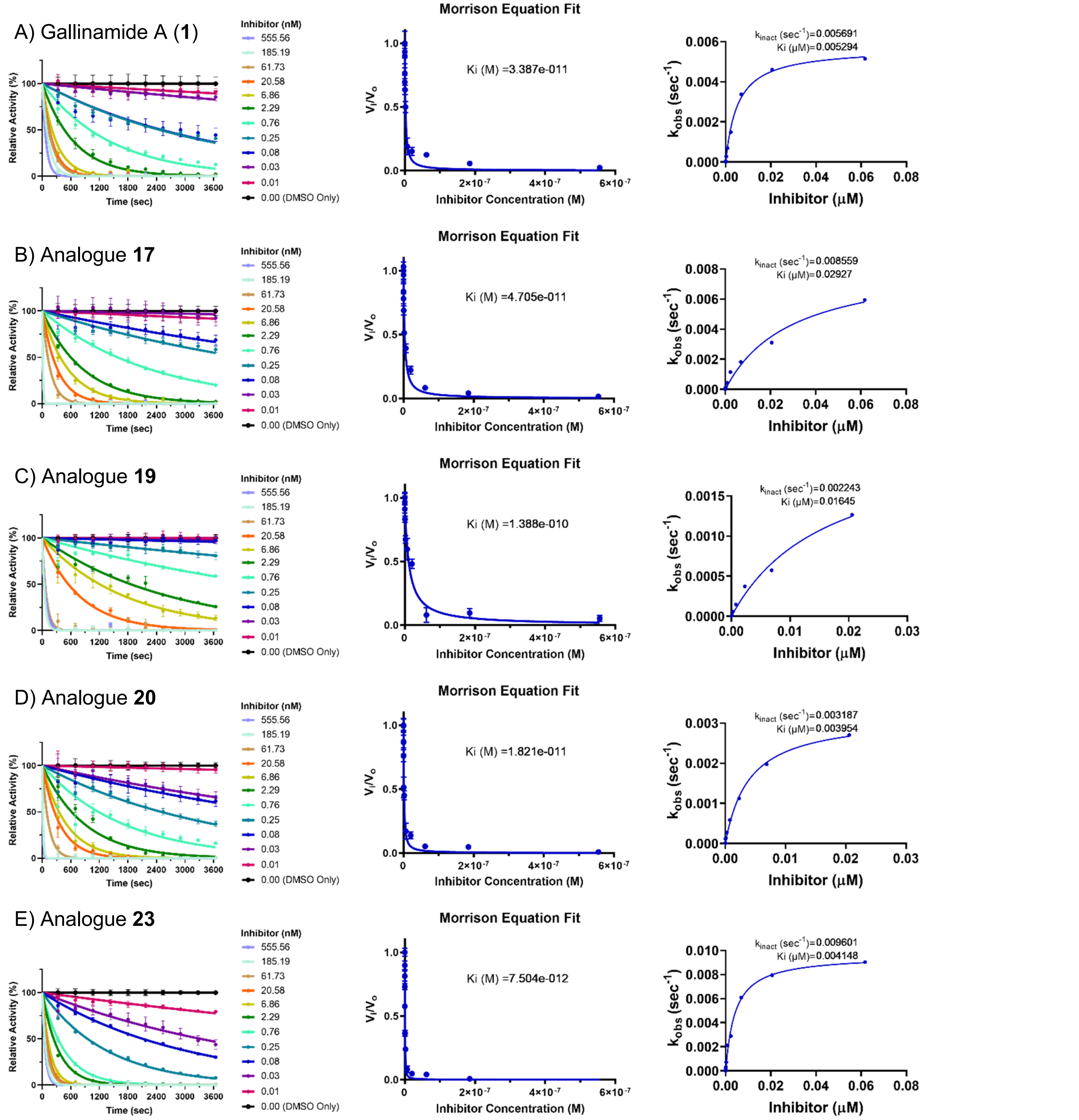


**Figure S6**. Calculation of $K_{obs}$ (left panel), calculation of $K_i$ using the Morrison equation fitting (middle panel) and $K_{inact}/K_i$ (right panel) for mouse cathepsin L assayed with Gallinamide A (**1**) and analogues **17**, **19**, **20** and **23**.

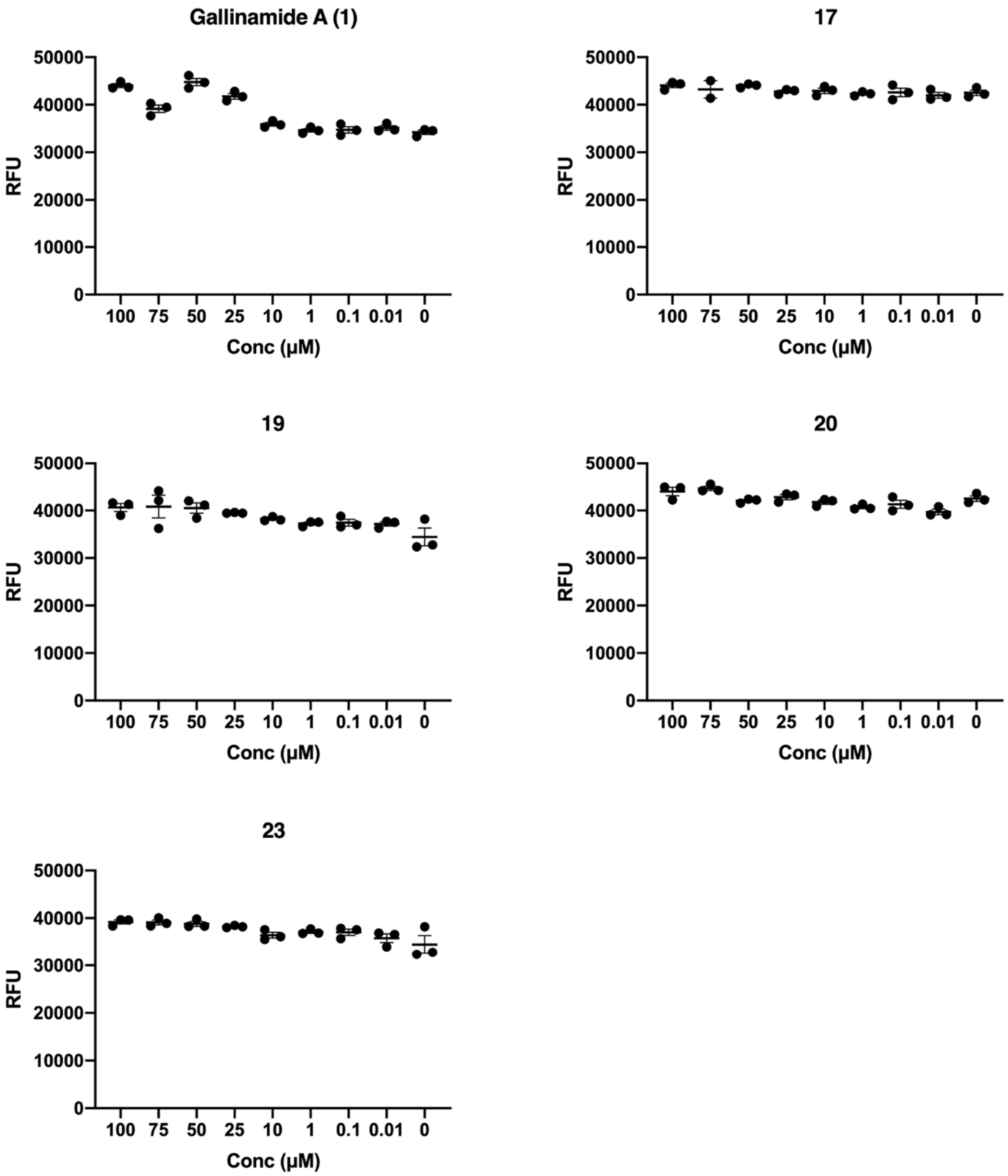


**Figure S7.** Cytotoxicity of lead compounds after 24-hour incubation with VeroE6 cells, determined by Alamar Blue HS assay. Data are the means ± SEM of technical triplicates for relative fluorescent units (RFU; Ex/Em 560/590 nm) and are representative of two independent biological replicates.

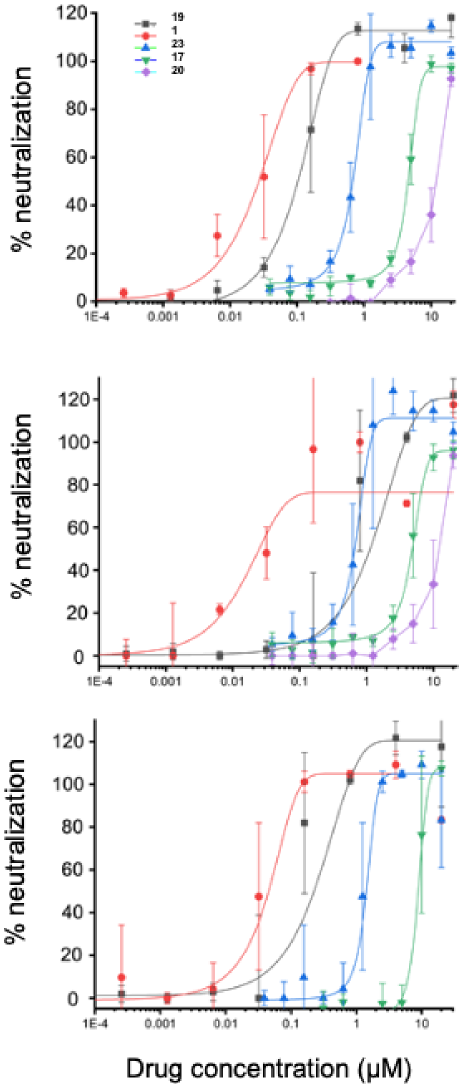


| Compound | $EC_{50}$ SARS-CoV-2 (nM)<br>Independent biological replicates | | | | Mean (nM) | SD (nM) |
|---|---|---|---|---|---|---|
| Gallinamide A (**1**) | 26.65 | 24.94 | 45.46 | 12.32 | 28.344 | 12.0439 |
| **17** | 4532.09 | 4852.88 | 8846.27 | | 6077.08 | 2403.54665 |
| **19** | 102.36 | 113.6 | 224.9 | 252.27 | 168.01667 | 67.17509 |
| **20** | >5000 | >5000 | | | N/A | N/A |
| **23** | 676.4 | 665.53 | 1418.13 | | 920.02 | 431.41 |

**Figure S8.** Determination of $EC_{50}$ for CatL inhibitors for SARS-CoV-2 in VeroE6 cells. SARS-CoV-2 inhibition data for lead compounds are shown as independent biological replicates. Individual data points and error bars, with lines of best fit, are provided followed by the summarized $EC_{50}$ values.

| | CPE for technical duplicates | | | | | | | | | | | |
|---|---|---|---|---|---|---|---|---|---|---|---|---|
| CatL inhibitor conc (μM) | **1** | | **17** | | **19** | | **20** | | **23** | | Viral Ctrl | Mock Ctrl |
| 20 | ND | ND | CPE | ND | ND | ND | ND | ND | ND | ND | CPE | ND |
| 10 | ND | ND | CPE | CPE | ND | ND | CPE | ND | ND | ND | CPE | ND |
| 5 | ND | ND | CPE | CPE | ND | ND | CPE | CPE | ND | ND | CPE | ND |
| 2.5 | ND | ND | CPE | CPE | ND | ND | CPE | CPE | ND | ND | CPE | ND |
| 1.25 | ND | ND | CPE | CPE | ND | ND | CPE | CPE | ND | ND | CPE | ND |
| 0.625 | ND | ND | CPE | CPE | ND | ND | CPE | CPE | ND | ND | CPE | ND |
| 0.31 | CPE | ND | CPE | CPE | ND | ND | CPE | CPE | ND | ND | CPE | ND |
| 0.156 | CPE | CPE | CPE | CPE | CPE | ND | CPE | CPE | CPE | ND | CPE | ND |

**Figure S9.** Inhibition of SARS-CoV-2 infection by gallinamide A and lead synthetic analogues on A549/ACE2 expressing cells. The presence or absence of cytopathic effect (CPE) was determined by microscopy. ND - no CPE. Viral Ctrl - positive control (not-treated with CatL inhibitor, infected cells), Mock Ctrl - negative control (not-treated with CatL inhibitor, not infected).

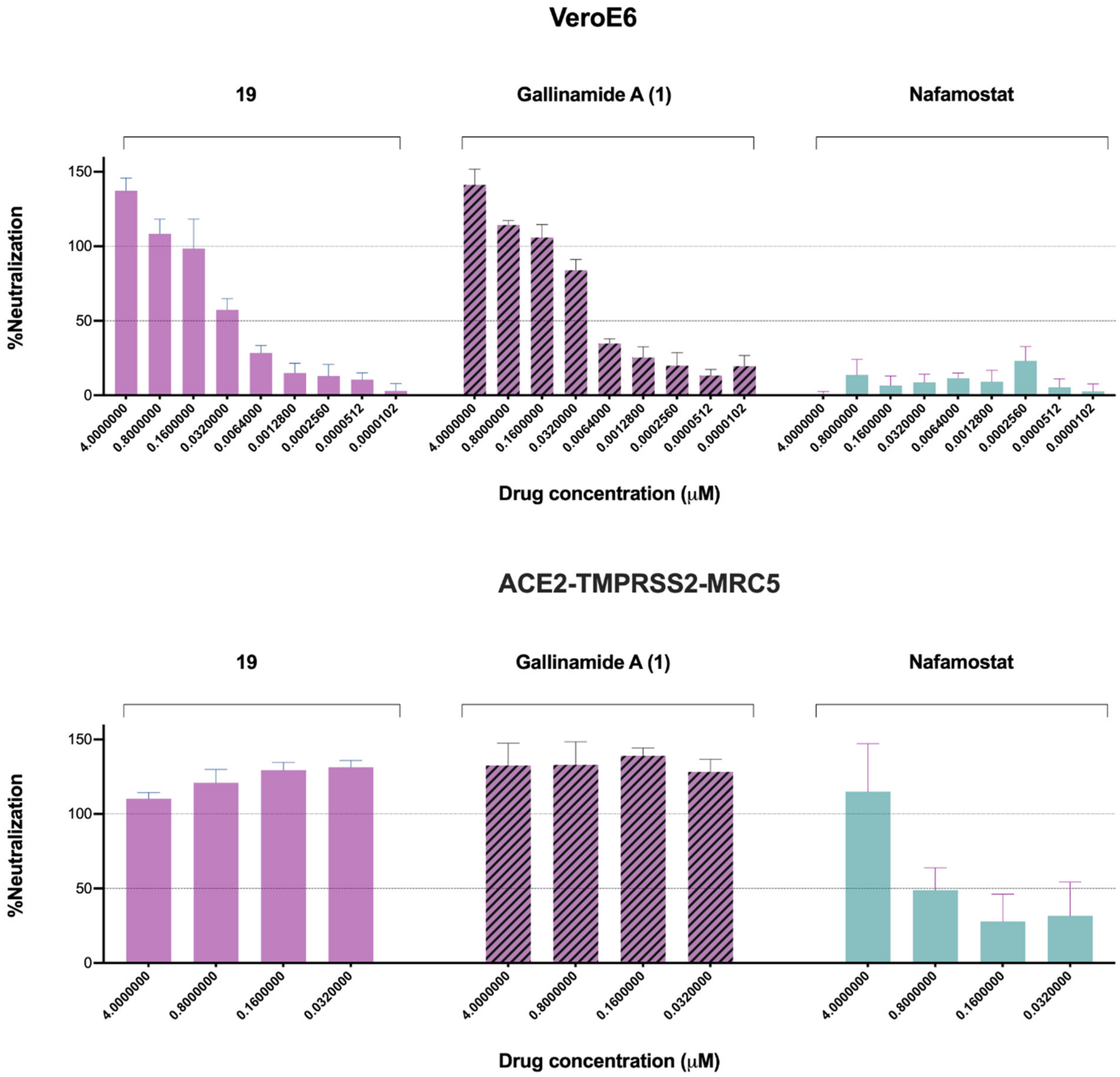


**Figure S10.** Inhibition of SARS-CoV-2 infection by lead CatL inhibitor compounds gallinamide A (**1**) and **19**, in comparison to the TMPRSS2 protease inhibitor nafamostat mesylate, directly compared in VeroE6 and ACE2/TMPR22S2 expressing MRC-5 cells. Data are the means ± SD.

### Supplementary Materials and Methods

#### Section S1: General methods for chemical synthesis of gallinamide A and gallinamide A analogues

All reagents and solvents were used as received unless otherwise noted. Anhydrous MeOH, DMF and $CH_2Cl_2$ were obtained from a PURE SOLV$^{TM}$ solvent dispensing unit. Unless otherwise stated, solution-phase reactions were carried out under an atmosphere of dry nitrogen or argon and at room temperature (22 °C). Reactions undertaken at -78 °C utilized a bath of dry ice and acetone. Reactions carried out at 0 °C employed a bath of water and ice. Reactions carried out at above room temperature utilized a magnetic stirrer with heating and a multi-well heating mantle for direct heating of the reaction flask.

Flash column chromatography was performed using 230–400 mesh Kieselgel 60 silica eluting with gradients as specified. Analytical thin layer chromatography (TLC) was performed on commercially prepared silica plates (Merck Kieselgel 60 0.25 mm F254). Visualization of TLC plates was undertaken with ultraviolet (UV) light at λ = 254 nm and staining with solutions of vanillin or phosphomolybdic acid, followed by exposure of the stained plates to heat.

$^{1}H$ NMR, $^{13}C$ NMR and 2D NMR spectra were recorded at 300 K using a Bruker DRX500, DRX400 or DRX300 spectrometer. Chemical shifts are reported in parts per million (ppm) and are referenced to solvent residual signals: DMSO- $d_6$ δ 2.50 [$^{1}H$] and δ 39.52 [$^{13}C$], $CD_3CN$ δ 1.94 [$^{1}H$] and δ 1.32 [$^{13}C$], $CDCl_3$ δ 7.26 [$^{1}H$] and δ 77.16 [$^{13}C$], MeOD δ 3.31 [$^{1}H$] and δ 49.00 [$^{13}C$]. $^{1}H$ NMR data is reported as chemical shift, multiplicity (s = singlet, d = doublet, t = triplet, q = quartet, dd = doublet of doublets, dt = doublet of triplets, tt = triplet of triplets, ddd = doublet of doublet of doublets, ddt = doublet of doublet of triplets, m = multiplet, br = broad), coupling constant (*J* Hz) and assignment where possible.

High resolution ESI+ mass spectra were measured on a Bruker–Daltonics Apex Ultra 7.0T Fourier transform mass spectrometer (FTICR). Low resolution ESI mass spectra were obtained on a Shimadzu 2020 ESI mass spectrometer operating in positive ion mode. Infrared (IR) absorption spectra were recorded on a Bruker ALPHA Spectrometer with Attenuated Total Reflection (ATR) capability. Compounds were deposited as films on the ATR plate *via* a $CH_2Cl_2$ solution. Optical rotations were recorded at ambient temperature (293K) on a Perkin–Elmer 341 polarimeter at 589 nm (sodium D line) with a cell path length of 1 dm. Melting points were determined with a SRS Optimelt melting point apparatus and are uncorrected.

Preparative reverse-phase HPLC was performed using a Waters 600 Multisolvent Delivery System and pump with Waters 486 Tuneable absorbance detector operating at 210-300 nm. Compounds were purified using an XBridge BEH $C_{18}$ 5μm (30 x 150 mm) column operating at total flow rates of 42.0 - 50.0 mL/min. A mobile phase of 0.1% TFA in water (Solvent A) and 0.1% TFA in MeCN (Solvent B) was used in all cases using gradients as reported.

Analytical UPLC-MS was performed on a Shimadzu UPLC-MS instrument consisting of a LC-M20A pump and a SPD-M30A diode array detector coupled to a Shimadzu 2020 mass spectrometer (ESI) operating in positive mode. Separations on the UPLC-MS system were performed using a Waters Acquity UPLC BEH $C_{18}$

1.7 µm (2.1 x 50 mm) column at a total flow rate of 0.60 mL/min. Separations were performed using a mobile phase of 0.1% formic acid in water (Solvent A) and 0.1% formic acid in MeCN (Solvent B).

Analytical chiral HPLC was performed using a RegisPack chiral column (250 x 4.6 mm, 5 m) at a total flow rate of 1.50 mL/min. Separations were performed using a mobile phase of n-hexane and 2-propanol.

### Section S2: Synthetic scheme for the preparation of gallinamide A analogues 24-33

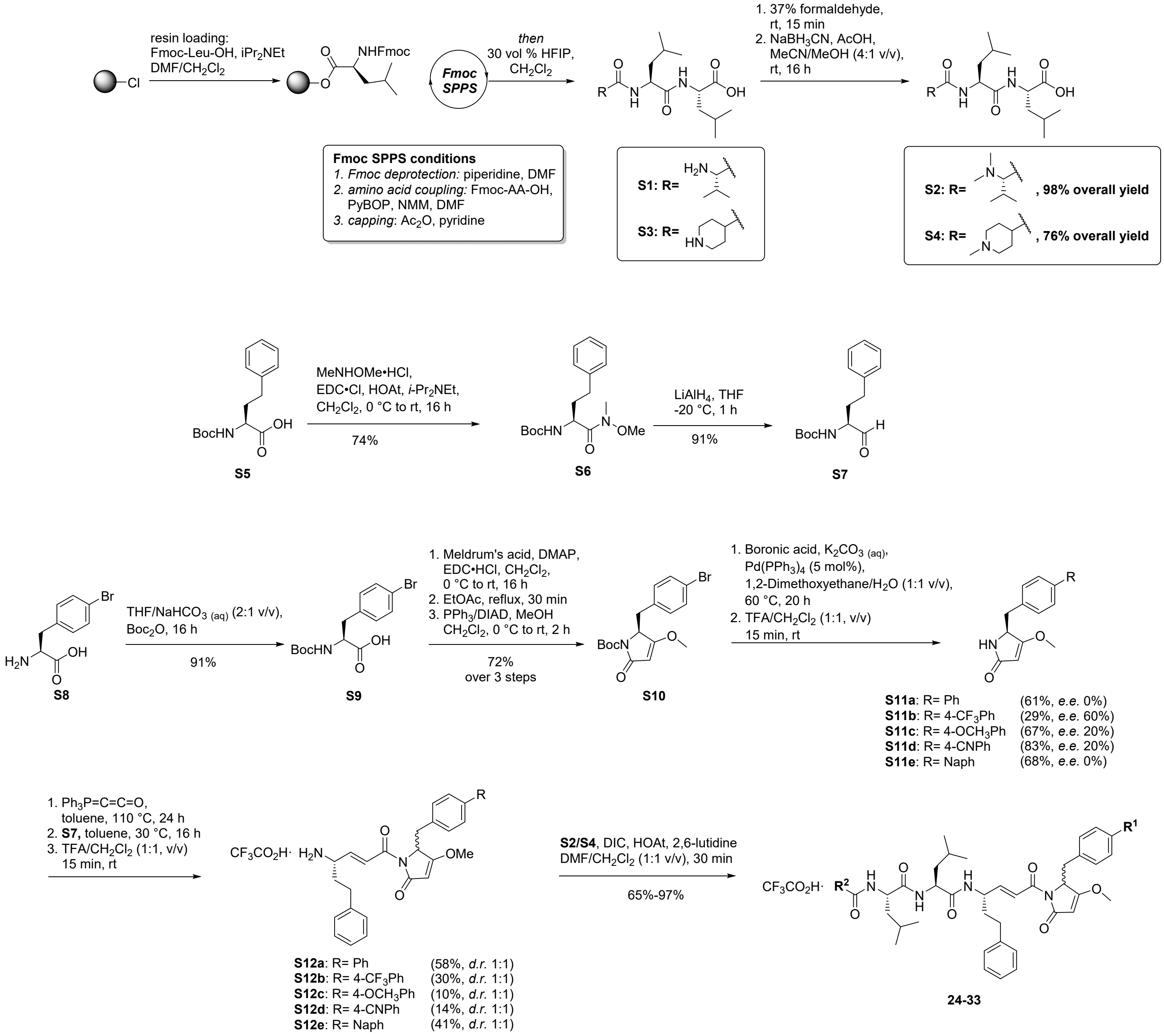


### Section S3: General procedures for chemical synthesis of gallinamide A analogues

The procedures below outline the synthesis of gallinamide A analogues **24-33**. The synthesis of analogues **2-10** and **11-23** was carried out as previously reported by Boudreau *et al (37)* and Stoye *et al,* respectively *(39).*

**General Procedure 1: Resin Loading**

2-Chlorotrityl chloride (2-CTC) resin (manufacturer's resin loading: 1.6 mmol/g, 500-600 mg) was swelled in dry $CH_2Cl_2$ (5 mL) for 30 min. The resin was subsequently shaken with a solution of 10% *i*-$Pr_2NEt$ in $CH_2Cl_2$ (5 mL) for 30 min and then washed with $CH_2Cl_2$ (5x5 mL). A solution of Fmoc-protected amino acid (1 equiv. based on resin loading) and *i*-$Pr_2NEt$ (2 equiv. based on resin loading) in $CH_2Cl_2$ (5 mL) was prepared and the resin was shaken with the solution for 16 h. The resin was subsequently washed with $CH_2Cl_2$ (5x5 mL), DMF (5x5 mL) and $CH_2Cl_2$ (5x5 mL) before treating with a capping cocktail of $CH_2Cl_2$/MeOH/ *i*-$Pr_2NEt$ (5 mL, 17:2:1 v/v/v) for 1 h. The resin was subsequently washed with DMF (5x5 mL), $CH_2Cl_2$ (5x5 mL) and DMF (5x5 mL). The efficiency of amino acid loading was determined by UV spectrophotometric analysis of the filtrate after Fmoc deprotection (20% piperidine in DMF, 2x5 min) at $\lambda$ = 301 nm ($\varepsilon$ = 7800 $M^{-1}$ $cm^{-1}$).

**General Procedure 2: Fmoc-strategy Iterative Peptide Coupling**

Fmoc-deprotection was achieved by treating the resin with 20vol.% piperidine in DMF (2x5 mL) for 5 min. The resin was subsequently washed with DMF (5x5 mL), $CH_2Cl_2$ (5x5 mL) and DMF (5x5 mL). A solution of Fmoc-protected amino acid (4 equiv. relative to resin loading), PyBOP (4 equiv. relative to resin loading) and NMM (8 equiv. relative to resin loading) in DMF (final concentraion >0.1 M) was pre-activated and the resin was treated with the solution for 1 h. The resin was subsequently washed with DMF (5x5 mL), $CH_2Cl_2$ (5x5 mL) and DMF (5x5 mL) before capping any uncoupled sequences by treating the resin with a solution of 10vol.% acetic anhydride in pyridine (2x5 mL). The resin was again washed with DMF (5x5 mL), $CH_2Cl_2$ (5x5 mL) and DMF (5x5 mL). The above process was repeated iteratively until the desired peptide sequence was assembled on resin.

**General Procedure 3: *N*-Methylation of Tripeptide**

To a solution of tripeptide in MeCN/MeOH (10 mL, 4:1 v/v) was added a solution of 37% aqueous formaldehyde (10 equiv.). The resulting mixture was stirred for 15 min or until the solution became clear. $NaBH_3CN$ (3 equiv.) was then added to the solution followed by acetic acid (0.2 mL). The reaction mixture was stirred for 16 h before the mixture was concentrated *in vacuo* and purified by preparative reverse phase HPLC.

**General Procedure 4: Suzuki Cross-coupling Reaction (0.52 mmol scale)**

A solution of anhydrous $K_2CO_3$ (237 mg, 1.72 mmol, 3.3 equiv.) in $H_2O$ (2 mL) was degassed for 15 min before Boc-*py*Phe(Br)-OH **S10** (199 mg, 0.52 mmol) and the boronic acid (0.78 mmol, 1.5 equiv.) was added, followed by addition of 1,2-dimethoxyethane (2 mL). The mixture was stirred for 10 min before $Pd(PPh_3)_4$ (30 mg, 26 μmol, 5 mol %) catalyst was added. The reaction mixture was subsequently warmed to 60 °C and stirred for 20 h. Upon completion, the mixture was diluted with $H_2O$ (4 mL) then extracted with $CH_2Cl_2$ (4x4 mL). The organic layers were collected and dried with anhydrous $MgSO_4$. The reaction was filtered through a pad of celite before the filtrate was concentrated *in vacuo* and the crude residue was used in further steps without purification.

**General Procedure 5: Boc-deprotection**

Boc-protected compound was dissolved in $CH_2Cl_2$/TFA (1:1 v/v, final concentration: 20 mM) and stirred for 15 min. Upon completion, the mixture was concentrated *in vacuo* and then purified.

**General Procedure 6: Formation of α,β-Unsaturated Pyrrolinone S12a-e with (triphenylphosphoranylidene)ketene (688 μmol scale)**

To a solution of pyrrolinone **S11a-e** (688 μmol) in toluene (1 mL) was added (triphenylphosphoranylidene)ketene (311 mg, 1.03 mmol, 1.5 equiv.). The reaction mixture was stirred at 110 °C for 24 h to afford a crude mixture of pyrrolinone-ylide. Immediately after completion, a solution of Boc-homophenylalaninal **S7** (435 mg, 1.65 mmol, 2.4 equiv.) in toluene (4 mL) was added to the mixture *in situ*. The reaction mixture was stirred for another 16 h at 30 °C. Upon completion, the crude mixture was concentrated *in vacuo* and purified by flash column chromatography. The major *E*-isomer was separated from the minor *Z*-isomer by RP-HPLC.

**General Procedure 7: Transformation of $CF_3CO_2H$ salt to HCl salt**

A $CF_3CO_2H$ salt compound was dissolved in 1 M HCl (5 mL) then slowly concentrated under a stream of nitrogen gas. This process was repeated 2 times before the compound was redissolved in MeCN/$H_2O$ and lyophilized to afford the HCl salt as a white fluffy solid.

**General Procedure 8: Formation of Gallinamide A Analogues 24-33 *via* Peptide Coupling Reaction (40.0 μmol scale)**

Tripeptide **S2** or **S4** (60.0 μmol, 1.5 equiv.), α,β-unsaturated pyrrolinone **S12a-e** (40.0 μmol) and HOAt (16.3 mg, 120 μmol, 3 equiv.) were dissolved in $CH_2Cl_2$/DMF (600 μL, 1:1 v/v). To the solution was added DIC (9.3 μL, 60.0 μmol, 1.5 equiv.) and 2,6-lutidine (11.6 μL,100 μmol, 2.5 equiv.). The reaction mixture was stirred at rt for 30 min to 1 h. Upon completion (as judged by UPLC-MS), the crude mixture was purified immediately by preparative RP-HPLC and then lyophilized.

### Section S4: Synthesis of tripeptide fragments S2 and S4

*L*-valyl-*L*-leucyl-*L*-leucine **(S1)**

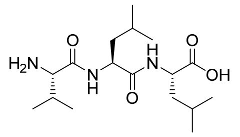


Fmoc-leucine (283 mg, 800 μmol, 1 equiv.) was loaded onto 2-CTC resin (500 mg, manufacturer's resin loading: 1.6 mmol/g) according to general procedure 1 (final resin loading: 0.79 mmol/g). Subsequently, Fmoc-leucine and Fmoc-valine were coupled according to general procedure 2. After the final coupling, the tripeptide was Fmoc deprotected by treating the resin with 20vol.% piperidine in DMF (2x5 mL) for 5 min, then washed with DMF (5x5 mL) and $CH_2Cl_2$ (10x5 mL). The tripeptide was then cleaved off from the resin by treating the resin with a solution of 30vol.% hexafluoro-2-propanol (HFIP) in $CH_2Cl_2$ (2x5 mL) then washed with $CH_2Cl_2$ (5x5 mL). The combined cleavage and washing solutions were concentrated *in vacuo* to afford the *titl*e compound **S1** as a white solid (179 mg) which was used without purification.

*N*,*N*-Dimethyl- *L*-valyl-*L*-leucyl-*L*-leucine. $CF_3CO_2H$ **(S2)**

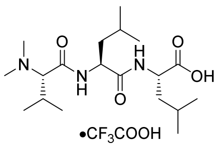


A solution of tripeptide **S1** (179 mg) was *N*-methylated according to general procedure 3. The crude product was purified by preparative RP-HPLC (gradient: 0%-40% B over 20 min, retention time: 7.5 min) and lyophilized to afford the *title* compound **S2** as a white fluffy solid (188 mg, 98% overall yield). $[\alpha]^{20}_D$ = -30° (c = 0.1, MeCN); **IR** (thin film) $\nu_{max}$ = 1645, 1550, 1467, 1369, 1190 $cm^{-1}$; **$^1$H NMR** (400 MHz, $CD_3CN$) δ 8.25 (d, 1H, *J* 8.5 Hz, NH), 7.34 (d, 1H, *J* 7.5 Hz, NH), 4.64-4.54 (m, 1H, Leu α-H), 4.36–4.25 (m, 1H, Leu α-H), 3.65 (d, 1H, *J* 7.9 Hz, Val α-H), 2.85 (s, 6H, 2 x $NCH_3$), 2.37-2.22 (m, 1H, Val CH), 1.75-1.54 (m, 6H, 2 x Leu CH, $CH_2$), 1.07 (d, 3H, *J* 6.7 Hz, $CH_3$), 0.96-0.90 (m, 12H, 4 x $CH_3$), 0.86 (d, 3H, *J* 6.7 Hz, $CH_3$); **$^{13}$C NMR** (100 MHz, $CD_3CN$) δ 174.3, 173.1, 166.5, 118.3, 73.2, 52.7, 52.2, 42.1, 40.6, 28.4, 25.6, 25.5, 23.2, 23.1, 22.2, 21.6, 19.4, 18.2. **HRMS** (+ESI) *m/z* Calc. for $C_{19}H_{37}N_3O_4$ $[M + H]^+$: 372.2862, found 372.2861.

(piperidine-4-carbonyl)- *L*-leucyl-*L*-leucine **(S3)**

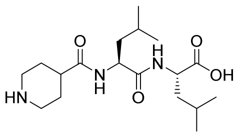

Fmoc-leucine (353 mg, 1.00 mmol, 1 equiv.) was loaded onto 2-CTC resin (600 mg, manufacturer's resin loading: 1.6 mmol/g) according to general procedure 1 (final resin loading: 0.41 mmol/g). Subsequently, Fmoc-leucine and Fmoc-isonipecotic acid were coupled according to general procedure 2. After the final coupling, the tripeptide was Fmoc deprotected by treating the resin with 20vol.% piperidine in DMF (2x5 mL) for 5 min, then washed with DMF (5x5 mL) and $CH_2Cl_2$ (10x5 mL). The tripeptide was then cleaved off from the resin by treating the resin with a solution of 30vol.% HFIP in $CH_2Cl_2$ (2x5 mL) then washed with $CH_2Cl_2$ (5x5 mL). The combined cleavage and washing solutions were concentrated *in vacuo* to afford the *titl*e compound **S3** as a white solid (89 mg) which was used without purification.

(1-methylpiperidine-4-carbonyl)- *L*-leucyl-*L*-leucine. $CF_3CO_2H$ **(S4)**

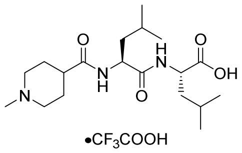


A solution of tripeptide **S3** (89 mg) was *N*-methylated according to general procedure 3. The crude product was purified by preparative RP-HPLC (gradient: 0%-40% B over 20 min, retention time: 8.2 min) and lyophilized to afford the *title* compound **S4** as a white fluffy solid (90 mg, 76% overall yield). $[\alpha]^{20}_D$ = -35° (c = 0.1, MeCN); **IR** (thin film) $\nu_{max}$ = 1664, 1537, 1194, 722 $cm^{-1}$; **$^1$H NMR** (500 MHz, $CD_3CN$) δ 7.27 (d, 1H, *J* 7.7 Hz, NH), 7.20 (d, 1H, *J* 8.1 Hz, NH), 4.45-4.22 (m, 2H, 2 x CH), 3.48 (d, 1H, *J* 13.0 Hz, CH), 3.29 (br s, 1H, alkyl-H), 2.88-2.83 (m, 2H, alkyl-H), 2.75 (s, 3H, $NCH_3$), 2.52–2.44(m, 1H), 2.13-1.95 (m, 4H, alkyl-H), 1.72-1.45 (m, 6H, alkyl-H), 0.92-0.87 (4 x d, 12H, 4 x $CH_3$); **$^{13}$C NMR** (125 MHz, $CD_3CN$) δ 174.8, 174.5,

173.7, 54.5, 54.5, 53.2, 51.8, 44.1, 41.5, 41.1, 40.2, 27.0, 27.0, 25.6, 25.6, 23.3, 23.2, 22.0, 21.8; **HRMS** (+ESI) *m/z* Calc. for $C_{19}H_{35}N_3O_4$ $[M + Na]^+$: 392.2520, found 392.2518.

### Section S5: Synthesis of pyrrolinone building blocks S12a-e

*tert*-Butyl (*S*)-(1-(methoxy(methyl)amino)-1-oxo-4-phenylbutan-2-yl)carbamate (Weinreb amide) **(S6)**

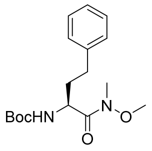


A solution of Boc-*L*-homophenylalanine **S5** (7.50 g, 26.8 mmol) in $CH_2Cl_2$ (100 mL) was cooled to 0 °C before *N*,*O*-dimethylhydroxylamine.HCl (1.17g, 12.0 mmol, 1.2 equiv.) was added, followed by 1-ethyl-3-(3-dimethylaminopropyl)carbodiimide.HCl (EDC.HCl) (2.30 g, 12.0 mmol, 1.2 equiv.), HOAt (816 mg, 6.00 mmol, 0.6 equiv.) and *i*-$Pr_2NEt$ (2.6 mL, 15.0 mmol, 1.5 equiv.). The reaction mixture was warmed to rt and stirred for 16 h. Upon completion, the mixture was diluted with $CH_2Cl_2$ (100 mL) and then washed with 1 M HCl (2x100 mL), followed by sat. aq. $NaHCO_3$ solution (2x100 mL). The organic layer was collected and dried with $MgSO_4$ before it was concentrated *in vacuo* to afford the *title* compound **S6** as a brown oil (6.39g, 74%). **$^1$H NMR** (300 MHz, $CDCl_3$) δ 7.20-7.35 (m, 5H,Ar-*H*), 5.33 (s, 1H, NH), 4.73 (br s, 1H, α-H), 3.67 (s, 3H, $OCH_3$), 3.21 (s, 3H, $CH_3$), 2.66-2.85 (m, 2H, $CH_2$), 1.82-2.26 (m, 2H, $CH_2$), 1.50 (s, 9H, 3 x $CH_3$); **$^{13}$C NMR** (75 MHz, $CDCl_3$) δ 173.2, 155.7, 141.3, 128.6, 128.5, 126.1, 79.7, 74.1, 61.6, 34.7, 32.2, 31.8, 28.5.

*tert*-Butyl (*S*)-(1-oxo-4-phenylbutan-2-yl)carbamate **(S7)**

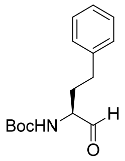


A solution of Weinreb amide **S6** (445 mg, 1.38 mmol) in THF (6 mL) was cooled to -20 °C before $LiAlH_4$ (1.73 mL, 1.73 mmol, 1.25 equiv.) was added dropwise. The reaction mixture was maintained at -20 °C and stirred for 1 h. Upon completion, the mixture was slowly quenched (dropwise to prevent excessive foaming) with 1 M HCl (140 µL). Afterwards, the mixture was diluted with EtOAc (20 mL) and washed with 1 M HCl (20 mL), then brine (20 mL). The organic layer was collected and dried with $MgSO_4$ before it was concentrated *in vacuo* to afford the *title* compound **S7** as a milky oil (329 mg, 91%). The product was used immediately in the next step without purification to minimize any decomposition of the aldehyde.

(*S*)-3-(4-bromophenyl)-2-((*tert*-butoxycarbonyl)amino)propanoic acid **(S9)**

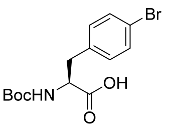

A solution of 4-bromo-L-phenylalanine **S8** (10.0 g, 41.0 mmol) and Boc-anhydride (17.9 g, 82.0 mmol, 2 equiv.) in sat. aq. $NaHCO_3$ solution/THF (250 mL, 2:3 v/v) was stirred for 16 h. Afterwards, the reaction mixture was diluted with water (100 mL), then washed with diethyl ether (4x200 mL). The aqueous layer was collected and acidified to pH 3 with 1 M HCl. The resulting suspension was extracted with EtOAc (4x200 mL), then dried with anhydrous $MgSO_4$ before it was concentrated *in vacuo* to afford the *title* compound **S9** as a white solid (12.9 g, 91%). **$[\alpha]^{20}_D$** = +21.9° (c = 1.0, EtOH); **$^1$H NMR** (300 MHz, DMSO-$d_6$) δ 7.47 (d, 2H, *J* 7.8 Hz, Ar-*H*), 7.21 (d, 2H, *J* 7.8 Hz, Ar-*H*), 3.99-4.12 (m, 1H, α-H), 2.99 (dd, 1H, *J* 4.3, 14.0 Hz, C*H*H), 2.79 (dd, 1H, *J* 10.8, 14.0 Hz, CH*H*), 1.31 (s, 9H, 3 x $CH_3$); **$^{13}$C NMR** (75 MHz, DMSO-$d_6$) δ 173.4, 155.4, 137.5, 131.4, 131.0, 119.5, 78.1, 54.9, 35.8, 28.1.

*tert*-Butyl (*S*)-2-(4-bromobenzyl)-3-methoxy-5-oxo-2,5-dihydro-1*H*-pyrrole-1-carboxylate **(S10)**

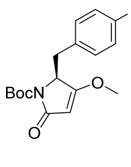


A solution of Meldrum's acid (1.84 g, 12.8 mmol, 1.1 equiv.) and DMAP (1.84 g, 15.1 mmol, 1.3 equiv.) in $CH_2Cl_2$ (100 mL) was cooled to 0 °C. Compound **S9** (4.00 g, 11.6 mmol) was added, followed by EDC.HCl (2.66 g, 13.9 mmol, 1.2 equiv.). The reaction mixture was warmed to rt and stirred for 16 h. The reaction mixture was diluted with EtOAc (500 mL) and then washed with brine (2x200 mL), 5% aq. citric acid (3x300 mL), then brine (1x200 mL). The organic layer was collected, dried with anhydrous $MgSO_4$ and heated at reflux for 30 min. Upon completion, the reaction mixture was cooled and concentrated *in vacuo.* To this crude mixture in $CH_2Cl_2$ (60 mL) was added $PPh_3$ (3.37 g, 12.9 mmol, 1.3 equiv.) and MeOH (0.52 mL, 12.9 mmol, 1.3 equiv.) before cooling to 0 °C. DIAD (2.53 mL, 12.9 mmol, 1.3 equiv.) was added dropwise to the cooled mixture, then the reaction mixture was warmed to rt and stirred for 2 h before concentrating *in vacuo*. The crude product was purified by flash column chromatography (eluent: 50% EtOAc in hexane) to afford the *title* compound **S10** as a white solid (3.50 g, 72% over 3 steps) with traces of inseparable DIAD. **$[\alpha]^{20}_D$** = +149° (c = 0.1, $CHCl_3$); **m.p.** = 88-90 °C; **IR** (thin film) $\nu_{max}$ = 2923, 2853, 1776, 1736, 1633, 1454, 1370, 1322, 1244, 1153, 1095, 1012, 983 $cm^{-1}$; **$^1$H NMR** (400 MHz, $CDCl_3$) δ 7.34 (m, 2H, Ar-*H*), 6.86 (m, 2H, Ar-*H*), 4.84 (s, 1H, CH), 4.64 (dd, 1H, *J* 3.1, 5.4 Hz, α-H), 3.77 (s, 3H, $OCH_3$), 3.40 (dd, 1H, *J* 5.2, 14.0 Hz, C*H*H), 3.06 (dd, 1H, *J* 3.2, 13.9 Hz, CH*H*), 1.58 (s, 9H, 3 x $CH_3$); **$^{13}$C NMR** (100 MHz, $CDCl_3$) δ 176.0, 168.5, 149.7, 133.4, 131.5, 131.3, 121.2, 95.5, 82.9, 60.0, 58.4, 34.8, 28.3; **HRMS** (+ESI) *m/z* Calc. for $C_{17}H_{20}BrNO_4$ $[M + Na]^+$: 404.0468, found 404.0467.

*(S)*-5-([1,1'-biphenyl]-4-ylmethyl)-4-methoxy-1,5-dihydro-2*H*-pyrrol-2-one **(S11a)**

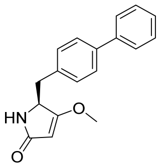


Compound **S10** (1.31 mmol) was cross-coupled with phenyl boronic acid (1.5 equiv.) according to general procedure 4, then Boc-deprotected according to general procedure 5. The crude product was purified by flash column chromatography (eluent: 80% EtOAc in hexane) to afford the *title* compound **S11a** as a white foam (224 mg, 61% over 2 steps). **[α]$^{20}_{D}$** = -0.5° (c = 0.1, $CHCl_3$); ***e.e.*:** 0% (Determined by chiral HPLC, gradient: 5%-20% isopropanol in hexane over 20 min.); **IR** (thin film) $v_{max}$ = 1683, 1623, 1487, 1363, 1230, 990, 807, 757, 698 cm$^{-1}$; **$^1$H NMR** (500 MHz, $CDCl_3$) δ 7.52-7.58 (m, 4H, Ar-*H*), 7.41-7.45 (m, 2H, Ar-*H*), 7.32-7.36 (m, 1H, Ar-*H*), 7.25-7.27 (m, 2H, Ar-*H*), 5.49 (s, 1H, NH), 4.84 (d, 1H, *J* 1.2 Hz, CH), 4.25 (dd, 1H, *J* 3.4, 9.5 Hz, α-H), 3.84 (s, 3H, $OCH_3$), 3.23 (dd, 1H, *J* 3.6, 13.6 Hz, C*H*H), 2.66 (dd, 1H, *J* 9.6, 14.3 Hz, CH*H*); **$^{13}$C NMR** (125 MHz, $CDCl_3$) δ 177.6, 173.7, 140.8, 140.2, 135.8, 129.7, 128.9, 127.6, 127.5, 127.2, 94.2, 58.8, 58.5, 38.5; **HRMS** (+ESI) *m/z* Calc. for $C_{18}H_{17}NO_2$ [M + Na]$^+$: 302.1152, found 302.1154.

(*S*)-4-methoxy-5-((4'-(trifluoromethyl)-[1,1'-biphenyl]-4-yl)methyl)-1,5-dihydro-2*H*-pyrrol-2-one **(S11b)**

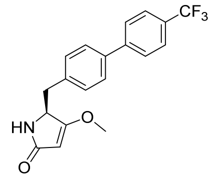


Compound **S10** (0.52 mmol) was cross-coupled with 4-trifluoro-phenyl boronic acid (1.5 equiv.) according to general procedure 4, then Boc-deprotected according to general procedure 5. The crude product was purified by flash column chromatography (eluent: 80% EtOAc in hexane) to afford the *title* compound **S11b** as a yellow oil (51.5 mg, 29% over 2 steps). **[α]$^{20}_{D}$** = -50° (c = 0.1, $CHCl_3$); ***e.e.*:** 60% (Determined by chiral HPLC, gradient: 5%-20% isopropanol in hexane over 20 min.); **IR** (thin film) $v_{max}$ = 2923, 2852, 1686, 1622, 1458, 1369, 1326, 1232, 1166, 1121, 1071, 814 cm$^{-1}$; **$^1$H NMR** (500 MHz, $CDCl_3$) δ 7.66-7.70 (m, 4H, Ar-*H*), 7.54 (d, 2H, *J* 8.4 Hz, Ar-*H*), 7.29 (d, 2H, *J* 8.2 Hz, Ar-*H*), 5.72 (s, 1H, NH), 5.01 (s, 1H, CH), 4.27 (dd, 1H, *J* 3.4, 9.1 Hz, α-H), 3.85 (s, 3H, $OCH_3$), 3.24 (dd, 1H, *J* 3.4, 13.4 Hz, C*H*H), 2.72 (dd, 1H, *J* 8.7, 14.8 Hz, CH*H*); **$^{13}$C NMR** (125 MHz, $CDCl_3$) δ 177.6, 174.0, 144.3, 138.7, 136.7, 130.0, 129.7, 129.4, 127.7, 127.4, 125.9 (q, *J* 3.7 Hz, $CF_3$), 94.2, 58.7, 58.6, 38.3; **HRMS** (+ESI) *m/z* Calc. for $C_{19}H_{16}F_3NO_2$ [M + Na]$^+$: 370.1025, found 370.1025.

(*S*)-4-methoxy-5-((4'-methoxy-[1,1'-biphenyl]-4-yl)methyl)-1,5-dihydro-2*H*-pyrrol-2-one **(S11c)**

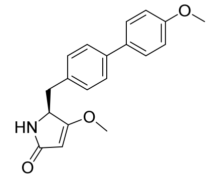


Compound **S10** (0.52 mmol) was cross-coupled with 4-methoxy-phenyl boronic acid (1.5 equiv.) according to general procedure 4, then Boc-deprotected according to general procedure 5. The crude product was purified by flash column chromatography (eluent: 80% EtOAc in hexane) to afford the *title* compound **S11c** as a colourless oil (107 mg, 67% over 2 steps). **[α]$^{20}_{D}$** = -18° (c = 0.1, $CHCl_3$); ***e.e.*:** 20% (Determined by chiral HPLC, gradient: 5%-20% isopropanol in hexane over 20 min.); **IR** (thin film) $v_{max}$ = 2923, 2854, 1683, 1621, 1500, 1459, 1367, 1247, 1178, 1127, 809 cm$^{-1}$; **$^1$H NMR** (400 MHz, $CDCl_3$) δ 7.47 (t, 4H, *J* 8.9 Hz, Ar-*H*), 7.20 (d, 1H, *J* 8.1 Hz, Ar-*H*), 6.95 (d, 1H, *J* 8.8 Hz, Ar-*H*), 5.96 (s, 1H, NH), 5.00 (s, 1H, CH), 4.21 (dd, 1H, *J* 3.1, 9.4 Hz, α-H), 3.83 (s, 3H, $OCH_3$), 3.78 (s, 3H, $OCH_3$), 3.17 (dd, 1H, *J* 3.1, 14.0 Hz, C*H*H), 2.64 (dd, 1H,

*J* 9.1, 14.4 Hz, CH*H*); **$^{13}$C NMR** (100 MHz, $CDCl_3$) δ 177.9, 159.3, 139.7, 134.9, 133.4, 129.7, 128.1, 127.1, 114.4, 94.0, 60.5, 59.0, 58.5, 55.5, 38.2; **HRMS** (+ESI) *m/z* Calc. for $C_{19}H_{19}NO_3$ $[M + Na]^+$: 332.1257, found 332.1258.

(*S*)-4'-((3-methoxy-5-oxo-2,5-dihydro-1*H*-pyrrol-2-yl)methyl)-[1,1'-biphenyl]-4-carbonitrile **(S11d)**

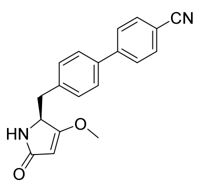


Compound **S10** (0.52 mmol) was cross-coupled with 4-cyano-phenyl boronic acid (1.5 equiv.) according to general procedure 4, then Boc-deprotected according to general procedure 5. The crude product was purified by flash column chromatography (eluent: 80% EtOAc in hexane) to afford the *title* compound **S11d** as a white foam (130 mg, 83% over 2 steps). **$[\alpha]^{20}_D$** = -29° (c = 0.1, $CHCl_3$); ***e.e.*:** 20% (Determined by chiral HPLC, gradient: 5%-20% isopropanol in hexane over 20 min.); **IR** (thin film) $\nu_{max}$ = 2923, 2852, 2226, 1681, 1621, 1494, 1454, 1363, 1230, 1176, 1116, 1079, 1027, 989, 954, 805, 748 $cm^{-1}$; **$^1$H NMR** (400 MHz, $CDCl_3$) δ 7.70 (d, 2H, *J* 8.4 Hz, Ar-*H*), 7.64 (d, 2H, *J* 8.4 Hz, Ar-*H*), 7.51 (d, 2H, *J* 8.1 Hz, Ar-*H*), 7.29 (d, 2H, *J* 8.3 Hz, Ar-*H*), 6.01 (d, 1H, *J* 13.3 Hz, NH), 4.97 (s, 1H, CH), 4.27 (dd, 1H, *J* 3.5, 8.5 Hz, α-H), 3.82 (s, 3H, $OCH_3$), 3.21 (dd, 1H, *J* 3.7, 13.7 Hz, C*H*H), 2.75 (dd, 1H, *J* 8.4, 14.0 Hz, CH*H*); **$^{13}$C NMR** (100 MHz, $CDCl_3$) δ 177.2, 173.8, 145.1, 137.9, 137.0, 137.0, 132.6, 130.0, 127.6, 127.4, 118.9, 110.9, 94.2, 58.3, 38.0; **HRMS** (+ESI) *m/z* Calc. for $C_{19}H_{16}N_2O_2$ $[M + Na]^+$: 327.1104, found 327.1106.

(*S*)-4-methoxy-5-(4-(naphthalen-2-yl)benzyl)-1,5-dihydro-2*H*-pyrrol-2-one **(S11e)**

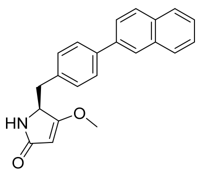


Compound **S10** (0.90 mmol) was cross-coupled with naphthyl boronic acid (1.5 equiv.) according to general procedure 4, then Boc-deprotected according to general procedure 5. The crude product was purified by flash column chromatography (eluent: 80% EtOAc in hexane) to afford the *title* compound **S11e** as a white foam (202 mg, 68% over 2 steps). **$[\alpha]^{20}_D$** = -16° (c = 0.1, $CHCl_3$); ***e.e.*:** 10% (Determined by chiral HPLC, gradient: 5%-20% isopropanol in hexane over 20 min.); **IR** (thin film) $\nu_{max}$ = 2924, 2853, 1683, 1622, 1502, 1459, 1363, 1231, 809 $cm^{-1}$; **$^1$H NMR** (400 MHz, $CDCl_3$) δ 8.02 (s, 1H, Ar-*H*), 7.85-7.92 (m, 3H, Ar-*H*), 7.72 (dd, 1H, *J* 1.7, 8.5 Hz, Ar-*H*), 7.67 (d, 2H, *J* 8.2 Hz, Ar-*H*), 7.46-7.53 (m, 2H, Ar-*H*), 7.31 (d, 2H, *J* 8.0 Hz, Ar-*H*), 5.72 (s, 1H, NH), 5.02 (s, 1H, CH), 4.27 (dd, 1H, *J* 3.3, 9.2 Hz, α-H), 3.84 (s, 3H, $OCH_3$), 3.25 (dd, 1H, *J* 3.6, 13.7 Hz, C*H*H), 2.71 (dd, 1H, *J* 9.2, 13.8 Hz, CH*H*); **$^{13}$C NMR** (100 MHz, $CDCl_3$) δ 177.5, 173.8, 140.0, 138.1, 135.8, 133.8, 132.7, 129.8, 128.6, 128.3, 127.8, 126.4, 126.1, 125.8, 125.5, 94.2, 58.7, 58.5, 38.4; **HRMS** (+ESI) *m/z* Calc. for $C_{22}H_{19}NO_2$ $[M + Na]^+$: 352.1308, found 352.1308.

5-([1,1'-biphenyl]-4-ylmethyl)-1-((*S*,*E*)-4-amino-6-phenylhex-2-enoyl)-4-methoxy-1,5-dihydro-2*H*-pyrrol-2-one **(S12a)**

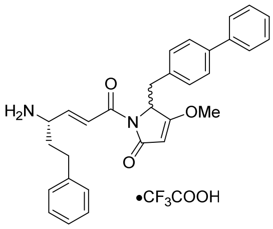


Compound **S11a** (688 µmol) was reacted according to general procedure 6. The crude product was purified by flash column chromatography (eluent: 40% EtOAc in hexane) to afford the Boc-protected compound as a yellow oil (*E/Z* 17:3 as determined by UPLC-MS). The *E*-isomer was isolated by preparative RP-HPLC (gradient: 50%-80% B over 15 min, retention time: 9.4-10.7 min (*E*-isomer), 11.0-11.7 min (*Z*-isomer)). After concentration *in vacuo*, the pure *E*-isomer was subjected to Boc-deprotection according to general procedure 5 and purified by preparative RP-HPLC (gradient: 30%-40% B over 20 min, retention time: 19.8-23.0 min). The product was lyophilized to afford the *title* compound **S12a** as a white fluffy solid (231 mg, 58% over 3 steps, *d.r.* 1:1 as determined by **$^1$H NMR**). **$[\alpha]^{20}_D$** = +24° (c = 0.1, MeOH); **IR** (thin film) $\nu_{max}$ = 2939, 1725, 1677, 1625, 1488, 1453, 1382, 1351, 1307, 1250, 1201, 1131, 969, 838, 801, 753, 722, 698 cm$^{-1}$; **$^1$H NMR** (500 MHz, $CDCl_3$, 1:1 mixture of diastereoisomers) δ 8.55 (br s, 2H, $NH_2$), 7.50-7.53 (m, 3H, 2 x Ar-*H*, CH=C*H*C=O), 7.35-7.44 (m, 4H, Ar-*H*), 7.29 (t, 1H, *J* 7.31 Hz, Ar-*H*), 7.20-7.23 (m, 2H, Ar-*H*), 7.13-7.16 (m, 3H, Ar-*H*), 7.03 (dd, 1H, *J* 4.0, 14.7 Hz, C*H*=CHC=O), 6.98 (dd, 2H, *J* 5.7, 7.9 Hz, Ar-*H*), 4.81-4.84 (m, 2H, 2 x CH), 3.96 (s, 1H, CH), 3.77 (d, 3H, *J* 4.7 Hz, $OCH_3$), 3.48 (ddd, 1H, *J* 4.8, 13.8, 28.5 Hz, C*H*H), 3.12 (dd, 1H, *J* 2.3, 14.4 Hz, CH*H*), 2.61-2.76 (m, 2H, $CH_2$), 2.09-2.23 (m, 2H, $CH_2$); **$^{13}$C NMR** (125 MHz, $CDCl_3$, 1:1 mixture of diastereoisomers) δ 178.8, 178.8, 170.3, 164.1, 163.9, 141.4, 141.4, 140.6, 140.6, 140.0, 140.9, 139.8, 139.8, 133.3, 133.3, 130.1, 128.9, 128.8, 128.8, 128.5, 128.5, 127.4, 127.4, 127.2, 127.0, 127.0, 127.0, 126.5, 126.5, 94.9, 94.9, 60.0, 60.0, 58.7, 52.6, 52.3, 34.5, 34.4, 34.3, 34.3, 31.4, 31.2; **HRMS** (+ESI) *m/z* Calc. for $C_{30}H_{30}N_2O_3$ $[M + Na]^+$: 489.2149, found 489.2151.

1-((*S*,*E*)-4-amino-6-phenylhex-2-enoyl)-4-methoxy-5-((4'-(trifluoromethyl)-[1,1'-biphenyl]-4-yl)methyl)-1,5-dihydro-2*H*-pyrrol-2-one **(S12b)**

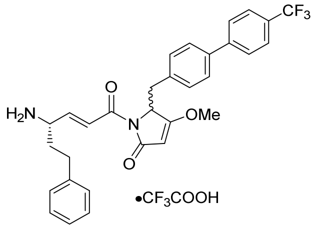


Compound **S11b** (188 µmol) was reacted according to general procedure 6. The crude product was purified by flash column chromatography (eluent: 40% EtOAc in hexane) to afford the Boc-protected compound as a yellow oil (*E/Z* 9:1 as determined by UPLC-MS). The *E*-isomer was isolated by preparative RP-HPLC (gradient: 50%-80% B over 15 min, retention time: 11.8-12.4 min (*E*-isomer), 12.7-13.1 min (*Z*-isomer)). After concentration *in vacuo*, the pure *E*-isomer was subjected to Boc-deprotection according to general procedure 5 and purified by preparative RP-HPLC (gradient: 40%-50% B over 10 min, retention time: 10.1-11.9 min). The product was lyophilized to afford the *title* compound **S12b** as a white fluffy solid (36.0 mg, 30% over 3 steps, *d.r.* 1:1 as determined by **$^1$H NMR**). **$[\alpha]^{20}_D$** = +33° (c = 0.1, MeOH); **IR** (thin film) $v_{max}$ = 2922, 1727, 1674, 1627, 1326, 1123, 1071, 970, 834 cm$^{-1}$; **$^1$H NMR** (400 MHz, MeOD, 1:1 mixture of diastereoisomers) δ 7.64-7.76 (m, 4H, 3 x Ar-*H*, CH=C*H*C=O), 7.52-7.57 (m, 3H, Ar-*H*), 7.19-7.32 (m, 5H, Ar-*H*), 7.11 (dd, 2H, *J* 1.4, 8.3 Hz, Ar-*H*), 6.98 (ddd, 1H, *J* 1.4, 8.0, 15.7 Hz, C*H*=CHC=O), 5.13 (s, 1H, CH), 5.05-5.06 (m, 1H, CH), 4.00 (d, 3H, *J* 5.6 Hz, $OCH_3$), 3.93-3.97 (m, 1H, CH), 3.62-3.69 (m, 1H, C*H*H), 3.30 (dd, 1H, *J* 2.4, 14.0 Hz, CH*H*), 2.63-2.83 (m, 2H, $CH_2$), 2.03-2.21 (m, 2H, $CH_2$); **$^{13}$C NMR** (100 MHz, MeOD, 1:1 mixture of diastereoisomers) δ 180.6, 180.5, 172.1, 172.0, 164.9, 164.8, 145.7, 145.6, 141.7, 141.6, 141.3, 141.3, 139.6, 135.9, 135.8, 131.5, 129.8, 129.7, 129.6, 129.5, 129.3, 129.0, 128.4, 128.3, 128.0, 127.6, 127.5, 126.8, 126.7, 126.7, 95.8, 95.7, 61.1, 61.1, 59.8, 59.7, 53.3, 53.1, 35.7, 35.5, 34.8, 34.7, 32.4, 32.2; **HRMS** (+ESI) *m/z* Calc. for $C_{31}H_{29}F_3N_2O_3$ [M + Na]$^+$: 557.2023, found 557.2027.

1-((*S*,*E*)-4-amino-6-phenylhex-2-enoyl)-4-methoxy-5-((4'-methoxy-[1,1'-biphenyl]-4-yl)methyl)-1,5-dihydro-2*H*-pyrrol-2-one **(S12c)**

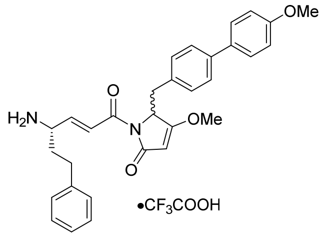


Compound **S11c** (297 µmol) was reacted according to general procedure 6. The crude product was purified by flash column chromatography (eluent: 40% EtOAc in hexane) to afford the Boc-protected compound as a yellow oil (*E/Z* 93:7 as determined by UPLC-MS). The *E*-isomer was isolated by preparative RP-HPLC (gradient: 50%-80% B over 15 min, retention time: 9.4-10.3 min (*E*-isomer), 10.6-11.3 min (*Z*-isomer)). After concentration *in vacuo*, the pure *E*-isomer was subjected to Boc-deprotection according to general procedure 5 and purified by preparative RP-HPLC (gradient: 45%-55% B over 10 min, retention time: 7.7-8.6 min). The product was lyophilized to afford the *title* compound **S12c** as a white fluffy solid (18.3 mg, 10% over 3 steps, *d.r.* 1:1 as determined by **$^1$H NMR**). **$[\alpha]^{20}_D$** = -12° (c = 0.1, MeOH); **IR** (thin film) $v_{max}$ = 2924, 1724, 1674, 1626, 1499, 1445, 1382, 1350, 1308, 1248, 1178, 1125, 972, 826 cm$^{-1}$; **$^1$H NMR** (400 MHz, DMSO-$d_6$, 1:1 mixture of diastereoisomers) δ 8.32 (br s, 2H, $NH_2$), 7.56 (d, 1H, *J* 8.2 Hz, CH=C*H*C=O), 7.40-7.49 (m, 4H, Ar-*H*), 7.21-7.31 (m, 5H, Ar-*H*), 6.89-7.00 (m, 5H, 4 x Ar-*H*, C*H*=CHC=O), 5.26 (d, 1H, *J* 3.7 Hz, CH), 5.02-5.05 (m, 1H, CH), 3.94-4.00 (m, 1H, CH), 3.91 (d, 3H, *J* 6.7 Hz, $OCH_3$), 3.77 (d, 3H, *J* 5.9 Hz, $OCH_3$), 3.44-3.55 (m, 1H, C*H*H), 3.07-3.12 (m, 1H, CH*H*), 2.55-2.74 (m, 2H, $CH_2$), 1.86-2.15 (m, 2H, $CH_2$); **$^{13}$C NMR** (100 MHz, DMSO-$d_6$, 1:1 mixture of diastereoisomers) δ 178.5, 178.5, 169.8, 169.7, 162.9, 162.9, 158.8, 158.8, 141.9, 140.5, 140.4, 138.2, 132.6, 132.5, 131.8, 131.7, 129.9, 129.9, 128.5, 128.5, 128.3, 128.2, 127.5, 127.4, 126.5, 126.4, 126.2, 126.2, 125.7, 125.6, 114.3, 114.3, 94.9, 59.2, 59.2, 59.1, 59.1, 55.1, 55.1, 51.2, 51.1, 33.9, 33.8, 33.3, 33.2, 30.7, 30.5; **HRMS** (+ESI) *m/z* Calc. for $C_{31}H_{32}N_2O_4$ [M + Na]$^+$: 519.2254, found 519.2256.

4'-((1-((*S*,*E*)-4-amino-6-phenylhex-2-enoyl)-3-methoxy-5-oxo-2,5-dihydro-1*H*-pyrrol-2-yl)methyl)-[1,1'-biphenyl]-4-carbonitrile **(S12d)**

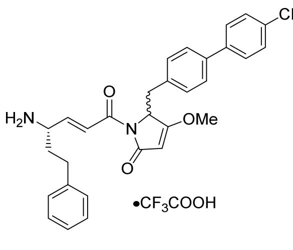

Compound **S11d** (427 µmol) was reacted according to general procedure 6. The crude product was purified by flash column chromatography (eluent: 40% EtOAc in hexane) to afford the Boc-protected compound as a yellow oil (*E/Z* 93:7 as determined by UPLC-MS). The *E*-isomer was isolated by preparative RP-HPLC (gradient: 50%-80% B over 15 min, retention time: 8.7-9.4 min (*E*-isomer), 9.5-9.7 min (*Z*-isomer)). After concentration *in vacuo*, the pure *E*-isomer was subjected to Boc-deprotection according to general procedure 5 and purified by preparative RP-HPLC (gradient: 55%-65% B over 10 min, retention time: 6.7-7.0 min). The product was lyophilized to afford the *title* compound **S12d** as a white fluffy solid (35.0 mg, 14% over 3 steps, *d.r.* 1:1 as determined by **$^1$H NMR**). **$[\alpha]^{20}_D$** = +42° (c = 0.1, MeOH); **IR** (thin film) $\nu_{max}$ = 2921, 2226, 1724, 1676, 1626, 1495, 1454, 1350, 1308, 1250, 1201, 1128, 968, 813 cm$^{-1}$; **$^1$H NMR** (400 MHz, DMSO-$d_6$, 1:1 mixture of diastereoisomers) δ 8.31 (br s, 2H, $NH_2$), 7.90 (d, 1H, *J* 8.4 Hz, Ar-*H*), 7.82-7.85 (m, 2H, Ar-*H*), 7.75 (d, 1H, *J* 7.5 Hz, Ar-*H*), 7.62 (dd, 2H, *J* 8.2, 14.6 Hz, Ar-*H*), 7.42 (dd, 1H, *J* 3.4, 15.8 Hz, CH=C*H*C=O), 7.27-7.31 (m, 2H, Ar-*H*), 7.19-7.22 (m, 3H, Ar-*H*), 7.04 (d, 2H, *J* 7.5 Hz, Ar-*H*), 6.89-6.97 (m, 1H, C*H*=CHC=O), 5.27 (d, 1H, *J* 5.0 Hz, CH), 5.05-5.08 (m, 1H, CH), 3.94 (br s, 1H, CH), 3.91 (d, 3H, *J* 7.2 Hz, $OCH_3$), 3.47-3.53 (m, 1H, C*H*H), 3.12-3.17 (m, 1H, CH*H*), 2.55-2.69 (m, 2H, $CH_2$), 1.88-2.09 (m, 2H, $CH_2$); **$^{13}$C NMR** (100 MHz, DMSO-$d_6$, 1:1 mixture of diastereoisomers) δ 178.5, 178.4, 169.8, 169.6, 162.9, 162.9, 143.9, 143.9, 141.9, 140.5, 140.4, 136.6, 136.6, 135.1, 135.0, 132.9, 132.8, 130.2, 130.2, 128.5, 128.5, 128.3, 128.2, 127.3, 127.2, 126.7, 126.6, 126.4, 126.2, 126.2, 118.8, 118.8, 110.0, 109.9, 94.9, 94.9, 59.2, 59.2, 59.0, 59.0, 51.2, 51.1, 33.9, 33.7, 33.5, 33.4, 30.6, 30.5; **HRMS** (+ESI) *m/z* Calc. for $C_{31}H_{29}N_3O_3$ [M + Na]$^+$: 514.2101, found 514.2104.

1-((*S*,*E*)-4-amino-6-phenylhex-2-enoyl)-4-methoxy-5-(4-(naphthalen-2-yl)benzyl)-1,5-dihydro-2*H*-pyrrol-2-one **(S12e)**

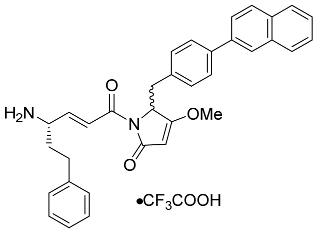

Compound **S11e** (606 µmol) was reacted according to general procedure 6. The crude product was purified by flash column chromatography (eluent: 40% EtOAc in hexane) to afford the Boc-protected compound as a yellow oil (*E/Z* 91:9 as determined by UPLC-MS). The *E*-isomer was isolated by preparative RP-HPLC (gradient: 60%-90% B over 15 min, retention time: 8.1-9.1 min (*E*-isomer), 9.3-9.8 min (*Z*-isomer)). After concentration *in vacuo*, the pure *E*-isomer was subjected to Boc-deprotection according to general procedure

5 and purified by preparative RP-HPLC (gradient: 40%-50% B over 10 min, retention time: 10.2-12.6 min). The product was lyophilized to afford the *title* compound **S12e** as a white fluffy solid (157 mg, 41% over 3 steps, *d.r.* 1:1 as determined by **$^1$H NMR**). **$[\alpha]^{20}_D$** = +32° (c = 0.1, MeCN); **IR** (thin film) $v_{max}$ = 2926, 1725, 1675, 1626, 1501, 1455, 1350, 1307, 1249, 1201, 1129, 970, 811 $cm^{-1}$; **$^1$H NMR** (400 MHz, DMSO-$d_6$, 1:1 mixture of diastereoisomers) δ 8.31 (br s, 2H, $NH_2$), 8.14 (dd, 1H, *J* 1.1, 31.7 Hz), 7.88-8.00 (m, 3H, Ar-*H*), 7.66-7.82 (m, 3H, Ar-*H*), 7.43-7.57 (m, 3H, 2 x Ar-*H*, CH=C*H*C=O), 7.18-7.32 (m, 5H, Ar-*H*), 7.05 (d, 2H, *J* 7.8 Hz, Ar-*H*), 6.91-7.00 (m, 1H, C*H*=CHC=O), 5.29 (d, 1H, *J* 4.2 Hz, CH), 5.06-5.10 (m, 1H, CH), 3.97 (br s, 1H, CH), 3.94 (d, 3H, *J* 6.3 Hz, $OCH_3$), 3.50-3.56 (m, 1H, C*H*H), 3.13-3.18 (m, 1H, CH*H*), 2.57-2.73 (m, 2H, $CH_2$), 1.90-2.12 (m, 2H, $CH_2$); **$^{13}$C NMR** (100 MHz, DMSO-$d_6$, 1:1 mixture of diastereoisomers) δ 178.5, 178.5, 169.8, 169.7, 162.9, 162.9, 158.1 (q, *J* 32.9 Hz, $CF_3CO_2H$), 141.9, 141.8, 140.5, 140.4, 138.3, 136.8, 136.7, 133.6, 133.5, 133.3, 133.2, 132.2, 132.2, 130.1, 130.1, 128.5, 128.5, 128.4, 128.2, 128.2, 128.1, 128.1, 127.5, 127.4, 126.5, 126.4, 126.4, 126.4, 126.2, 126.2, 126.1, 126.1, 125.0, 124.9, 124.8, 124.7, 94.9, 59.2, 59.2, 59.1, 59.1, 51.2, 51.1, 33.9, 33.8, 33.4, 33.3, 30.7, 30.5; **HRMS** (+ESI) *m/z* Calc. for $C_{34}H_32N_2O_3$ [M + H]$^+$: 517.2486, found 517.2488.

### Section S6: Synthesis of gallinamide A analogues 24-33

(2*S*)-*N*-((3*S*,*E*)-6-(2-((1,1’-biphenyl)-4-ylmethyl)-3-methoxy-5-oxo-2,5-dihydro-1*H*-pyrrol-1-yl)- 6-oxo-1-phenylhex-4-en-3-yl)-2-((*S*)-2-((*S*)-2-(dimethylamino)-3-methylbutanamido)- 4-methylpentanamido)-4-methylpentanamide. $CF_3CO_2H$ **(24)**

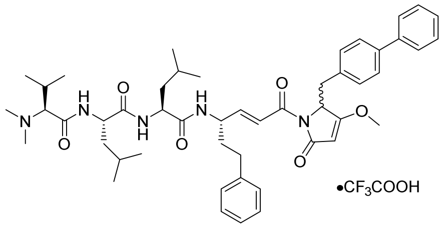


Compound **S12a** (10.0 µmol) and tripeptide **S2** (15.0 µmol) were transformed into the corresponding HCl salts according to general procedure 7 and then coupled according to general procedure 8. The crude mixture was purified by preparative RP-HPLC (gradient: 40%-60% B over 18 min, retention time: 11.8-13.0 min). The product was lyophilized to afford the *title* compound **24** as a white fluffy solid (8.10 mg, 87%, *dr* 1:1). **$[\alpha]^{20}_D$** = -31° (c = 0.1, MeOH); **IR** (thin film) $v_{max}$ = 3275, 2929, 1724, 1678, 1635, 1548, 1452, 1349, 1204, 1137 $cm^{-1}$; **$^1$H NMR** (400 MHz, DMSO-$d_6$, 1:1 mixture of diastereoisomers) δ 9.58 (br s, 1H, NH), 8.76 (d, 1H, *J* 7.6 Hz, NH), 8.32 (d, 1H, *J* 9.0 Hz, NH), 8.28 (d, 1H, *J* 7.8 Hz, NH), 7.59-7.63 (m, 2H, Ar-*H*), 7.51-7.54 (m, 2H, Ar-*H*), 7.40-7.45 (m, 2H, Ar-*H*), 7.31-7.36 (m, 1H, Ar-*H*), 7.16-7.29 (m, 6H, 5 x Ar-*H*, CH=C*H*C=O), 6.93-6.99 (m, 3H, 2 x Ar-*H*, C*H*=CHC=O), 5.21 (s, 1H, CH), 4.99 (dd, 1H, *J* 2.8, 5.4 Hz, CH), 4.51-4.57 (m, 1H, CH), 4.37-4.45 (m, 2H, 2 x CH), 3.88 (s, 3H, $OCH_3$), 3.63-3.66 (m, 1H, CH), 3.44-3.52 (m, 1H, C*H*H), 3.09 (dd, 1H, *J* 2.5, 14.0 Hz, CH*H*), 2.76 (s, 3H, $NCH_3$), 2.73 (s, 3H, $NCH_3$), 2.55-2.68 (m, 2H, $CH_2$), 2.24-2.33 (m, 1H, CH), 1.76-1.94 (m, 2H, $CH_2$), 1.47-1.64 (m, 6H, 2 x CH, 2 x $CH_2$), 1.01 (d, 3H, *J* 6.7 Hz, $CH_3$), 0.81-0.89 (m, 15H, 5 x $CH_3$); **$^{13}$C NMR** (100 MHz, DMSO-$d_6$, 1:1 mixture of diastereoisomers) δ 178.6, 172.0, 171.4, 170.0, 170.0, 165.3, 164.2, 164.2, 149.2, 149.0, 141.8, 141.8, 140.0, 140.0, 138.9, 134.0, 134.0, 130.5, 129.4,

129.4, 128.9, 128.9, 128.7, 127.9, 127.8, 126.9, 126.9, 126.7, 126.6, 126.3, 122.2, 95.5, 72.1, 59.5, 51.5, 51.5, 49.6, 49.4, 42.3, 41.3, 41.2, 41.0, 35.7, 35.7, 34.0, 34.0, 34.0, 31.9, 31.9, 31.8, 29.5, 29.5, 29.4, 29.2, 27.0, 24.7, 24.6, 24.6, 23.5, 23.5, 23.4, 21.9, 21.9, 19.7, 16.9; **HRMS** (+ESI) *m/z* Calc. for $C_{49}H_{65}N_5O_6$ [M + Na]$^+$: 842.4827, found 842.4828.

*N*-((2*S*)-1-(((2*S*)-1-(((3*S*,*E*)-6-(2-((1,1'-biphenyl)-4-ylmethyl)-3-methoxy-5-oxo-2,5-dihydro- 1*H*-pyrrol-1-yl)-6-oxo-1-phenylhex-4-en-3-yl)amino)-4-methyl-1-oxopentan-2-yl)amino)- 4-methyl-1-oxopentan-2-yl)-1-methylpiperidine-4-carboxamide. $CF_3CO_2H$ **(25)**

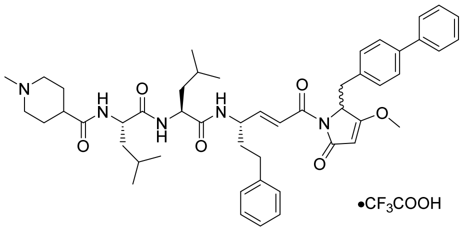

Compound **S12a** (40.0 µmol) and tripeptide **S4** (60.0 µmol) were transformed into the corresponding HCl salts according to general procedure 7 and then coupled according to general procedure 8. The crude mixture was purified by preparative RP-HPLC (gradient: 45%-65% B over 18 min, retention time: 7.8-9.1 min). The product was lyophilized to afford the *title* compound **25** as a white fluffy solid (25.9 mg, 70%, *dr* 1:1). **[α]$^{20}_{D}$** = -32° (c = 0.1, MeOH); **IR** (thin film) $\nu_{max}$ = 3291, 2954, 2926, 2854, 1723, 1673, 1633, 1546, 1450, 1348, 1200, 1130, 970 cm$^{-1}$; **$^1$H NMR** (400 MHz, DMSO-*d*$_6$, 1:1 mixture of diastereoisomers) δ 8.21 (d, 1H, *J* 8.1 Hz, NH), 8.10 (d, 1H, *J* 8.8 Hz, NH), 7.93 (d, 1H, *J* 7.5 Hz, NH), 7.57-7.62 (m, 2H, Ar-*H*), 7.52 (dd, 2H, *J* 6.5, 8.1 Hz, Ar-*H*), 7.39-7.44 (m, 2H, Ar-*H*), 7.31-7.35 (m, 1H, Ar-*H*), 7.16-7.28 (m, 6H, 5 x Ar-*H*, CH=C*H*C=O), 6.92-6.97 (m, 3H, 2 x Ar-*H*, C*H*=CHC=O), 5.20 (s, 1H, CH), 4.98 (dd, 1H, *J* 2.7, 5.4 Hz, CH), 4.40-4.47 (m, 1H, CH), 4.28-4.37 (m, 2H, 2 x CH), 3.87 (s, 3H, $OCH_3$), 3.37-3.49 (m, 2H, CH, C*H*H), 3.08 (dd, 1H, *J* 2.7, 14.3 Hz, CH*H*), 2.87-2.96 (m, 2H, $CH_2$), 2.74 (s, 3H, $NCH_3$), 2.54-2.69 (m, 2H, $CH_2$), 2.43 (t, 1H, *J* 12.2 Hz, CH), 1.67-1.93 (m, 6H, 3 x $CH_2$), 1.46-1.64 (m, 7H, CH, 3 x $CH_2$), 0.80-0.88 (m, 12H, 4 x $CH_3$); **$^{13}$C NMR** (100 MHz, DMSO-*d*$_6$, 1:1 mixture of diastereoisomers) δ 178.6, 173.2, 172.3, 172.1, 170.0, 170.0, 164.3, 164.2, 158.6 (q, *J* 30.9 Hz, $CF_3CO_2H$), 149.1, 148.9, 141.8, 141.8, 140.0, 140.0, 138.9, 138.9, 134.0, 130.5, 129.4, 129.3, 128.9, 128.9, 128.8, 128.7, 127.8, 126.8, 126.8, 126.6, 126.6, 126.3, 122.5, 122.3, 94.5, 59.5, 53.2, 53.2, 51.6, 51.4, 49.6, 49.5, 43.1, 41.4, 41.2, 38.9, 35.8, 34.0, 34.0, 31.9, 31.9, 26.7, 26.3, 24.7, 23.5, 23.4, 23.4, 22.2, 22.2, 22.0; **HRMS** (+ESI) *m/z* Calc. for $C_{49}H_{63}N_5O_6$ [M + Na]$^+$: 840.4671, found 840.4671.

(2*S*)-2-((*S*)-2-(dimethylamino)-3-methylbutanamido)-*N*-((2*S*)-1-(((3*S*,*E*)-6-(3-methoxy-5-oxo-2- ((4'-(trifluoromethyl)-(1,1'-biphenyl)-4-yl)methyl)-2,5-dihydro-1*H*-pyrrol-1-yl)-6-oxo- 1-phenylhex-4-en-3-yl)amino)-4-methyl-1-oxopentan-2-yl)-4-methylpentanamide. $CF_3CO_2H$ **(26)**

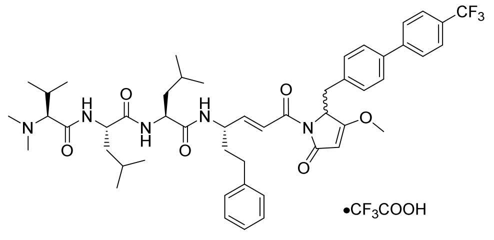

Compound **S12b** (5.00 μmol) and tripeptide **S2** (7.50 μmol) were transformed into the corresponding HCl salts according to general procedure 7 and then coupled according to general procedure 8. The crude mixture was purified by preparative RP-HPLC (gradient: 45%-65% B over 18 min, retention time: 11.4-12.1 min). The product was lyophilized to afford the *title* compound **26** as a white fluffy solid (3.80 mg, 76%, *dr* 1:1). $[\alpha]^{20}_D$ = -19° (c = 0.1, MeOH); **IR** (thin film) $v_{max}$ = 3274, 2955, 2913, 2856, 1726, 1635, 1551, 1455, 1327, 1193, 1128, 1071 $cm^{-1}$; **$^1$H NMR** (400 MHz, DMSO-$d_6$, 1:1 mixture of diastereoisomers) δ 9.71 (br s, 1H, NH), 8.76 (d, 1H, *J* 8.0 Hz, NH), 8.30 (dd, 1H, *J* 1.8, 8.6 Hz, NH), 8.26 (d, 1H, *J* 8.2 Hz, NH), 7.75-7.85 (m, 4H, Ar-*H*), 7.60 (dd, 2H, *J* 6.1, 8.2 Hz, Ar-*H*), 7.16-7.28 (m, 6H, 5 x Ar-*H*, CH=C*H*C=O), 7.00-7.02 (m, 2H, Ar-*H*), 6.95 (ddd, 1H, *J* 1.2, 5.4, 15.6 Hz, C*H*=CHC=O), 5.21 (d, 1H, *J* 1.9 Hz, CH), 4.99 (dd, 1H, *J* 2.7, 5.4 Hz, CH), 4.50-4.56 (m, 1H, CH), 4.36-4.46 (m, 2H, 2 x CH), 3.88 (s, 3H, $OCH_3$), 3.63-3.66 (m, 1H, CH), 3.44-3.51 (m, 1H, C*H*H), 3.10 (dd, 1H, *J* 2.5, 13.6 Hz, CH*H*), 2.73 (s, 6H, 2 x $NCH_3$), 2.54-2.67 (m, 2H, $CH_2$), 2.22-2.33 (m, 1H, CH), 1.79-1.94 (m, 2H, $CH_2$), 1.45-1.63 (m, 6H, 2 x CH, 2 x $CH_2$), 1.01 (d, 3H, *J* 6.8 Hz, $CH_3$), 0.79-0.89 (m, 15H, 5 x $CH_3$); **$^{13}$C NMR** (100 MHz, DMSO-$d_6$, 1:1 mixture of diastereoisomers) δ 178.5, 172.0, 171.4, 170.0, 169.9, 165.4, 164.3, 158.4, 158.1, 149.1, 149.1, 144.0, 141.8, 137.3, 135.3, 130.6, 128.9, 128.9, 128.7, 127.7, 127.7, 127.1, 126.3, 126.2, 123.5, 122.4, 122.3, 95.5, 72.0, 59.5, 59.5, 59.5, 51.6, 49.6, 49.5, 42.0, 41.5, 41.2, 41.0, 35.8, 35.7, 34.1, 34.0, 31.9, 31.9, 29.5, 27.0, 24.7, 24.6, 23.5, 23.4, 21.9, 21.9, 19.7, 17.1; **HRMS** (+ESI) *m/z* Calc. for $C_{50}H_{64}F_3N_5O_6$ $[M + Na]^+$: 910.4701, found 910.4698.

*N*-((2*S*)-1-(((2*S*)-1-(((3*S*,*E*)-6-(3-methoxy-5-oxo-2-((4'-(trifluoromethyl)-(1,1'-biphenyl)-4-yl)methyl)-2,5-dihydro-1*H*-pyrrol-1-yl)-6-oxo-1-phenylhex-4-en-3-yl)amino)-4-methyl-1-oxopentan-2-yl)amino)-4-methyl-1-oxopentan-2-yl)-1-methylpiperidine-4-carboxamide. $CF_3CO_2H$ **(27)**

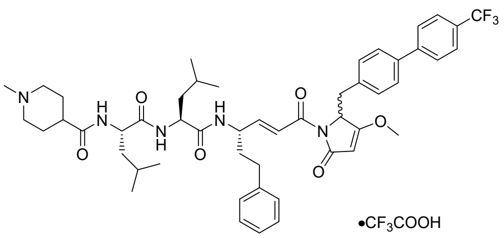

Compound **S12b** (17.0 μmol) and tripeptide **S4** (25.5 μmol) were transformed into the corresponding HCl salts according to general procedure 7 and then coupled according to general procedure 8. The crude mixture was purified by preparative RP-HPLC (gradient: 45%-65% B over 18 min, retention time: 9.4-10.8 min). The product was lyophilized to afford the *title* compound **27** as a white fluffy solid (14.1 mg, 83%, *dr* 1:1). **[α]$^{20}_{D}$** = -20° (c = 0.1, MeOH); **IR** (thin film) $v_{max}$ = 3303, 2957, 2927, 1673, 1631, 1546, 1455, 1327, 1249, 1203, 1170, 1128, 1071 cm$^{-1}$; **$^1$H NMR** (400 MHz, DMSO-$d_6$, 1:1 mixture of diastereoisomers) δ 8.20 (dd, 1H, *J* 1.9, 8.3 Hz, NH), 8.09 (d, 1H, *J* 8.3 Hz, NH), 7.93 (dd, 1H, *J* 2.8, 8.3 Hz, NH), 7.81-7.85 (m, 2H, Ar-*H*), 7.75-7.78 (m, 2H, Ar-*H*), 7.60 (dd 2H, *J* 6.8, 8.3 Hz, Ar-*H*), 7.16-7.28 (m, 6H, 5 x Ar-*H*, CH=C*H*C=O), 7.01 (d, 2H, *J* 8.1 Hz, Ar-*H*), 6.95 (dd, 1H, *J* 5.6, 15.6 Hz, C*H*=CHC=O), 5.21 (d, 1H, *J* 1.8 Hz, CH), 4.99 (dd, 1H, *J* 2.6, 5.6 Hz, CH), 4.39-4.46 (m, 1H, CH), 4.28-4.37 (m, 2H, 2 x CH), 3.88 (s, 3H, $OCH_3$), 3.40-3.50 (m, 2H, CH, C*H*H), 3.10 (dd, 1H, *J* 2.6, 14.1 Hz, CH*H*), 2.85-2.94 (m, 2H, $CH_2$), 2.74 (d, 3H, *J* 4.3 Hz, $NCH_3$), 2.53-2.69 (m, 2H, $CH_2$), 2.43 (tt, 1H, *J* 3.7, 12.1 Hz, CH), 1.67-1.91 (m, 6H, 3 x $CH_2$), 1.43-1.62 (m, 7H, CH, 3 x $CH_2$), 0.80-0.87 (m, 12H, 4 x $CH_3$); **$^{13}$C NMR** (100 MHz, DMSO-$d_6$, 1:1 mixture of diastereoisomers) δ 178.5, 173.1, 173.1, 172.3, 172.1, 170.0, 169.4, 164.3, 164.2, 158.4 (q, *J* 30.9 Hz, $CF_3CO_2H$), 149.1, 149.0, 144.0, 141.8, 141.8, 137.3, 135.3, 130.6, 128.9, 128.7, 127.7, 127.7, 127.1, 126.3, 126.2, 123.5, 122.4, 122.3, 95.5, 59.5, 59.5, 59.5, 53.2, 51.6, 51.4, 51.1, 49.6, 49.5, 43.1, 41.3, 41.2, 38.8, 35.8, 35.7, 34.1, 34.0, 31.9, 31.9, 26.7, 26.3, 24.7, 23.5, 23.4, 23.4, 22.2, 22.2, 22.0; **HRMS** (+ESI) *m/z* Calc. for $C_{50}H_{62}F_3N_5O_6$ [M + Na]$^+$: 886.4725, found 886.4724.

(2*S*)-2-((*S*)-2-(dimethylamino)-3-methylbutanamido)-*N*-((2*S*)-1-(((3*S*,*E*)-6-(3-methoxy-2-((4'-methoxy-(1,1'-biphenyl)-4-yl)methyl)-5-oxo-2,5-dihydro-1*H*-pyrrol-1-yl)-6-oxo-1-phenylhex-4-en-3-yl)amino)-4-methyl-1-oxopentan-2-yl)-4-methylpentanamide. $CF_3CO_2H$ **(28)**

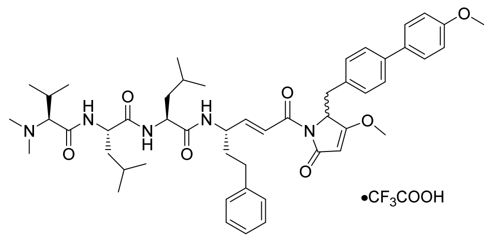


Compound **S12c** (9.40 μmol) and tripeptide **S2** (14.1 μmol) were transformed into the corresponding HCl salts according to general procedure 7 and then coupled according to general procedure 8. The crude mixture was purified by preparative RP-HPLC (gradient: 45%-65% B over 18 min, retention time: 9.2-10.0 min). The product was lyophilized to afford the *title* compound **28** as a white fluffy solid (5.90 mg, 65%, *dr* 1:1). **[α]$^{20}_{D}$** = -23° (c = 0.1, MeOH); **IR** (thin film) $v_{max}$ = 3292, 2924, 1676, 1549, 1499, 1453, 1349, 1248, 1202, 1137, 1040, 972, 805, 724 cm$^{-1}$; **$^1$H NMR** (500 MHz, DMSO-$d_6$, 1:1 mixture of diastereoisomers) δ 9.58 (br s, 1H, NH), 8.74 (d, 1H, *J* 8.6 Hz, NH), 8.29 (d, 1H, *J* 8.1 Hz, NH), 8.26 (d, 1H, *J* 8.1 Hz, NH), 7.44-7.55 (m, 4H, Ar-*H*), 7.17-7.27 (m, 6H, 5 x Ar-*H*, CH=C*H*C=O), 6.91-6.99 (m, 5H, 4 x Ar-*H*, C*H*=CHC=O), 5.19 (s, 1H, CH), 4.96 (dd, 1H, *J* 2.6, 5.3 Hz, CH), 4.50-4.55 (m, 1H, CH), 4.36-4.45 (m, 2H, 2 x CH), 3.87 (s, 3H, $OCH_3$), 3.78 (s, 3H, $OCH_3$), 3.63-3.64 (m, 1H, CH), 3.42-3.47 (m, 1H, C*H*H), 3.06 (dd, 1H, *J* 2.4, 14.0 Hz, CH*H*), 2.75 (s, 3H, $NCH_3$), 2.72 (s, 3H, $NCH_3$), 2.55-2.68 (m, 2H, $CH_2$), 2.24-2.32 (m, 1H, CH), 1.75-1.92 (m, 2H, $CH_2$), 1.43-1.63 (m, 6H, 2 x CH, 2 x $CH_2$), 1.00 (d, 3H, *J* 6.9 Hz, $CH_3$), 0.80-0.88 (m, 15H, 5 x $CH_3$); **$^{13}$C NMR** (125 MHz, DMSO-$d_6$, 1:1 mixture of diastereoisomers) δ 178.1, 171.5, 170.9, 169.6, 169.5, 164.9, 163.8, 163.8, 158.3 (q, *J* 31.3 Hz, $CF_3CO_2H$), 148.7, 148.5, 141.4, 141.3, 138.1, 132.7, 132.7, 131.9, 131.9, 129.9, 128.5, 128.4,

128.2, 127.5, 127.5, 125.8, 125.6, 125.6, 122.0, 121.8, 114.3, 114.3, 95.0, 71.6, 59.1, 59.1, 59.0, 55.1, 51.1, 51.0, 49.1, 49.0, 41.8, 40.8, 40.7, 40.5, 35.3, 35.2, 33.5, 33.4, 31.4, 31.4, 31.3, 29.0, 29.0, 29.0, 29.0, 28.7, 26.5, 24.2, 24.2, 24.2, 23.0, 23.0, 23.0, 22.1, 21.5, 21.4, 19.2, 16.4; **HRMS** (+ESI) *m/z* Calc. for $C_{50}H_{67}N_5O_7$ $[M + Na]^+$: 872.4933, found 872.4936.

*N*-((2*S*)-1-(((2*S*)-1-(((3*S*,*E*)-6-(3-methoxy-2-((4'-methoxy-(1,1'-biphenyl)-4-yl)methyl)-5-oxo-2,5-dihydro-1*H*-pyrrol-1-yl)-6-oxo-1-phenylhex-4-en-3-yl)amino)-4-methyl-1-oxopentan-2-yl)amino)-4-methyl-1-oxopentan-2-yl)-1-methylpiperidine-4-carboxamide. $CF_3CO_2H$ **(29)**

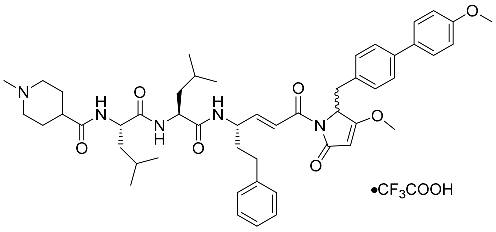


Compound **S12c** (9.40 µmol) and tripeptide **S4** (14.1 µmol) were transformed into the corresponding HCl salts according to general procedure 7 and then coupled according to general procedure 8. The crude mixture was purified by preparative RP-HPLC (gradient: 45%-65% B over 18 min, retention time: 8.2-9.0 min). The product was lyophilized to afford the *title* compound **29** as a white fluffy solid (5.90 mg, 68%, *dr* 1:1). $[\alpha]^{20}_D$ = -25° (c = 0.1, MeOH); **IR** (thin film) $\nu_{max}$ = 3303, 2957, 2923, 2850, 1678, 1545, 1699, 1447, 1348, 1249, 1207, 1181, 1135, 1041, 963, 801, 724 $cm^{-1}$; $^1$**H NMR** (500 MHz, DMSO-$d_6$, 1:1 mixture of diastereoisomers) δ 9.31 (br s, 1H, NH), 8.21 (d, 1H, *J* 8.3 Hz, NH), 8.08 (d, 1H, *J* 8.0 Hz, NH), 7.93 (dd, 1H, *J* 1.2, 8.1 Hz, NH), 7.51-7.55 (m, 2H, Ar-*H*), 7.46 (t, 2H, *J* 7.8 Hz, Ar-*H*), 7.17-7.28 (m, 6H, 5 x Ar-*H*, CH=C*H*C=O), 6.91-6.99 (m, 5H, 4 x Ar-*H*, C*H*=CHC=O), 5.19 (s, 1H, CH), 4.97 (dd, 1H, *J* 2.5, 5.4 Hz, CH), 4.39-4.46 (m, 1H, CH), 4.29-4.36 (m, 2H, 2 x CH), 3.87 (s, 3H, $OCH_3$), 3.78 (d, 3H, *J* 1.9 Hz, $OCH_3$), 3.25-3.27 (m, 1H, C*H*H), 3.05 (dd, 1H, *J* 2.2, 14.3 Hz, CH*H*), 2.86-2.95 (m, 2H, $CH_2$), 2.75 (d, 3H, *J* 4.3 Hz, $NCH_3$), 2.54-2.68 (m, 2H, $CH_2$), 2.42 (tt, 1H, *J* 3.7, 12.1 Hz, CH), 1.40-2.02 (m, 13H, CH, 6 x $CH_2$), 0.80-0.88 (m, 12H, 4 x $CH_3$); $^{13}$**C NMR** (125 MHz, DMSO-$d_6$, 1:1 mixture of diastereoisomers) δ 178.7, 173.3, 172.4, 172.2, 170.2, 170.1, 164.4, 164.4, 159.4, 159.4, 149.1, 142.0, 142.0, 138.7, 133.3, 132.5, 132.5, 130.5, 129.1, 129.1, 128.9, 128.1, 128.1, 126.4, 126.2, 122.6, 122.4, 114.9, 114.9, 95.6, 59.7, 55.8, 53.4, 53.4, 51.7, 51.5, 51.2, 49.8, 49.6, 43.2, 41.5, 41.3, 39.0, 35.9, 35.9, 34.1, 34.1, 32.0, 32.0, 26.8, 26.4, 24.9, 23.6, 23.6, 23.5, 22.4, 22.1; **HRMS** (+ESI) *m/z* Calc. for $C_{50}H_{65}N_5O_7$ $[M + Na]^+$: 870.4776, found 870.4777.

(2*S*)-*N*-((3*S*,*E*)-6-(2-((4'-cyano-(1,1'-biphenyl)-4-yl)methyl)-3-methoxy-5-oxo-2,5-dihydro-1*H*-pyrrol-1-yl)-6-oxo-1-phenylhex-4-en-3-yl)-2-((*S*)-2-((*S*)-2-(dimethylamino)-3-methylbutanamido)-4-methylpentanamido)-4-methylpentanamide. $CF_3CO_2H$ **(30)**

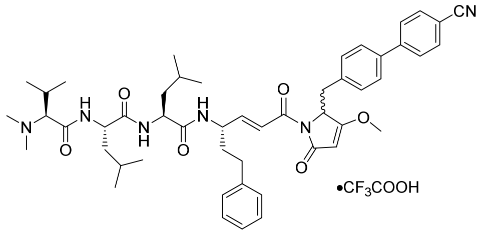

Compound **S12d** (21.0 μmol) and tripeptide **S2** (31.5 μmol) were transformed into the corresponding HCl salts according to general procedure 7 and then coupled according to general procedure 8. The crude mixture was purified by preparative RP-HPLC (gradient: 45%-65% B over 18 min, retention time: 8.2-9.4 min). The product was lyophilized to afford the *title* compound **30** as a white fluffy solid (18.5 mg, 92%, *dr* 1:1). $[\alpha]^{20}_D$ = -18° (c = 0.1, MeOH); **IR** (thin film) $\nu_{max}$ = 3288, 2958, 2926, 2869, 1674, 1635, 1546, 1508, 1455, 1351, 1199, 1134, 981, 619 $cm^{-1}$; **$^1$H NMR** (400 MHz, DMSO-$d_6$, 1:1 mixture of diastereoisomers) δ 9.70 (br s, 1H, NH), 8.76 (d, 1H, *J* 8.7 Hz, NH), 8.30 (dd, 1H, *J* 2.5, 8.7 Hz, NH), 8.26 (dd, 1H, *J* 1.4, 8.4 Hz, NH), 7.80-7.89 (m, 4H, Ar-*H*), 7.2 (dd, 2H, *J* 5.8, 7.8 Hz, Ar-*H*), 7.16-7.28 (m, 6H, 5 x Ar-*H*, CH=C*H*C=O), 6.96-7.02 (m, 2H, Ar=*H*), 6.95 (dd, 1H, *J* 5.0, 15.4 Hz, C*H*=CHC=O), 5.20 (d, 1H, *J* 2.0 Hz, CH), 4.99 (dd, 1H, *J* 2.7, 5.3 Hz, CH), 4.50-4.56 (m, 1H, CH), 4.36-4.44 (m, 2H, 2 x CH), 3.87 (s, 3H, $OCH_3$), 3.63-3.66 (m, 1H, CH), 3.44-3.50 (m, 1H, C*H*H), 3.10 (dd, 1H, *J* 2.1, 14.3 Hz, CH*H*), 2.74 (s, 6H, 2 x $NCH_3$), 2.54-2.70 (m, 2H, $CH_2$), 2.23-2.33 (m, 1H, CH), 1.75-1.93 (m, 2H, $CH_2$), 1.44-1.67 (m, 6H, 2 x CH, 2 x $CH_2$), 1.01 (d, 3H, *J* 6.8 Hz, $CH_3$), 0.79-0.89 (m, 15H, 5 x $CH_3$); **$^{13}$C NMR** (100 MHz, DMSO-$d_6$, 1:1 mixture of diastereoisomers) δ 178.5, 172.0, 171.4, 170.0, 169.9, 165.4, 164.3, 164.2, 158.5 (q, *J* 32.5 Hz, $CF_3CO_2H$), 149.2, 149.1, 144.5, 144.5, 141.8, 137.0, 135.7, 135.7, 133.3, 133.3, 130.7, 128.9, 128.9, 128.7, 127.7, 127.7, 127.1, 127.1, 126.3, 122.3, 122.2, 119.3, 110.4, 110.4, 95.5, 72.0, 59.5, 59.5, 59.5, 51.6, 51.5, 49.6, 49.5, 42.0, 41.4, 41.2, 41.2, 41.0, 35.7, 35.7, 34.1, 34.0, 31.9, 31.9, 27.0, 24.7, 24.6, 24.6, 23.5, 23.4, 21.9, 21.9, 19.7, 17.0; **HRMS** (+ESI) *m/z* Calc. for $C_{50}H_{64}N_6O_6$ $[M + Na]^+$: 867.4780, found 867.4778.

*N*-((2*S*)-1-(((2*S*)-1-(((3*S*,*E*)-6-(2-((4'-cyano-(1,1'-biphenyl)-4-yl)methyl)-3-methoxy-5-oxo-2,5-dihydro-1*H*-pyrrol-1-yl)-6-oxo-1-phenylhex-4-en-3-yl)amino)-4-methyl-1-oxopentan-2-yl)amino)-4-methyl-1-oxopentan-2-yl)-1-methylpiperidine-4-carboxamide. $CF_3CO_2H$ **(31)**

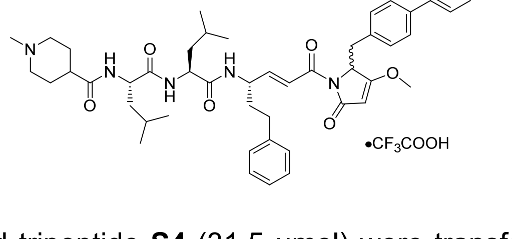

Compound **S12d** (21.0 μmol) and tripeptide **S4** (31.5 μmol) were transformed into the corresponding HCl salts according to general procedure 7 and then coupled according to general procedure 8. The crude mixture was purified by preparative RP-HPLC (gradient: 42%-62% B over 18 min, retention time: 8.7-9.8 min). The product was lyophilized to afford the *title* compound **31** as a white fluffy solid (13.0 mg, 65%, *dr* 1:1). $[\alpha]^{20}_D$ =

-18° (c = 0.1, MeOH); **IR** (thin film) $v_{max}$ = 3295, 2959, 2906, 1676, 1631, 1540, 1454, 1349, 1205, 1134, 963, 837, 801, 723 $cm^{-1}$; **$^1$H NMR** (400 MHz, DMSO-$d_6$, 1:1 mixture of diastereoisomers) δ 8.20 (dd, 1H, *J* 2.7, 8.4 Hz, NH), 8.09 (d, 1H, *J* 7.9 Hz, NH), 7.93 (dd, 1H, *J* 3.7, 8.4 Hz, NH), 7.80-7.89 (m, 4H, Ar-*H*), 7.62 (dd, 2H, *J* 6.7, 8.4 Hz, Ar-*H*), 7.16-7.28 (m, 6H, 5 x Ar-*H*, CH=C*H*C=O), 7.00-7.02 (m, 2H, Ar-*H*), 6.95 (dd, 1H, *J* 5.4, 15.4 Hz, C*H*=CHC=O), 5.20 (d, 1H, *J* 2.2 Hz, CH), 4.99 (dd, 1H, *J* 3.0, 5.4 Hz, CH), 4.39-4.46 (m, 1H, CH), 4.28-4.37 (m, 2H, 2 x CH), 3.87 (s, 3H, $OCH_3$), 3.41-3.50 (m, 2H, CH, C*H*H), 3.10 (dd, 1H, *J* 2.5, 14.1 Hz, CH*H*), 2.85-2.94 (m, 2H, $CH_2$), 2.75 (d, 3H, *J* 4.2 Hz, $NCH_3$), 2.53-2.68 (m, 2H, $CH_2$), 2.43 (tt, 1H, *J* 3.7, 12.1 Hz, CH), 1.65-1.91 (m, 6H, 3 x $CH_2$), 1.41-1.63 (m, 7H, CH, 3 x $CH_2$), 0.80-0.87 (m, 12H, 4 x $CH_3$); **$^{13}$C NMR** (100 MHz, DMSO-$d_6$, 1:1 mixture of diastereoisomers) 178.0, 172.7, 171.8, 171.6, 169.5, 169.4, 163.8, 158.4 (q, *J* 31.6 Hz, $CF_3CO_2H$), 148.7, 148.6, 144.0, 144.0, 141.3, 136.5, 135.2, 132.8, 132.8, 130.2, 128.4, 128.4, 128.2, 127.2, 127.2, 126.6, 126.6, 125.8, 121.9, 121.8, 118.8, 109.9, 95.0, 59.1, 59.0, 59.0, 52.8, 52.7, 51.1, 50.9, 49.1, 49.0, 42.6, 40.9, 40.7, 38.3, 35.2, 33.6, 31.4, 31.4, 26.2, 25.8, 24.2, 23.0, 22.9, 22.9, 21.7, 21.7, 21.5; **HRMS** (+ESI) *m/z* Calc. for $C_{50}H_{62}N_6O_6$ $[M + Na]^+$: 865.4623, found 865.4620.

(2*S*)-2-((*S*)-2-(dimethylamino)-3-methylbutanamido)-*N*-((2*S*)-1-(((3*S*,*E*)-6-(3-methoxy-2-(4-(naphthalen-2-yl)benzyl)-5-oxo-2,5-dihydro-1H-pyrrol-1-yl)-6-oxo-1-phenylhex-4-en-3-yl)amino)-4-methyl-1-oxopentan-2-yl)-4-methylpentanamide. $CF_3CO_2H$ **(32)**

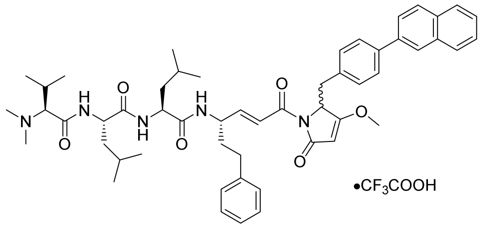


Compound **S12e** (40.0 µmol) and tripeptide **S2** (60.0 µmol) were transformed into the corresponding HCl salts according to general procedure 7 and then coupled according to general procedure 8. The crude mixture was purified by preparative RP-HPLC (gradient: 50%-70% B over 18 min, retention time: 8.6-10.7 min). The product was lyophilized to afford the *title* compound **32** as a white fluffy solid (34.9 mg, 89%, *dr* 1:1). **$[\alpha]^{20}_D$** = -21° (c = 0.1, MeOH); **IR** (thin film) $v_{max}$ = 3289, 2957, 2924, 2871, 1947, 1726, 1671, 1636, 1542, 1349, 1203, 1134, 971 $cm^{-1}$; **$^1$H NMR** (400 MHz, DMSO-$d_6$, 1:1 mixture of diastereoisomers) δ 9.75 (br s, 1H, NH), 8.7 (d, 1H, *J* 6.4 Hz, NH), 8.27-8.30 (m, 2H, 2 x NH), 8.16 (d, 1H, *J* 7.4 Hz, Ar-*H*), 7.91-7.98 (m, 4H, Ar-*H*), 7.79 (dt, 1H, *J* 1.7, 8.6 Hz, Ar-*H*), 7.68 (dd, 1H, *J* 5.9, 8.0 Hz), 7.49-7.55 (m, 2H, Ar-*H*), 7.16-7.28 (m, 6H, 5 x Ar=*H*, CH=C*H*C=O), 6.95-7.02 (m, 3H, 2 x Ar-*H*, C*H*=CHC=O), 5.20 (s, 1H, CH), 4.99 (dd, 1H, *J* 2.7, 5.3 Hz, CH), 4.52 (dd, 1H, *J* 8.2, 18.1 Hz, CH), 4.38-4.48 (m, 2H, 2 x CH), 3.89 (s, 3H, $OCH_3$), 3.65 (br s, 1H, CH), 3.47-3.53 (m, 1H, C*H*H), 3.11 (dd, 1H, *J* 1.9, 14.5 Hz, CH*H*), 2.73 (s, 6H, 2 x $NCH_3$), 2.55-2.67 (m, 2H, $CH_2$), 2.21-2.29 (m, 1H, CH), 1.77-1.94 (m, 2H, $CH_2$), 1.44-1.68 (m, 6H, 2 x CH, 2 x $CH_2$), 1.00 (d, 3H, *J* 6.5 Hz, $CH_3$), 0.80-0.89 (m, 15H, 5 x $CH_3$); **$^{13}$C NMR** (100 MHz, DMSO-$d_6$, 1:1 mixture of diastereoisomers) 178.6, 172.0, 171.4, 170.0, 170.0, 164.3, 164.3, 158.5 (q, *J* 30.3 Hz, $CF_3CO_2H$), 149.1, 149.1, 141.8, 138.7, 137.4, 137.3, 134.2, 133.8, 132.7, 130.5, 128.9, 128.9, 128.9, 128.8, 128.7, 128.6, 127.9, 126.9, 126.8, 126.5, 126.3, 125.4, 125.3, 122.4, 95.5, 72.0, 59.5, 51.6, 51.5, 49.6, 49.5, 41.2, 41.0, 35.8, 35.7, 34.1, 34.0,

31.9, 31.9, 27.0, 24.7, 24.7, 24.6, 23.5, 23.5, 23.4, 21.9, 21.9, 19.6, 17.1; **HRMS** (+ESI) *m/z* Calc. for $C_{53}H_{67}N_5O_6$ $[M + Na]^+$: 892.4984, found 892.4981.

*N*-((2*S*)-1-(((2*S*)-1-(((3*S*,*E*)-6-(3-methoxy-2-(4-(naphthalen-2-yl)benzyl)-5-oxo-2,5-dihydro-1*H*-pyrrol-1-yl)-6-oxo-1-phenylhex-4-en-3-yl)amino)-4-methyl-1-oxopentan-2-yl)amino)-4-methyl-1-oxopentan-2-yl)-1-methylpiperidine-4-carboxamide. $CF_3CO_2H$ **(33)**

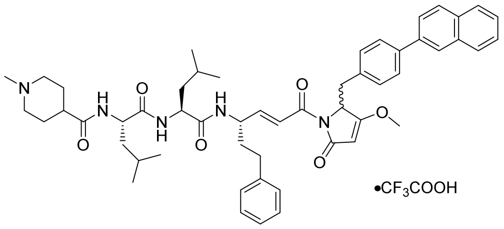


Compound **S12e** (31.0 µmol) and tripeptide **S4** (46.5 µmol) were transformed into the corresponding HCl salts according to general procedure 7 and then coupled according to general procedure 8. The crude mixture was purified by preparative RP-HPLC (gradient: 50%-70% B over 18 min, retention time: 7.5-8.9 min). The product was lyophilized to afford the *title* compound **33** as a white fluffy solid (29.5 mg, 97%, *dr* 1:1). **$[\alpha]^{20}_D$** = -18° (c = 0.1, MeOH); **IR** (thin film) $\nu_{max}$ = 3305, 2959, 1674, 1630, 1542, 1454, 1348, 1203, 1132, 962, 805 $cm^{-1}$; **$^1$H NMR** (400 MHz, DMSO-$d_6$, 1:1 mixture of diastereoisomers) δ 8.22 (d, 1H, *J* 8.2 Hz, NH), 8.16 (d, 1H, *J* 9.0 Hz, NH), 8.10 (d, 1H, *J* 8.0 Hz, NH), 7.91-7.98 (m, 4H, Ar-*H*), 7.76-7.81 (m, 1H, Ar-*H*), 7.68 (dd, 2H, *J* 6.6, 8.1 Hz, Ar-*H*), 7.49-4.55 (m, 2H, Ar-*H*), 7.15-7.28 (m, 6H, 5 x Ar-*H*, CH=C*H*C=O), 6.94-7.02 (m, 3H, 2 x Ar-*H*), C*H*=CHC=O), 5.21 (s, 1H, CH), 5.00 (dd, 1H, *J* 2.7, 5.5 Hz, CH), 4.40-4.47 (m, 1H, CH), 4.29-4.38 (m, 2H, 2 x CH), 3.89 (s, 3H, OCH3), 3.40-3.52 (m, 2H, CH, C*H*H), 3.11 (dd, 1H, *J* 2.3, 14.1 Hz, CH*H*), 2.86-2.96 (m, 2H, CH2), 2.74 (s, 3H, NCH3), 2.54-2.69 (m, 2H, CH2), 2.40-2.46 (m, 1H, CH), 1.70-1.90 (m, 6H, 3 x CH2), 1.46-1.67 (m, 7H, CH, 3 x CH2), 0.80-0.88 (m, 12H, 4 x CH3); **$^{13}$C NMR** (100 MHz, DMSO-$d_6$, 1:1 mixture of diastereoisomers) 178.1, 172.7, 171.8, 171.6, 169.5, 169.5, 163.8, 163.8, 158.1 (q, *J* 32.4 Hz, $CF_3CO_2H$), 148.6, 148.5, 141.4, 141.3, 138.3, 138.2, 136.9, 133.7, 133.3, 133.3, 132.2, 130.1, 128.4, 128.4, 128.4, 128.3, 128.2, 128.1, 127.4, 126.5, 126.4, 126.1, 125.8, 125.0, 122.0, 121.8, 95.0, 59.0, 52.8, 52.7, 51.1, 50.9, 49.1, 49.0, 42.6, 40.9, 40.7, 38.4, 35.3, 35.3, 33.6, 33.5, 31.4, 31.4, 26.2, 25.8, 24.2, 23.0, 22.9, 22.9, 21.7, 21.7, 21.5; **HRMS** (+ESI) *m/z* Calc. for $C_{53}H_{65}N_5O_6$ $[M + Na]^+$: 890.4827, found 890.4824.

### Section S7: NMR spectra for gallinamide A analogues 24-33

**Analogue 24** $^{1}$H NMR (400 MHz, DMSO-$d_6$)

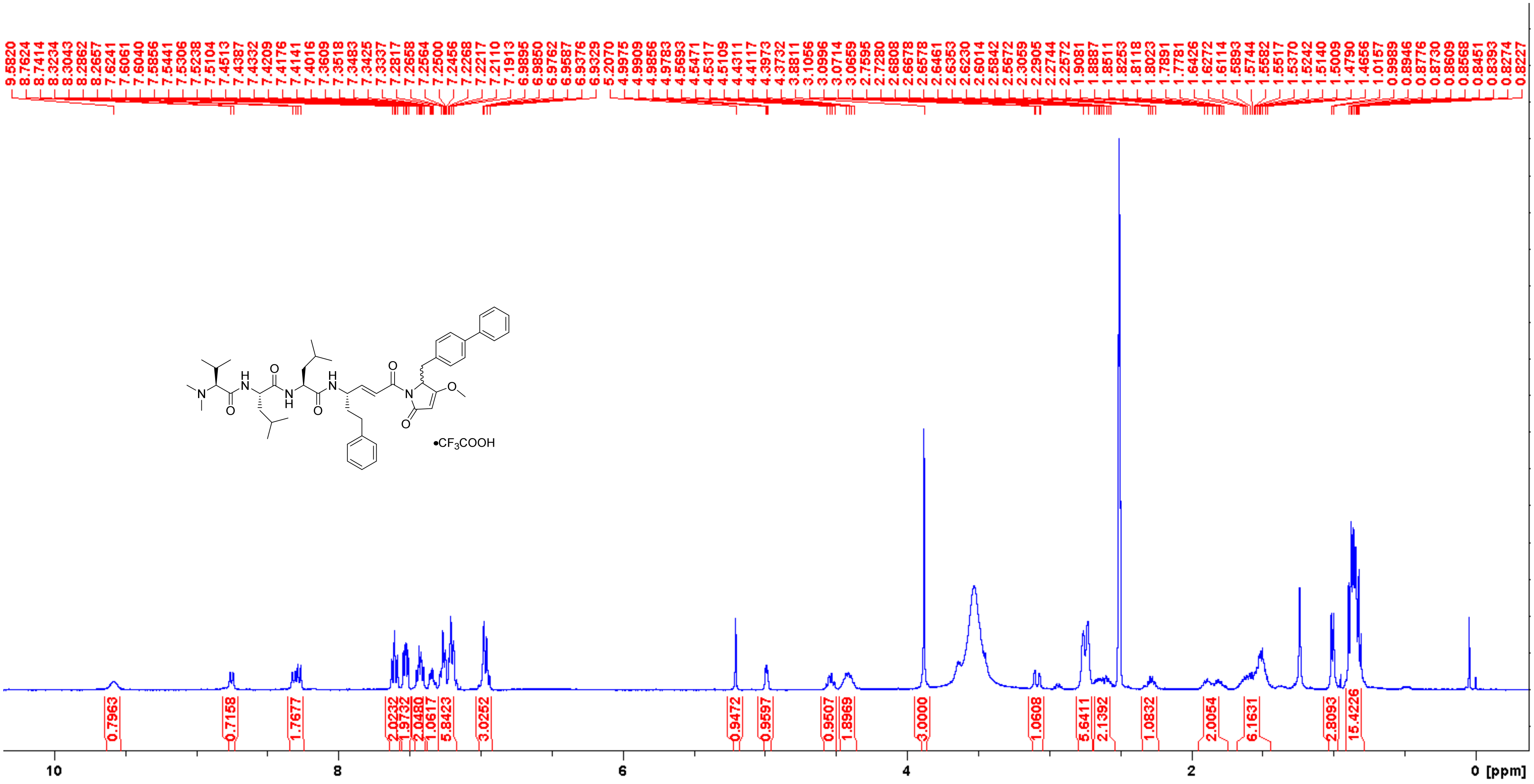

**Analogue 24** $^{13}C$ NMR (100 MHz, DMSO-$d_6$)

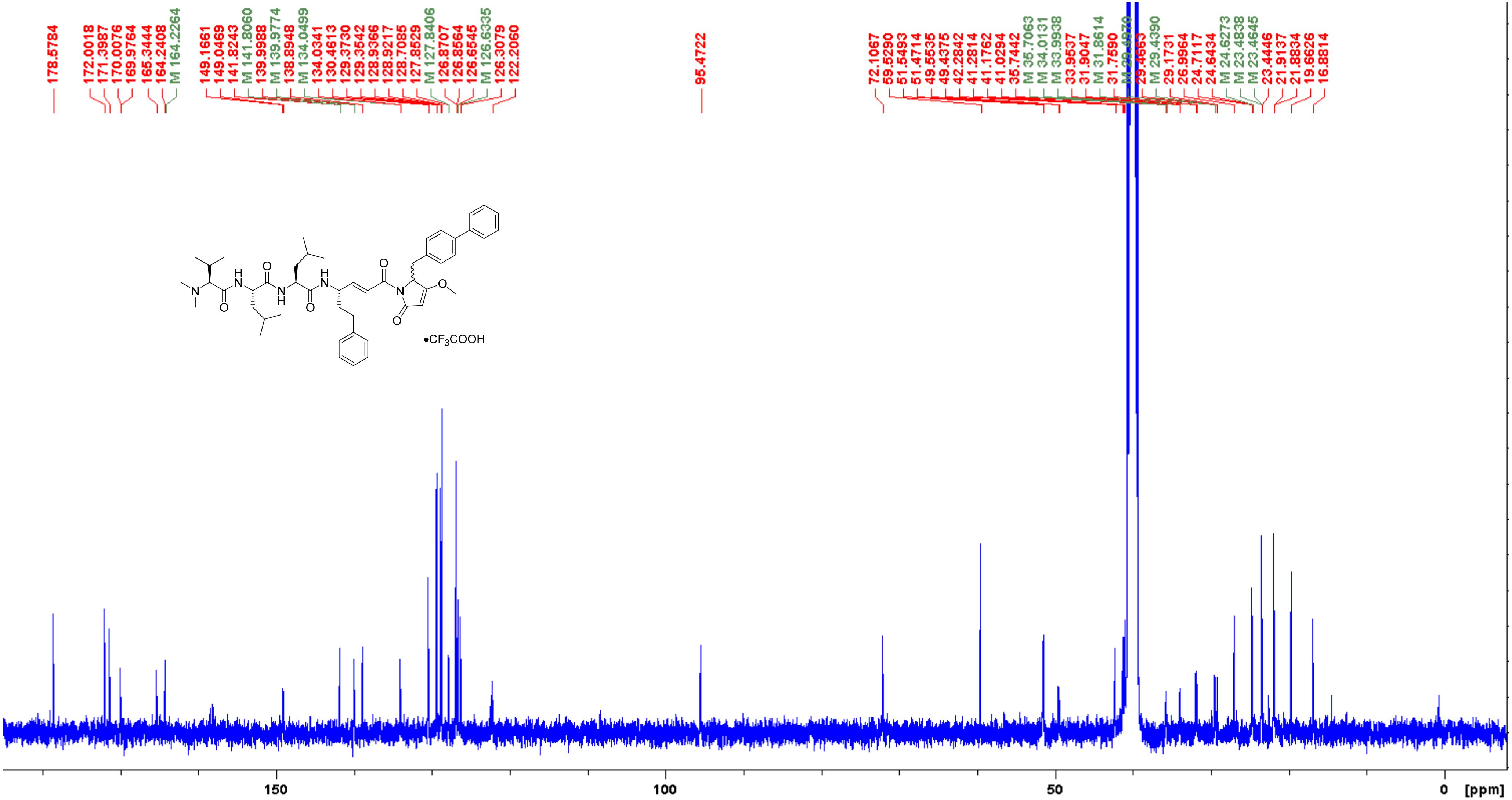

**Analogue 24** COSY (400 MHz, DMSO-$d_6$)

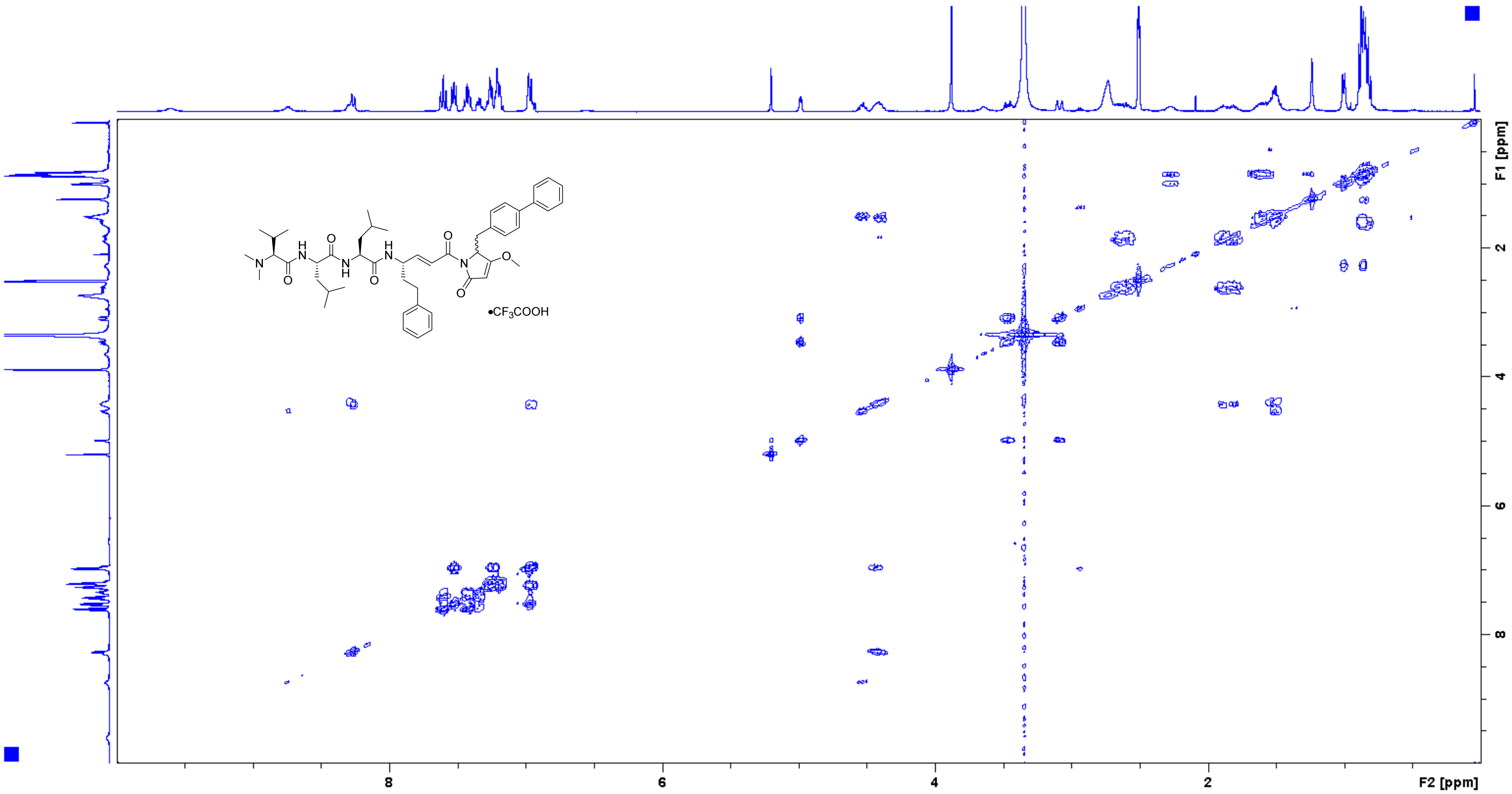

**Analogue 24** HSQC (400/100 MHz, DMSO-$d_6$)

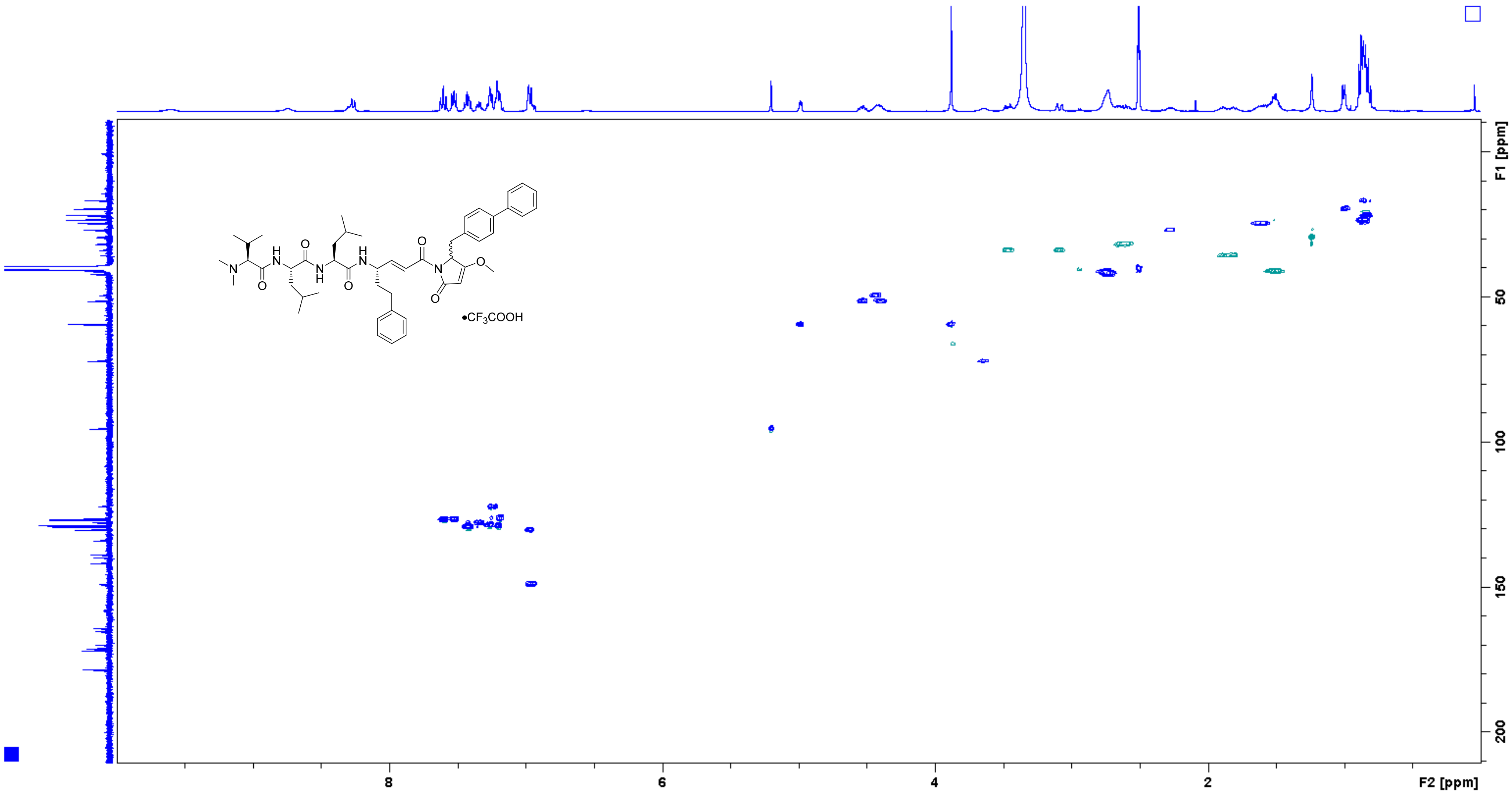

**Analogue 24** HMBC (400/100 MHz, DMSO-$d_6$)

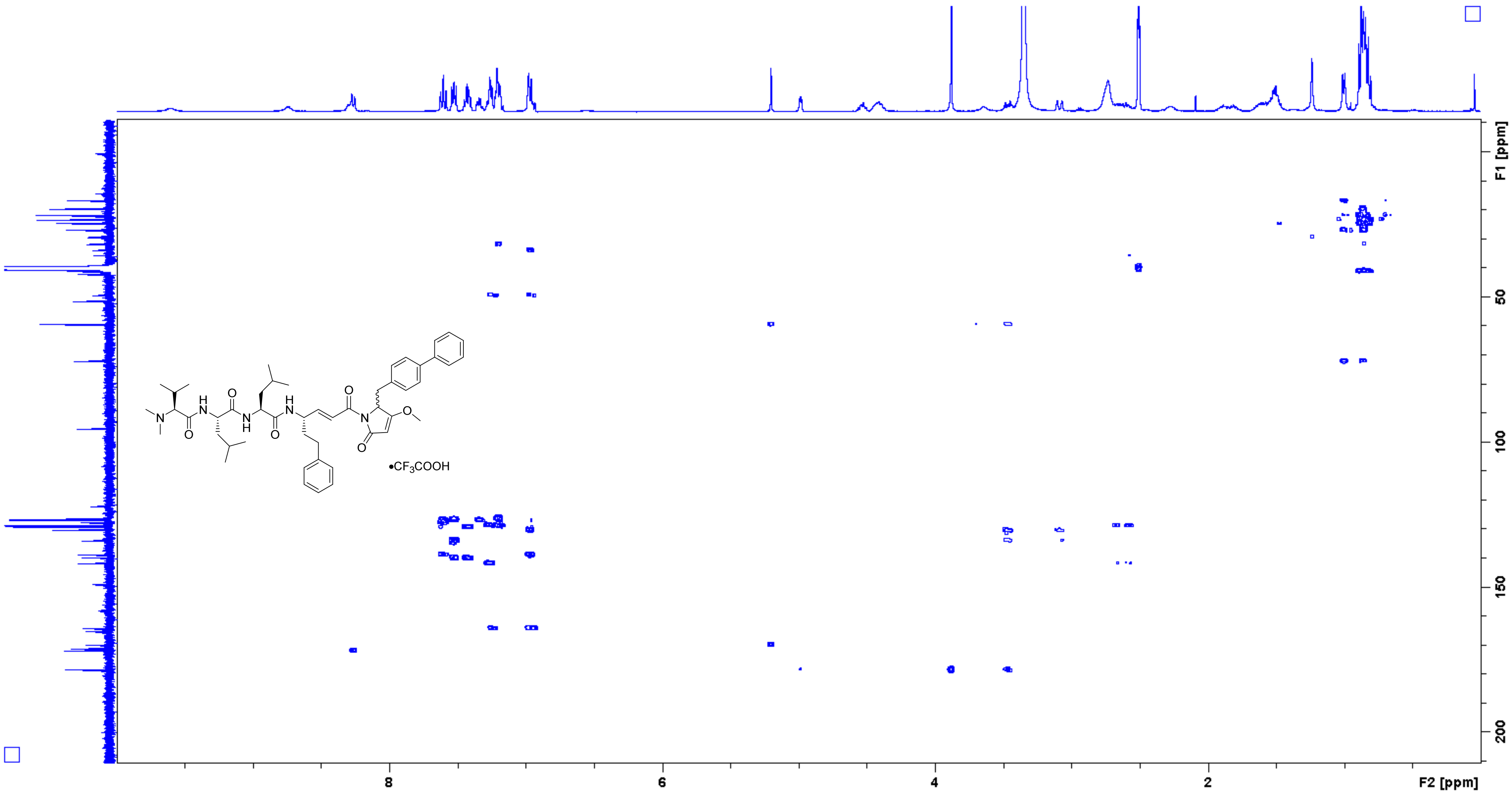

**Analogue 25** $^{1}$H NMR (400 MHz, DMSO-$d_6$)

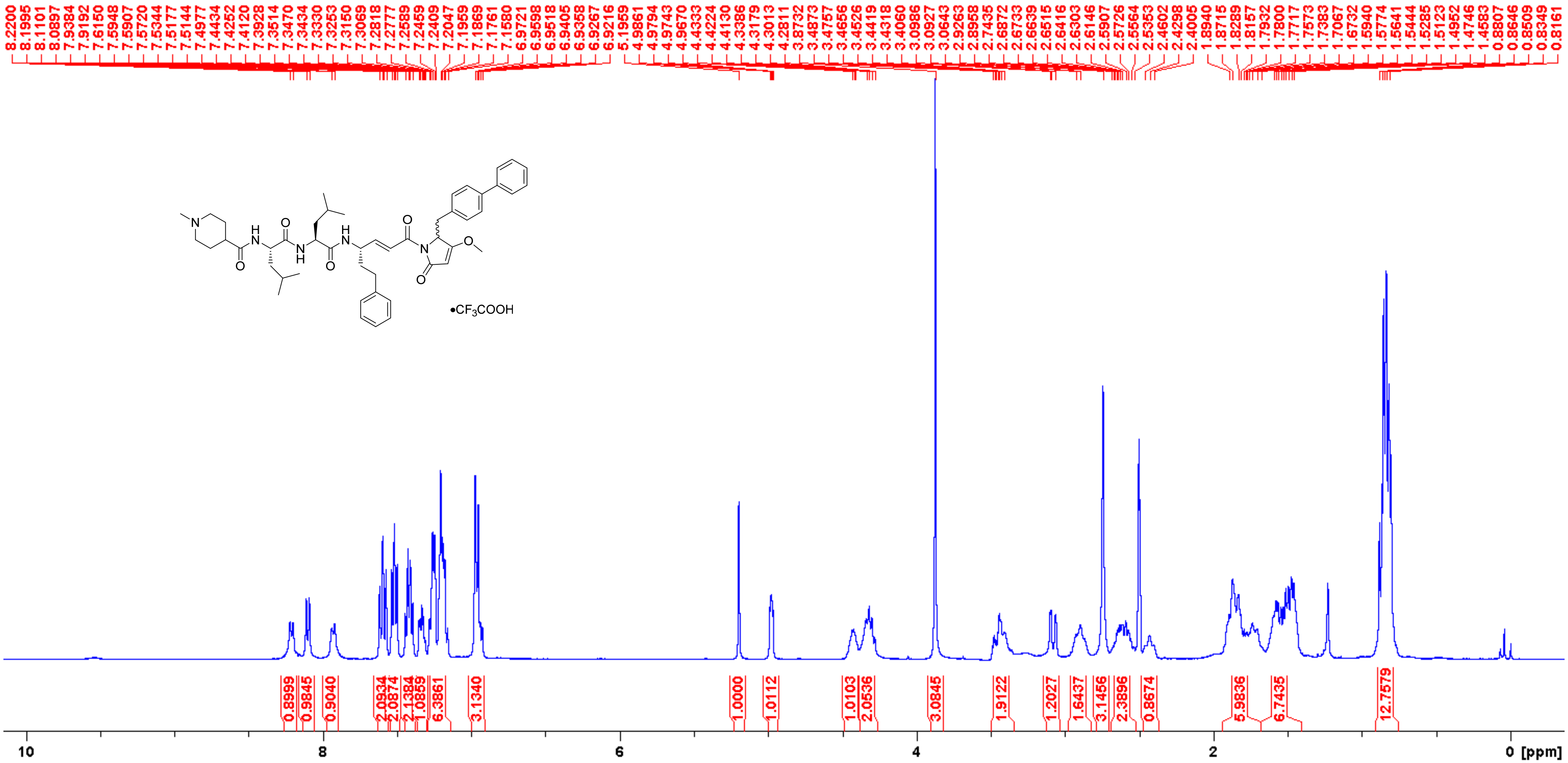

**Analogue 25** $^{13}$C NMR (100 MHz, DMSO-$d_6$)

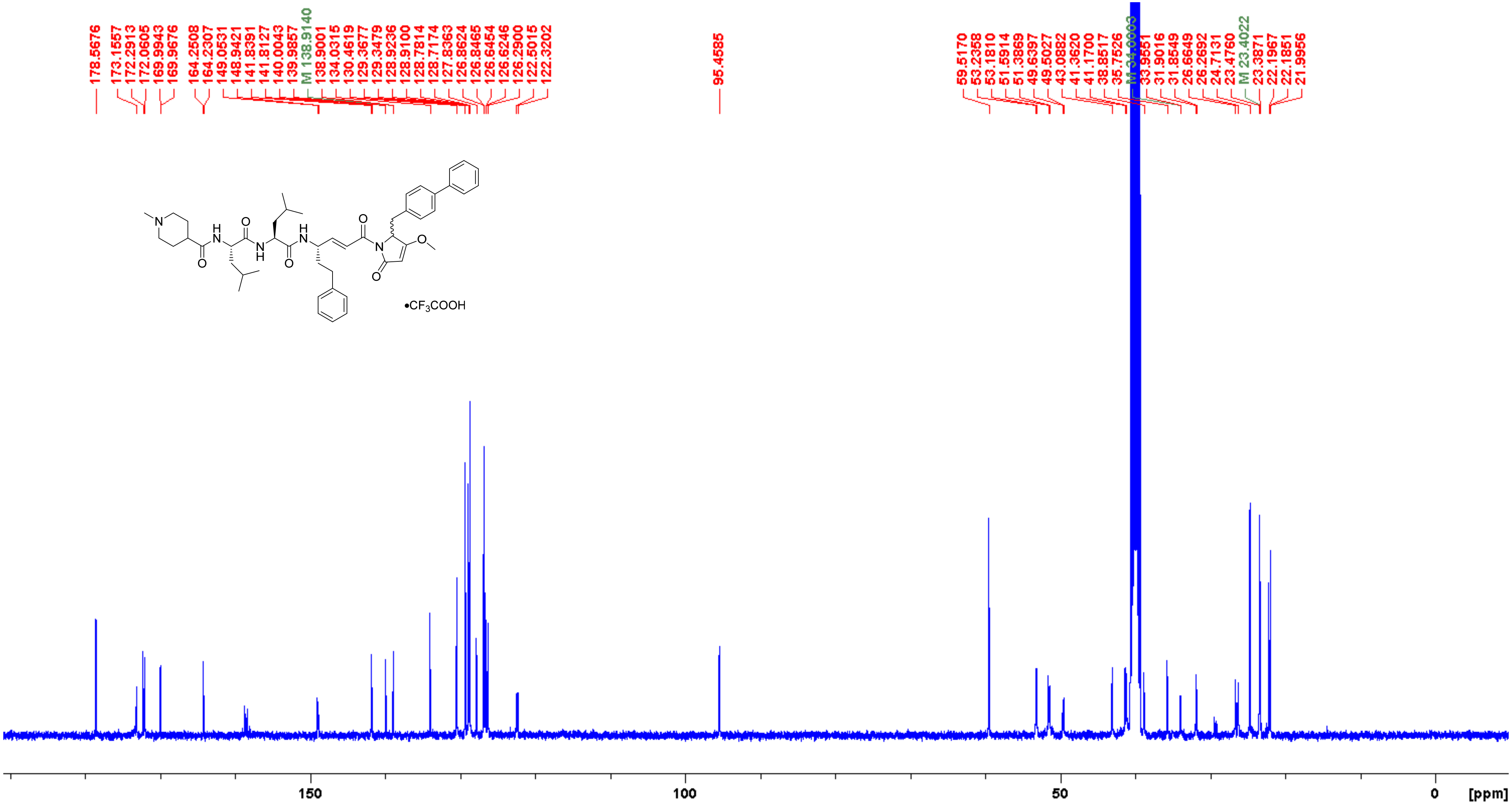

**Analogue 25** COSY (400 MHz, DMSO-$d_6$)

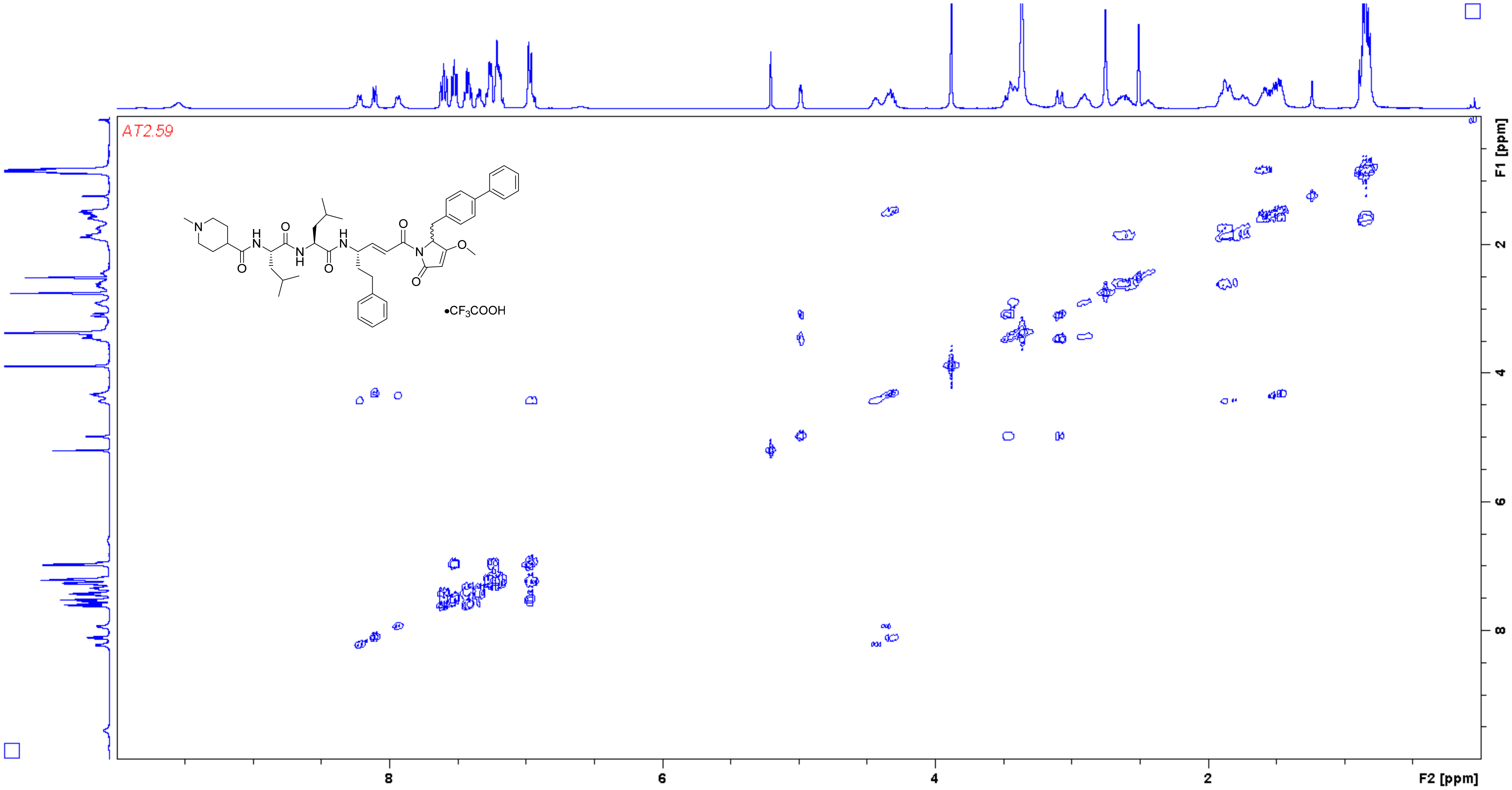

**Analogue 25** HSQC (400/100 MHz, DMSO-$d_6$)

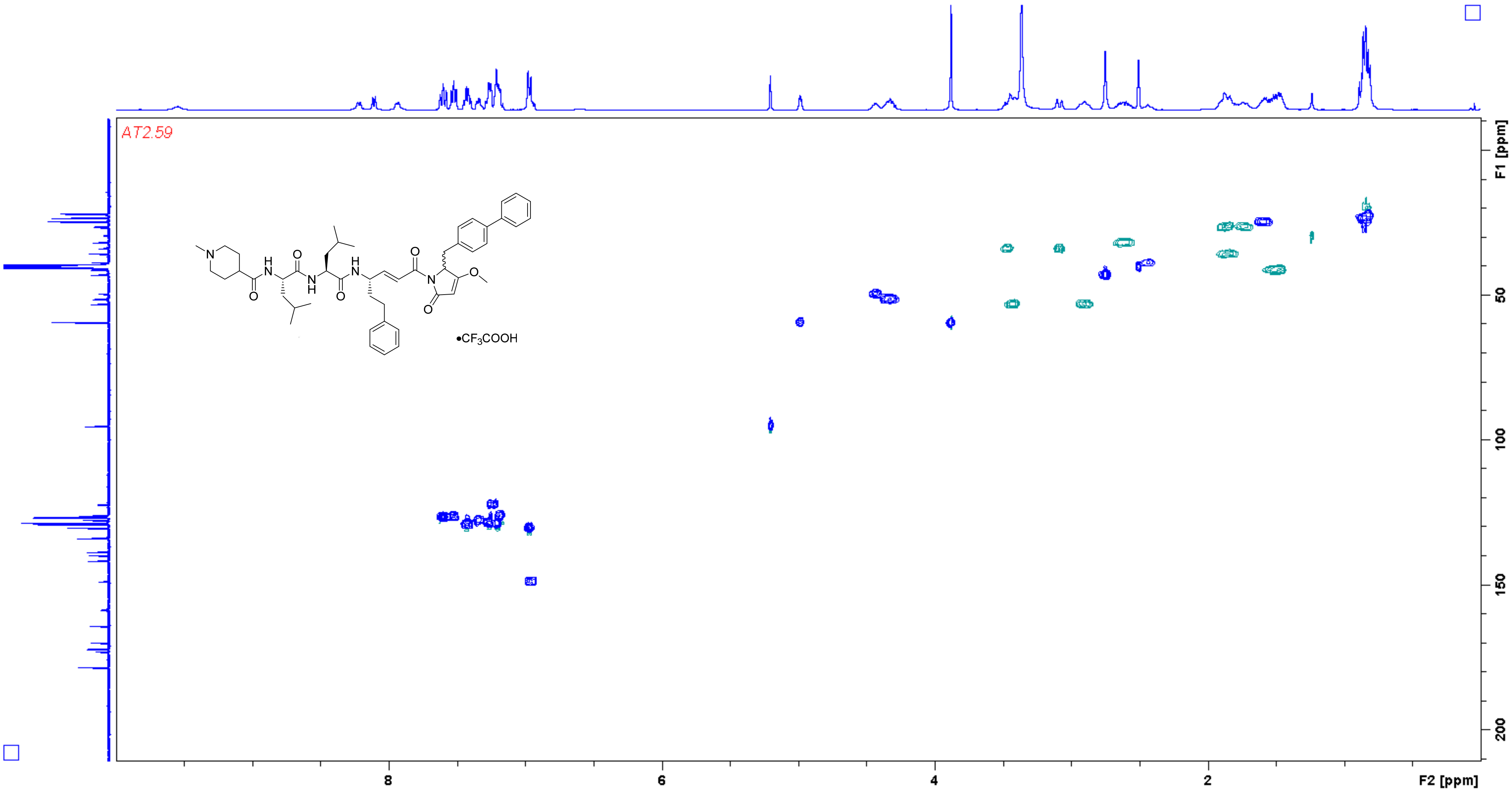

**Analogue 25** HMBC (400/100 MHz, DMSO-$d_6$)

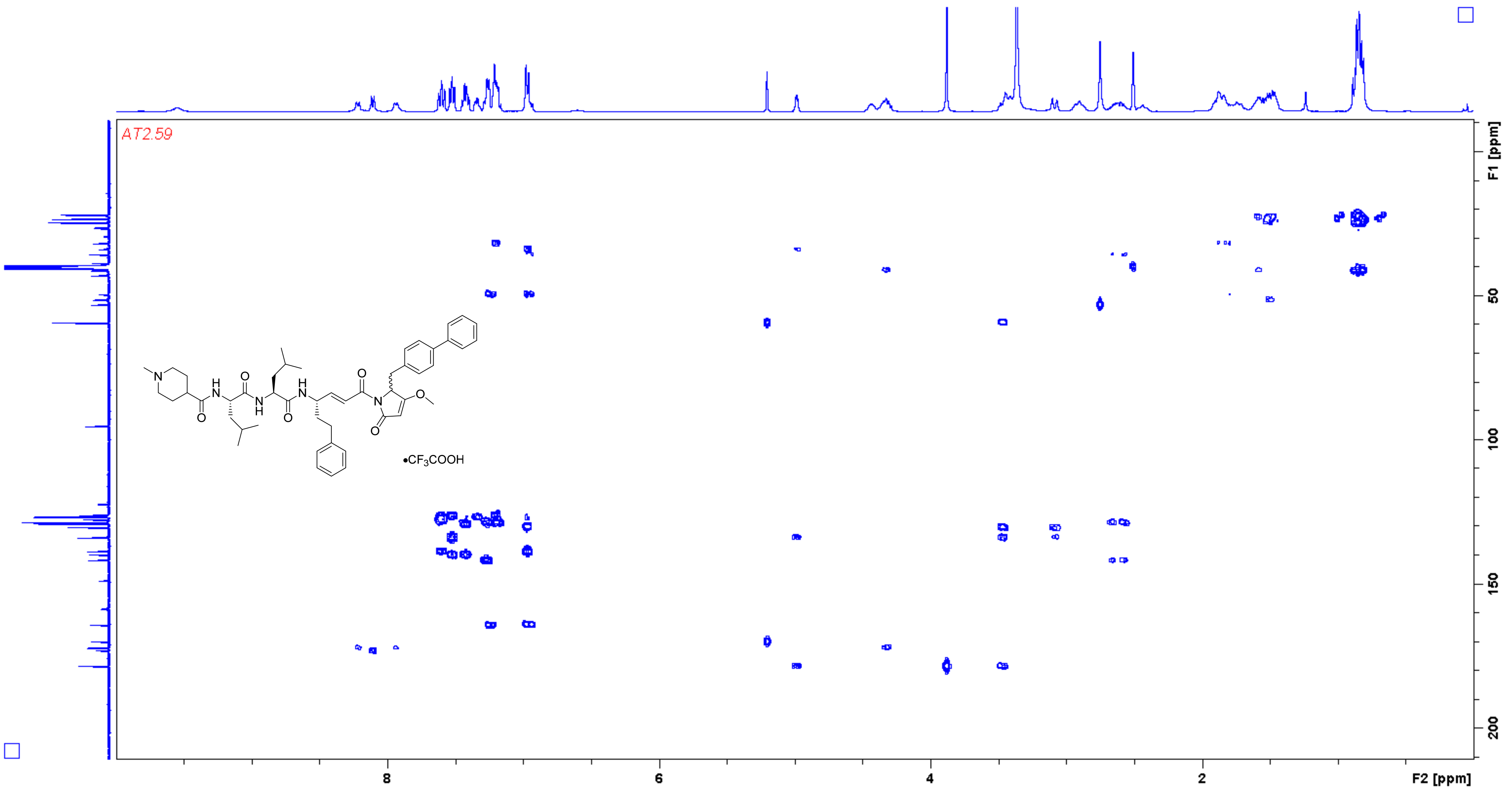

**Analogue 26** $^{1}$H NMR (400 MHz, DMSO-$d_6$)

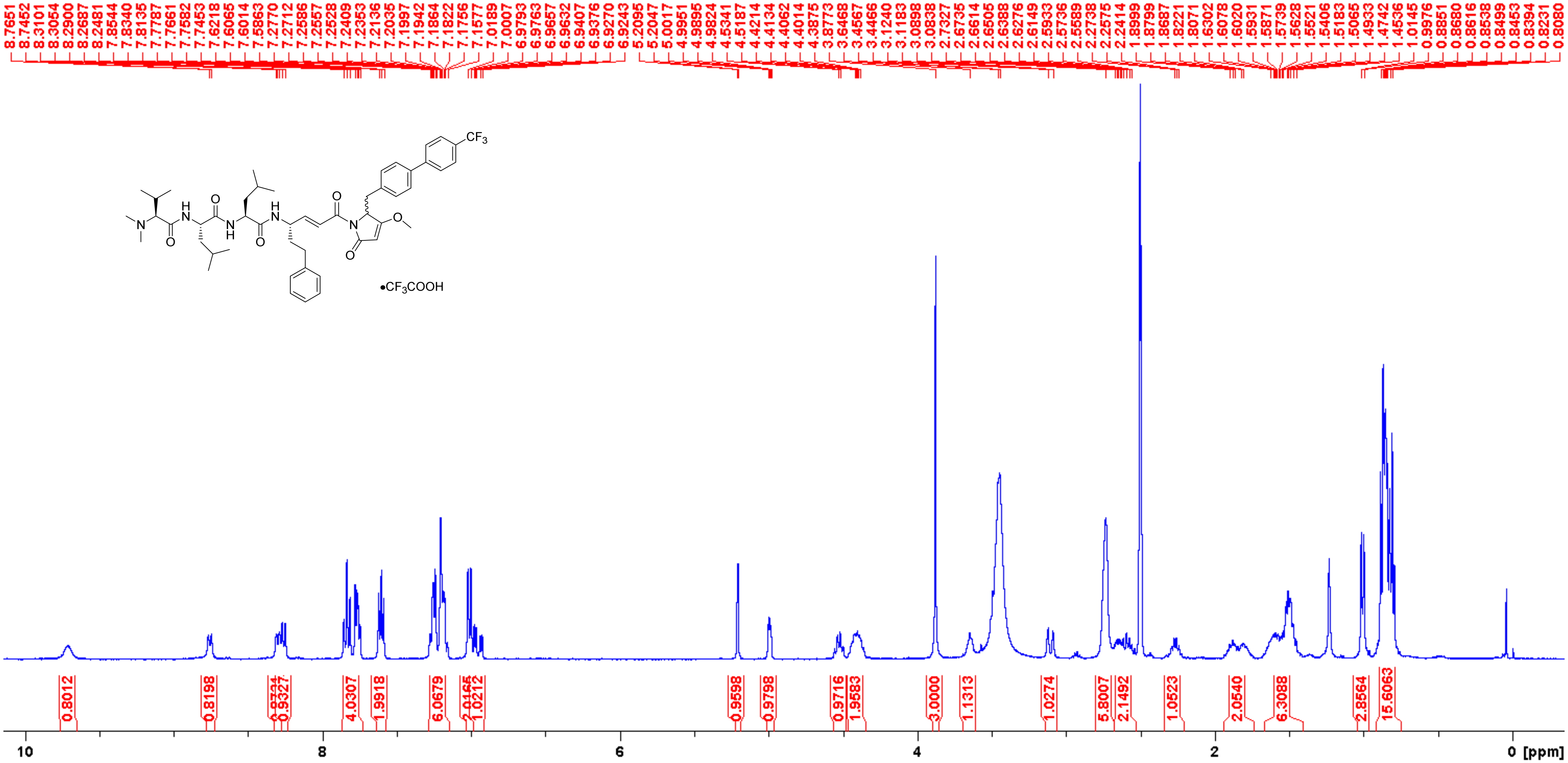

**Analogue 26** $^{13}$C NMR (100 MHz, DMSO-$d_6$)

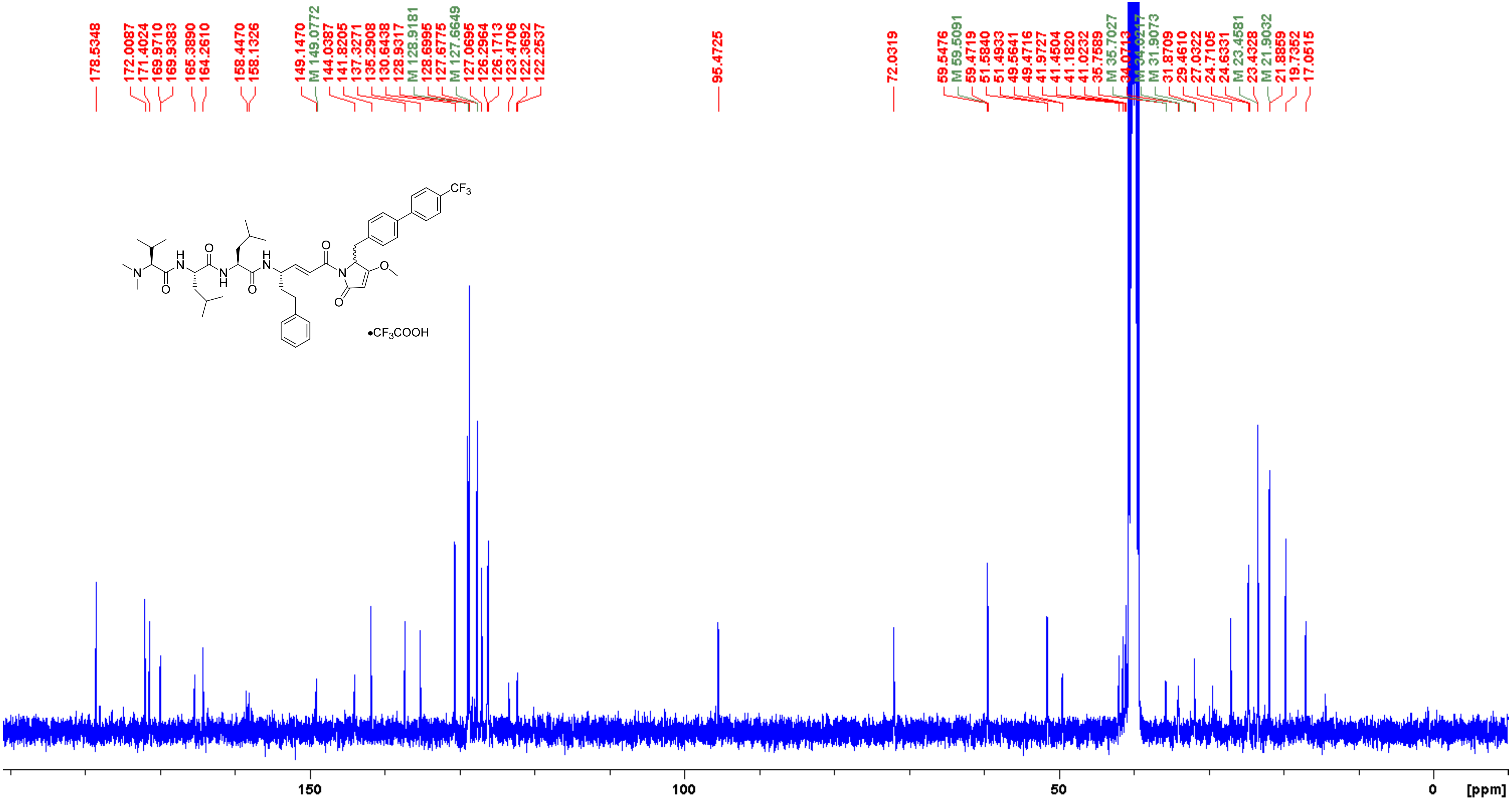

**Analogue 26** COSY (400 MHz, DMSO-$d_6$)

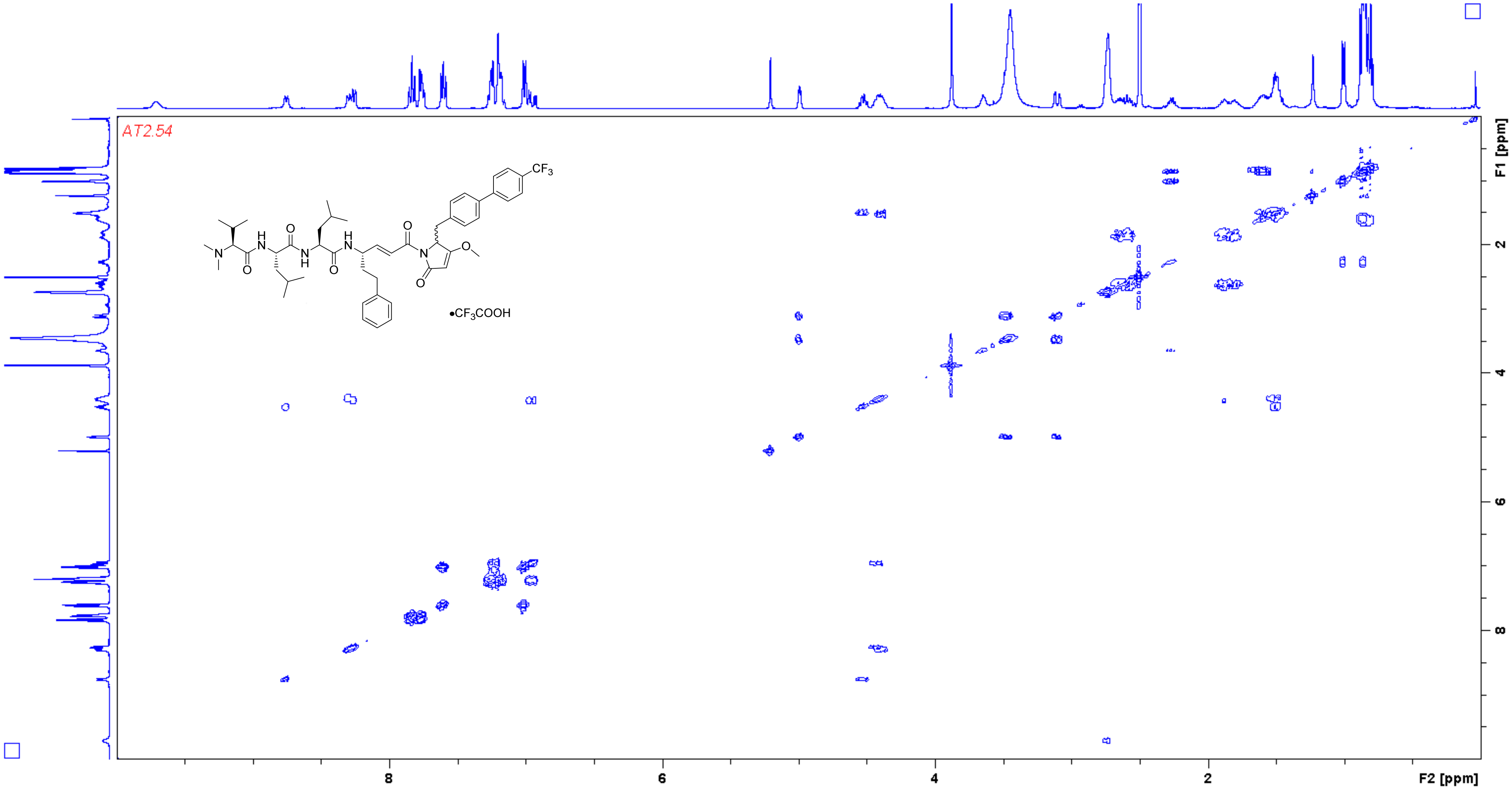

**Analogue 26** HSQC (400/100 MHz, DMSO-$d_6$)

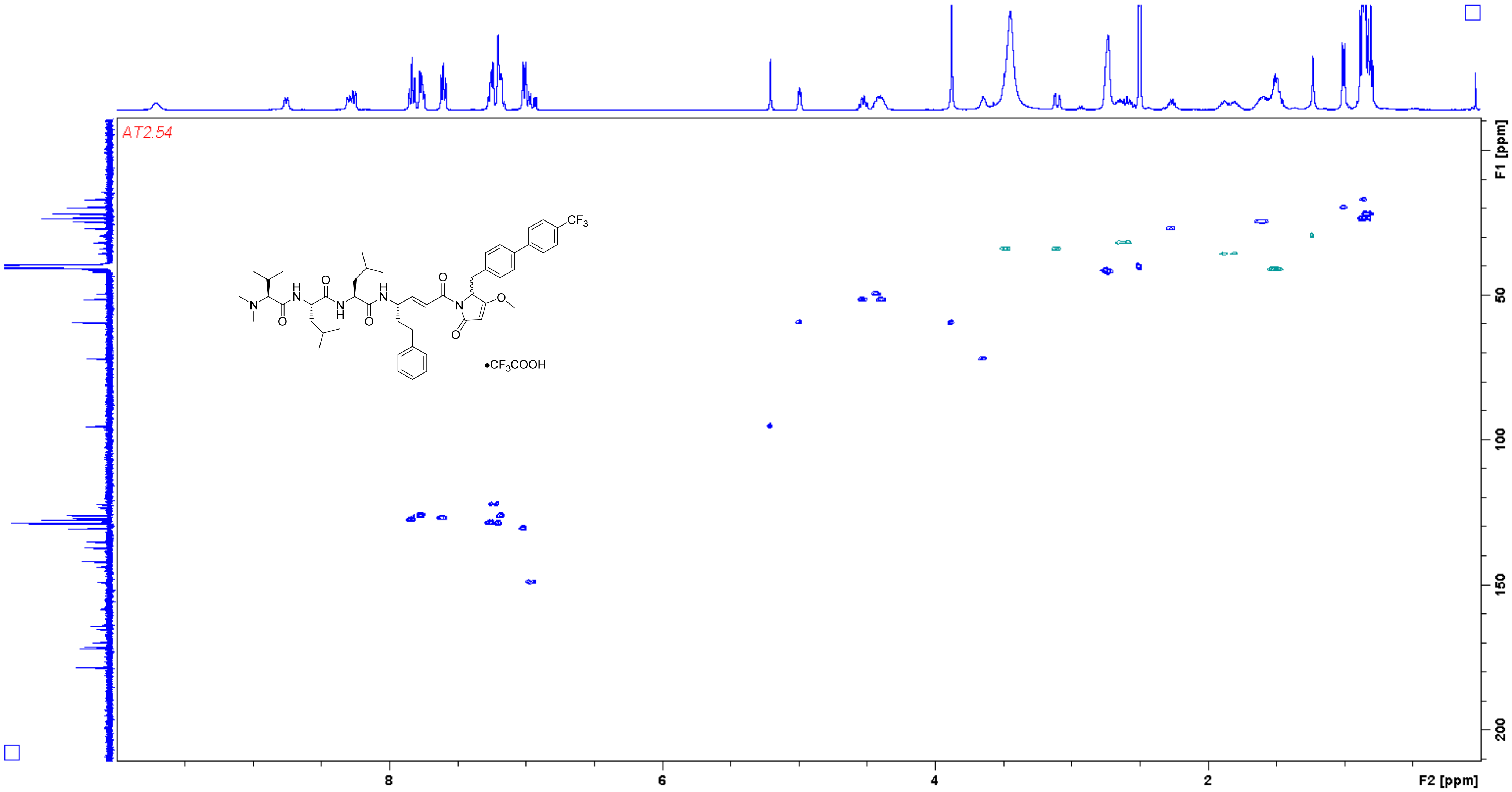

**Analogue 26** HMBC (400/100 MHz, DMSO-$d_6$)

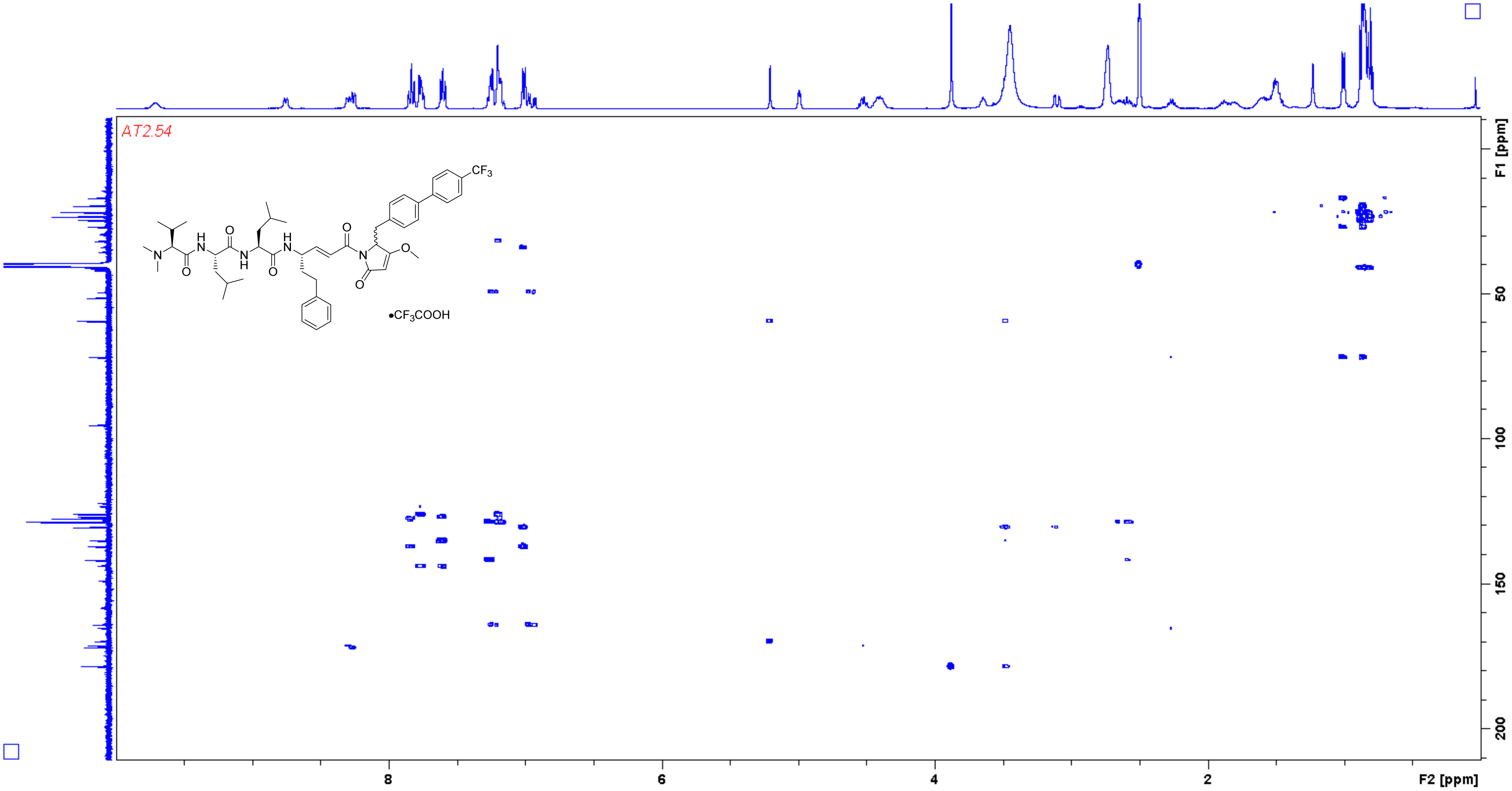

**Analogue 27** [1]H NMR (400 MHz, DMSO-$d_6$)

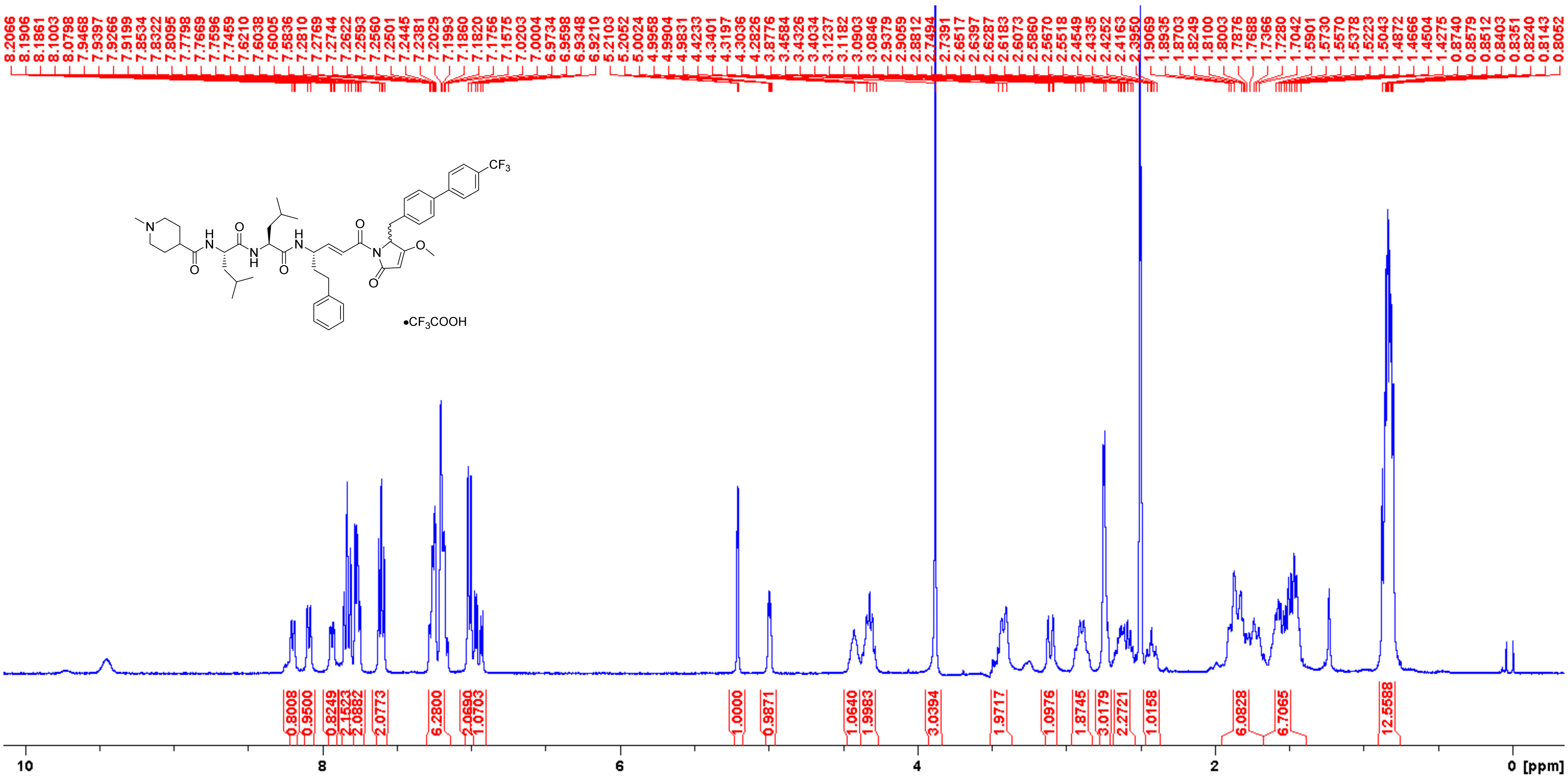

**Analogue 27** $^{13}$C NMR (100 MHz, DMSO-$d_6$)

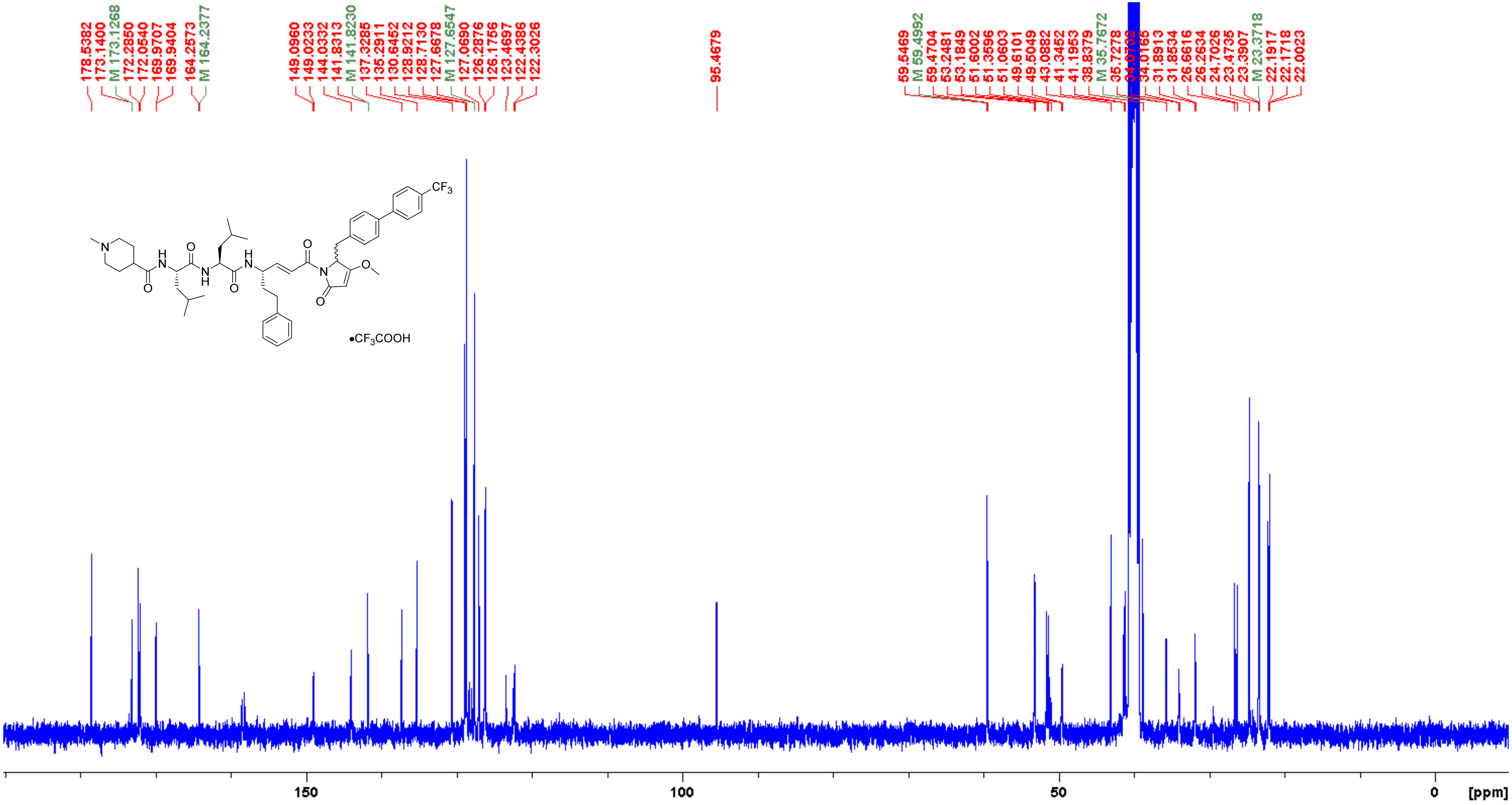

**Analogue 27** COSY (400 MHz, DMSO-$d_6$)

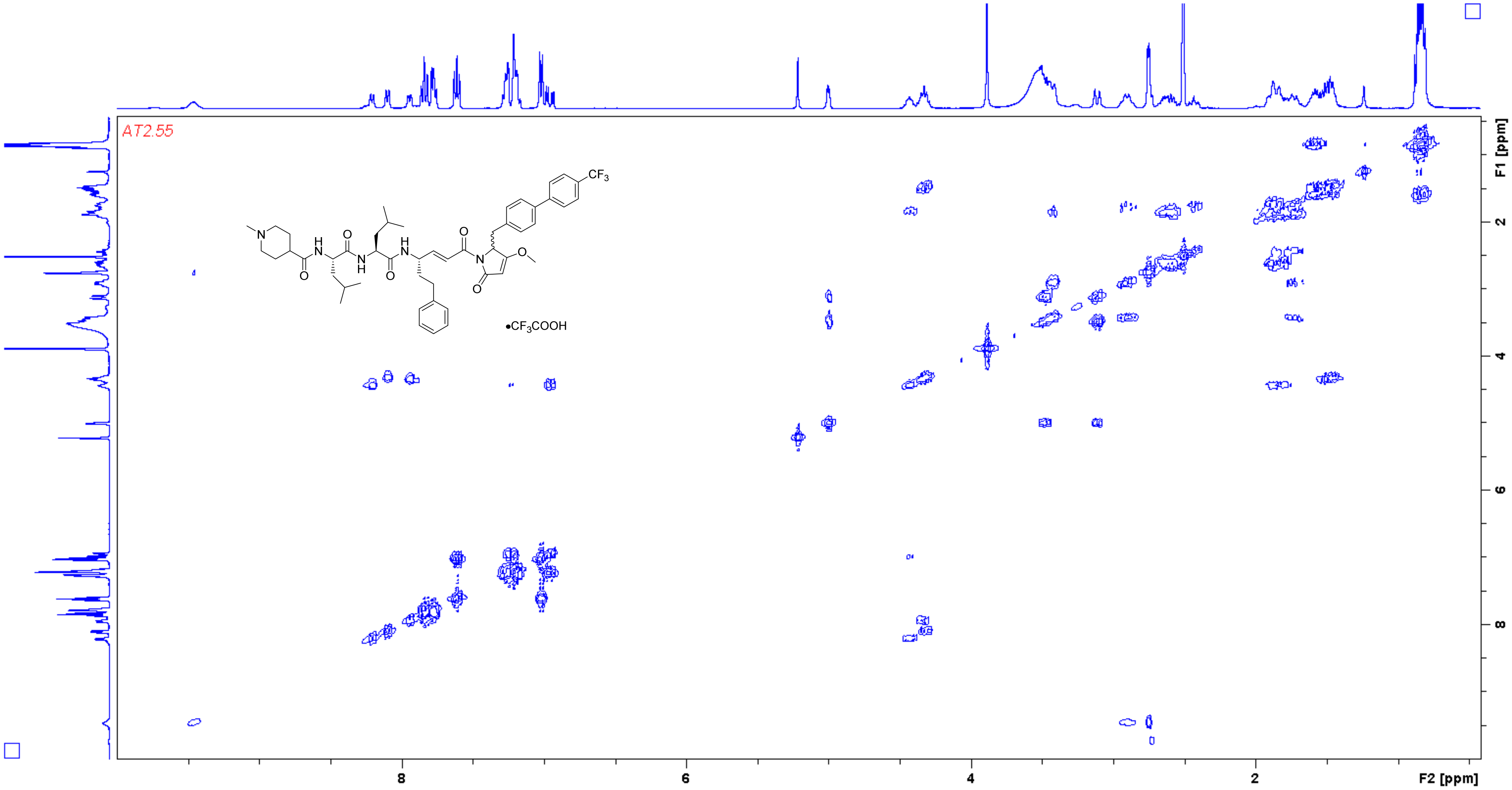

**Analogue 27** HSQC (400/100 MHz, DMSO-$d_6$)

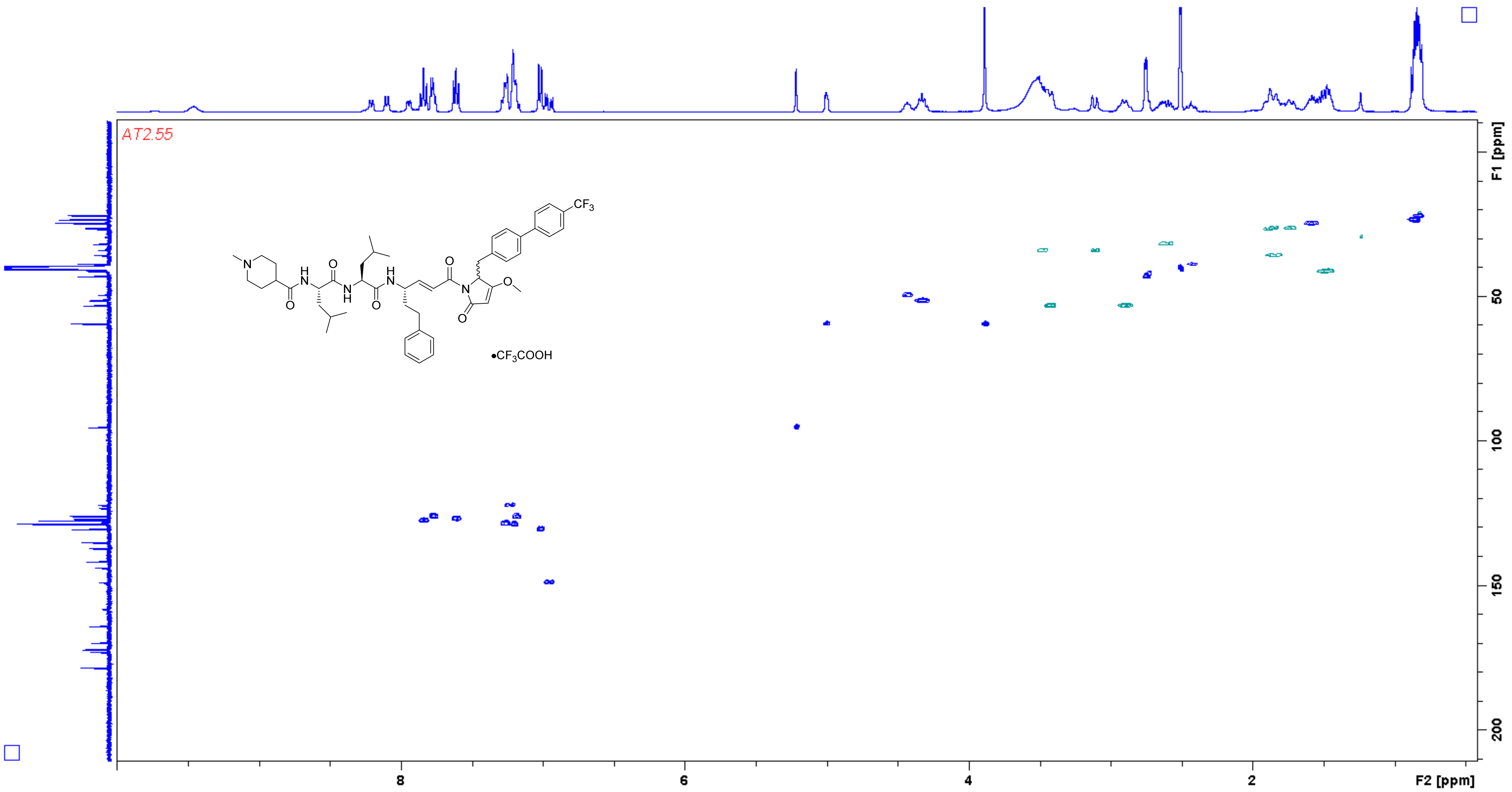

**Analogue 27** HMBC (400/100 MHz, DMSO-$d_6$)

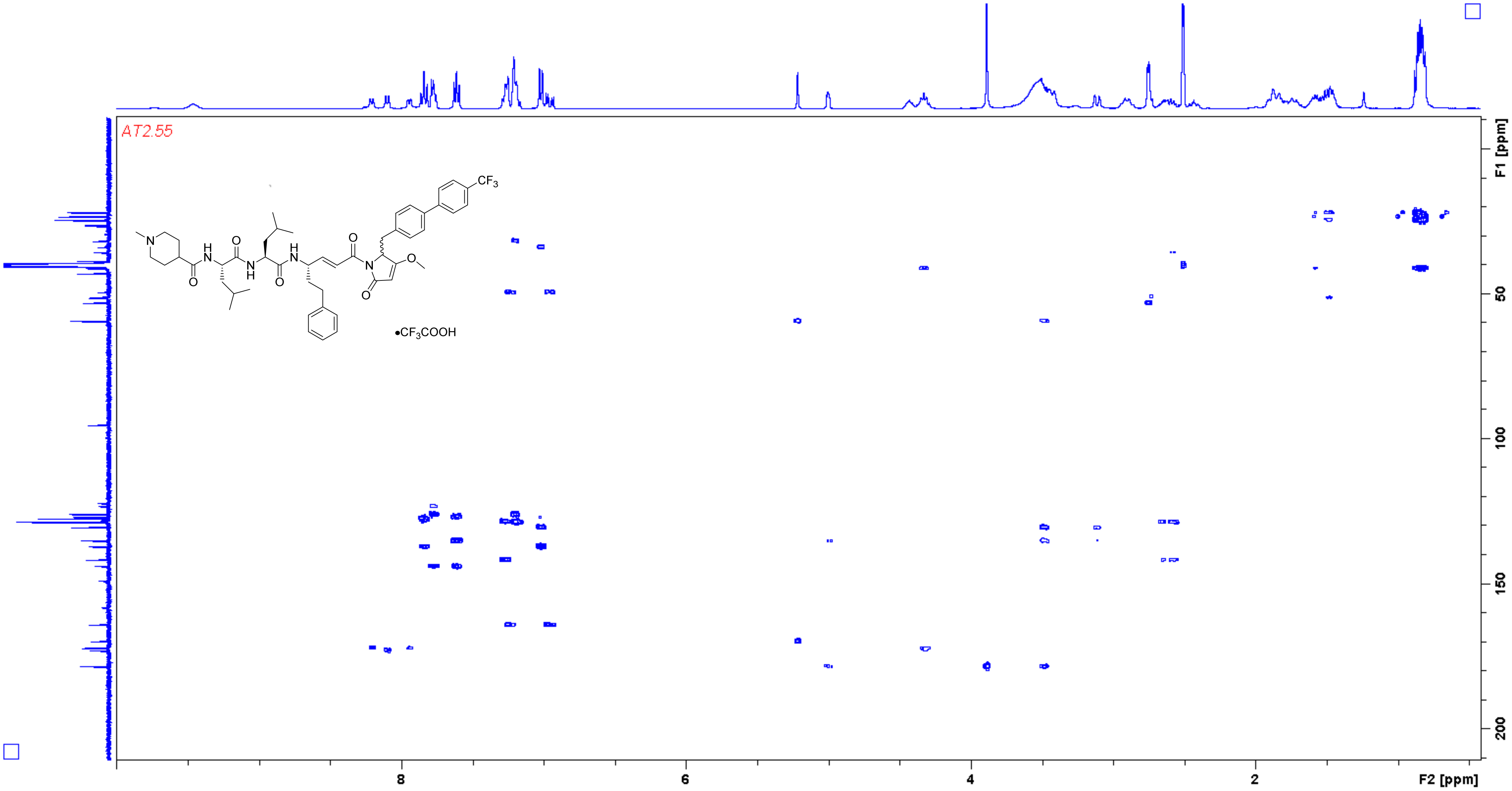

**Analogue 28** [1]H NMR (500 MHz, DMSO-$d_6$)

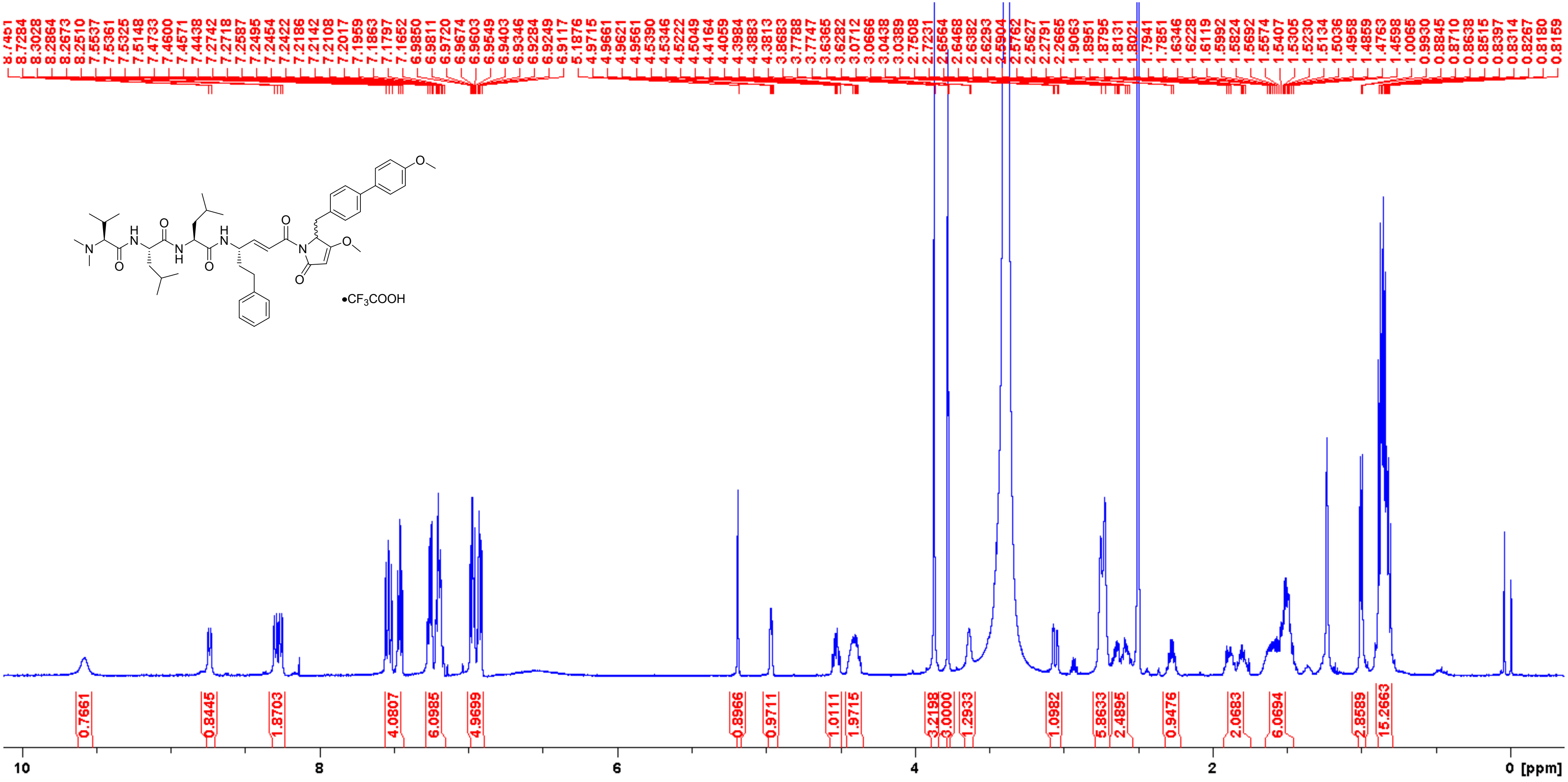

**Analogue 28** $^{13}$C NMR (125 MHz, DMSO-$d_6$)

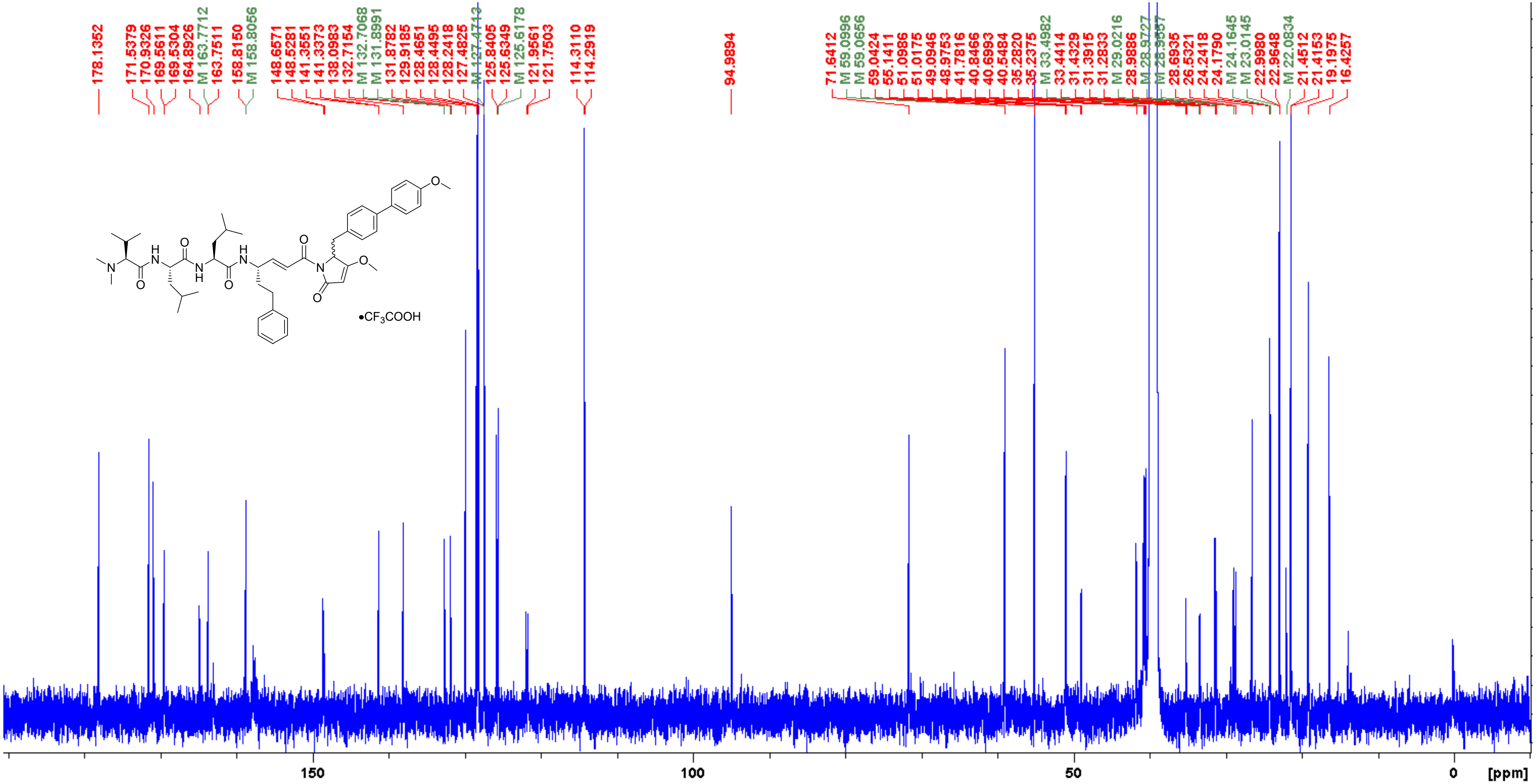

**Analogue 28** COSY (500 MHz, DMSO-$d_6$)

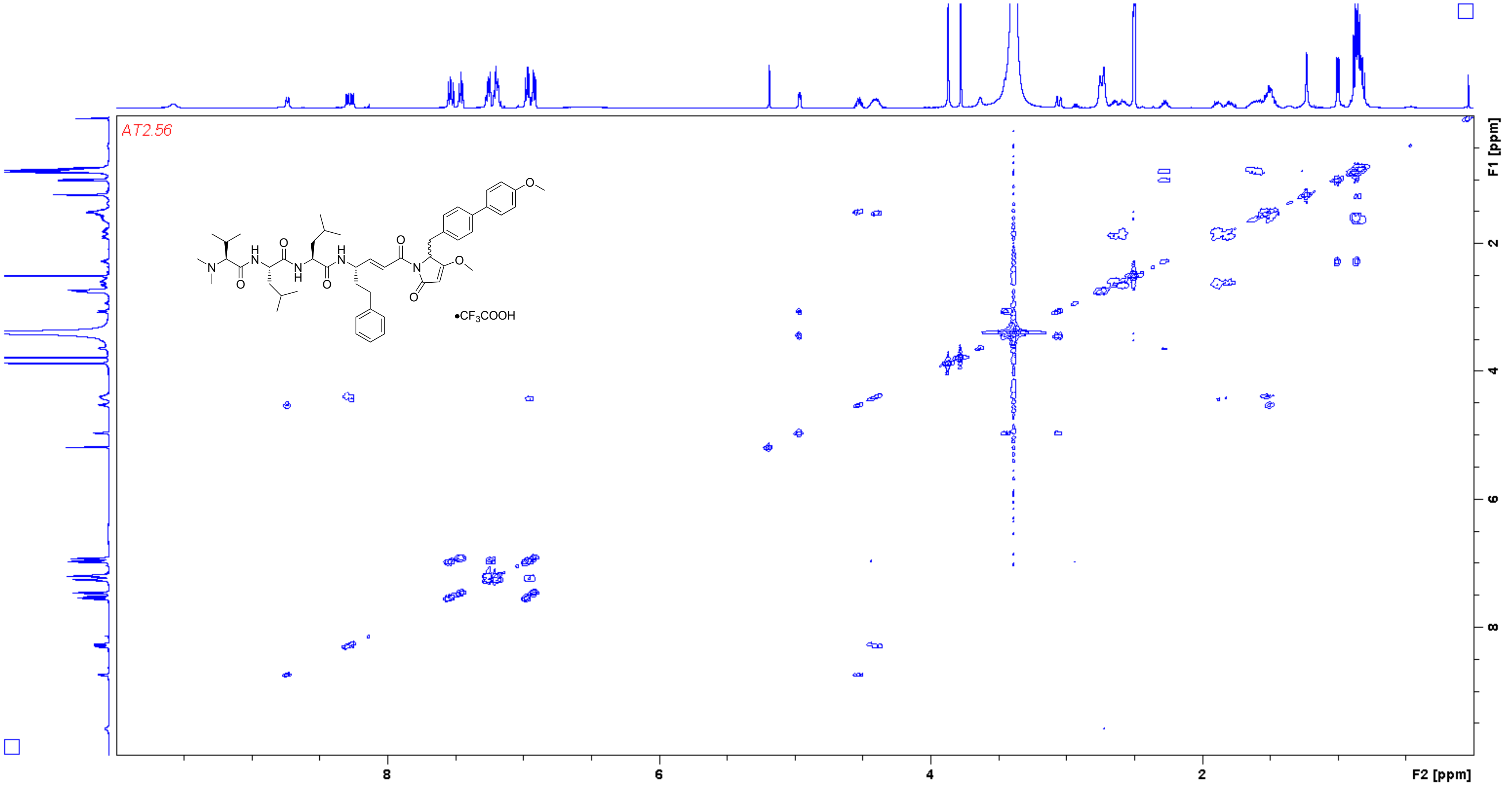

**Analogue 28** HSQC (500/125 MHz, DMSO-$d_6$)

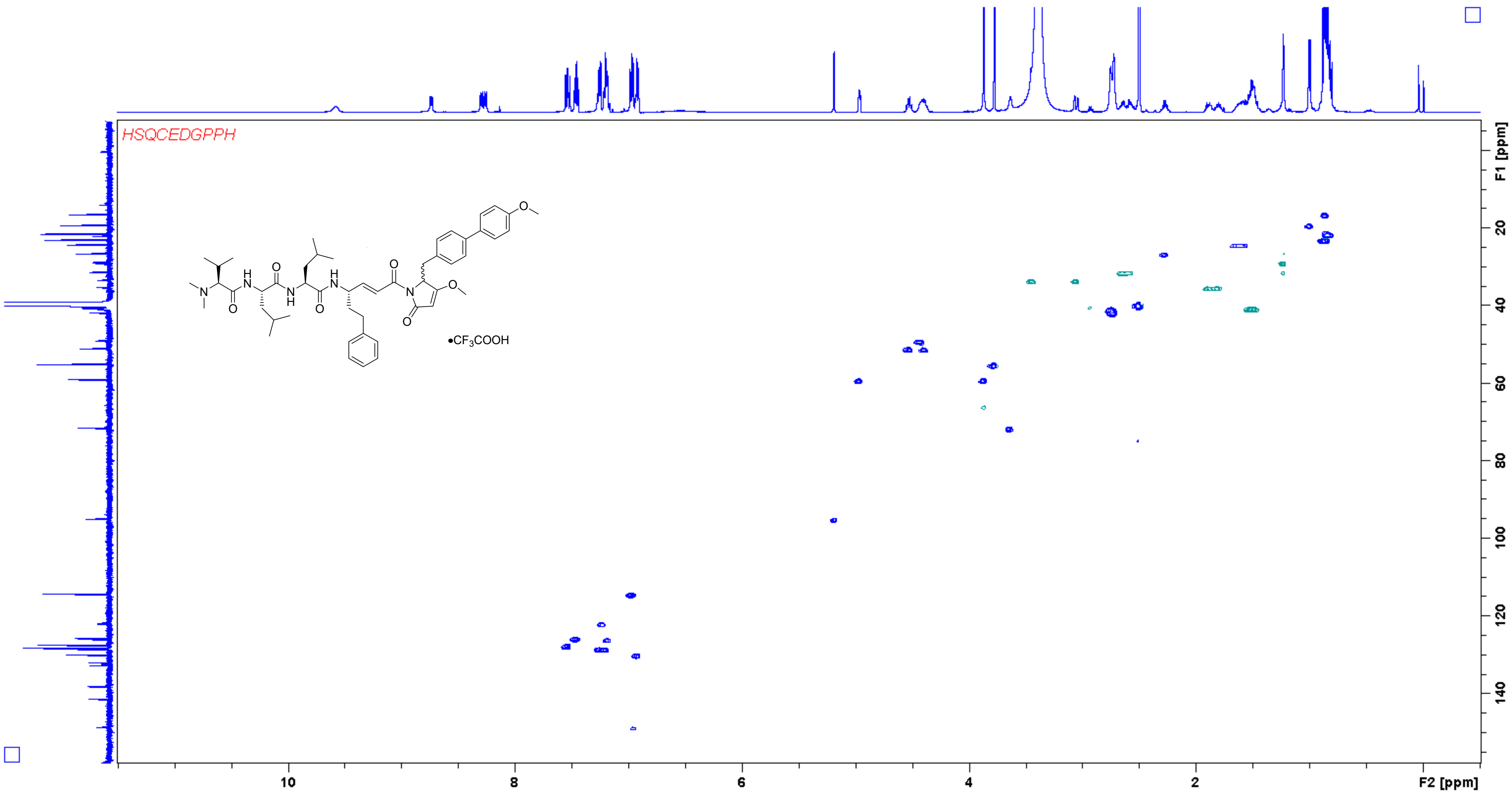

**Analogue 28** HMBC (500/125 MHz, DMSO-$d_6$)

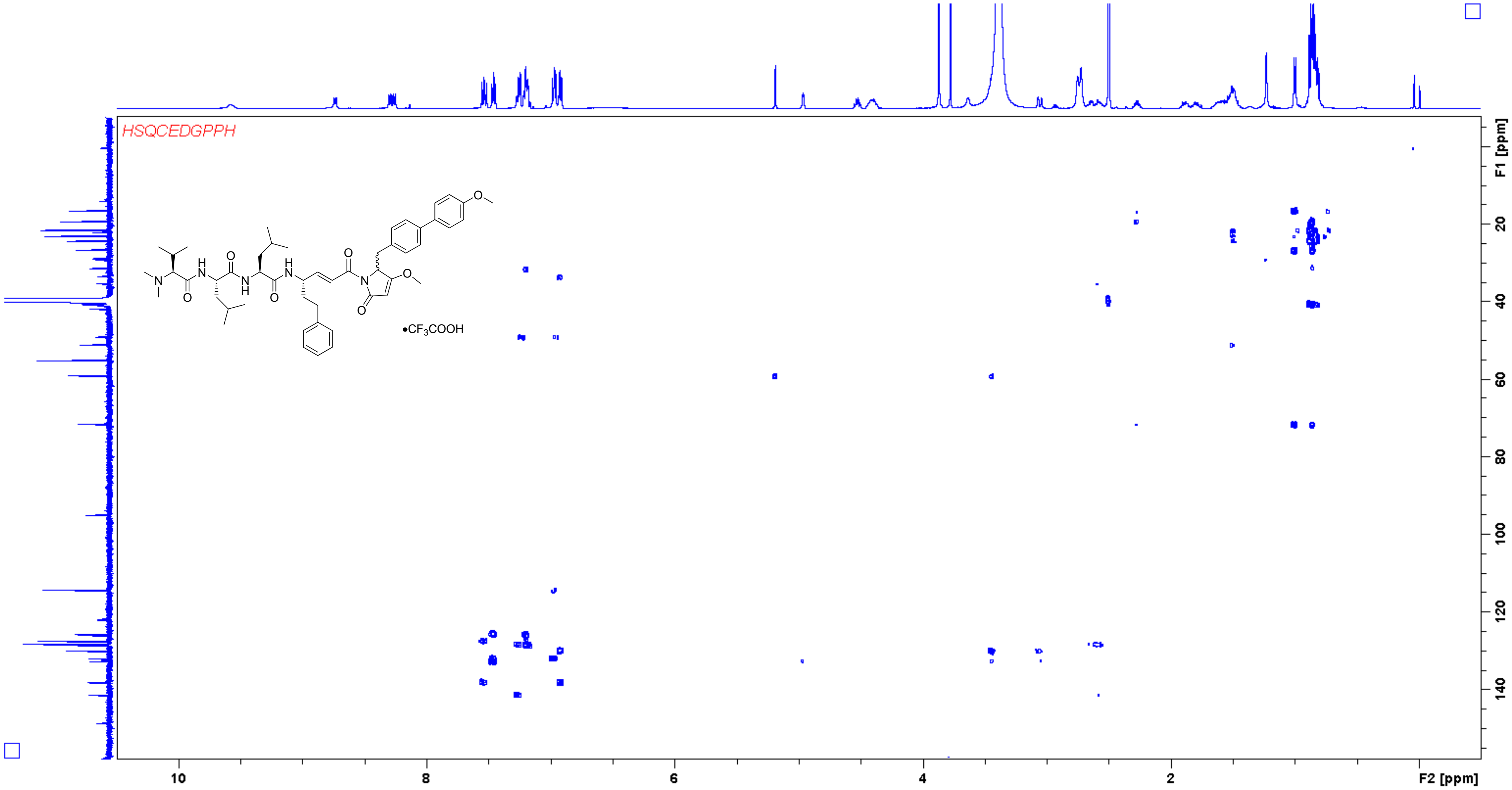

**Analogue 29** $^{1}$H NMR (500 MHz, DMSO-$d_6$)

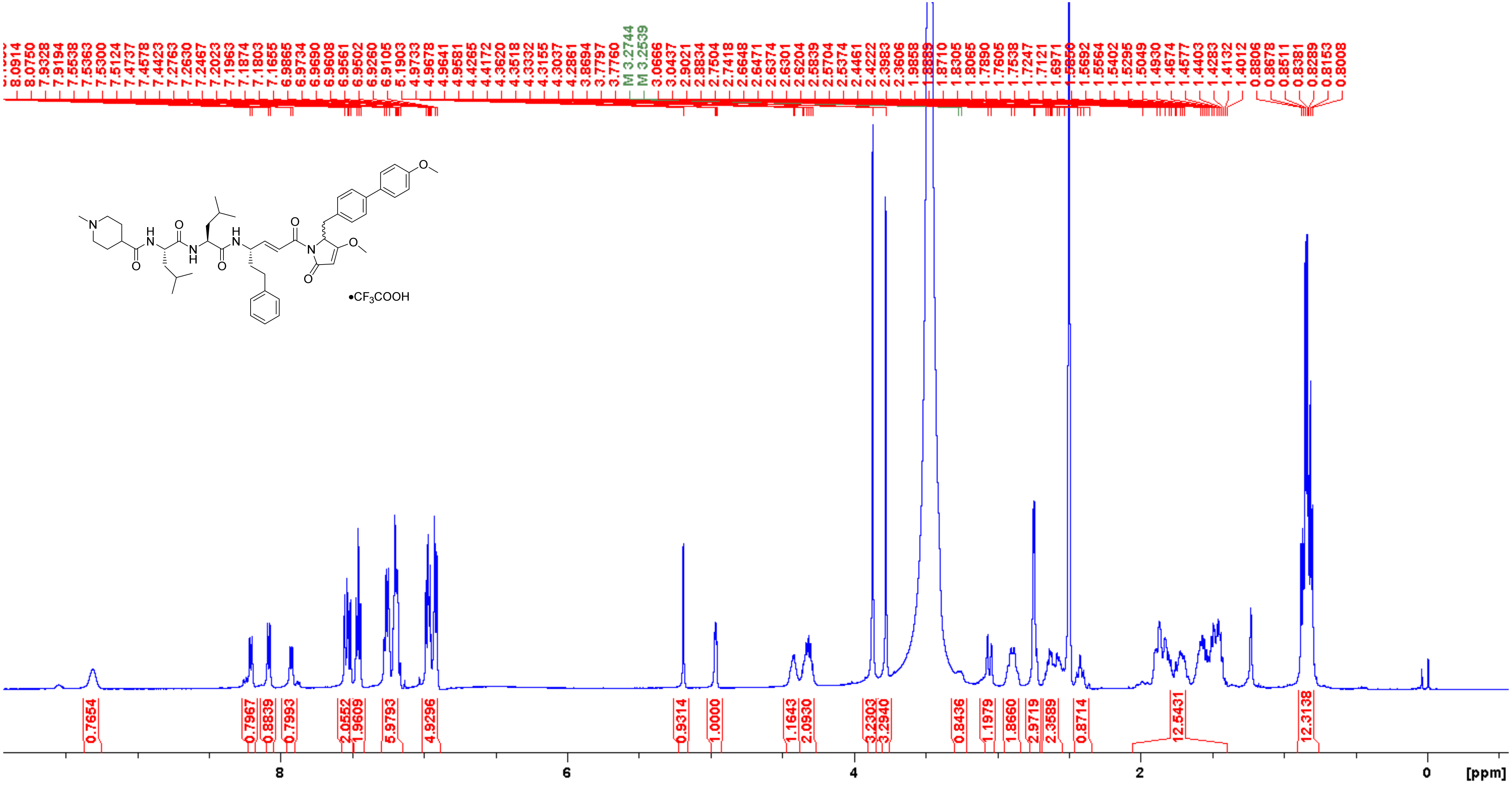

**Analogue 29** $^{13}$C NMR (125 MHz, DMSO-$d_6$)

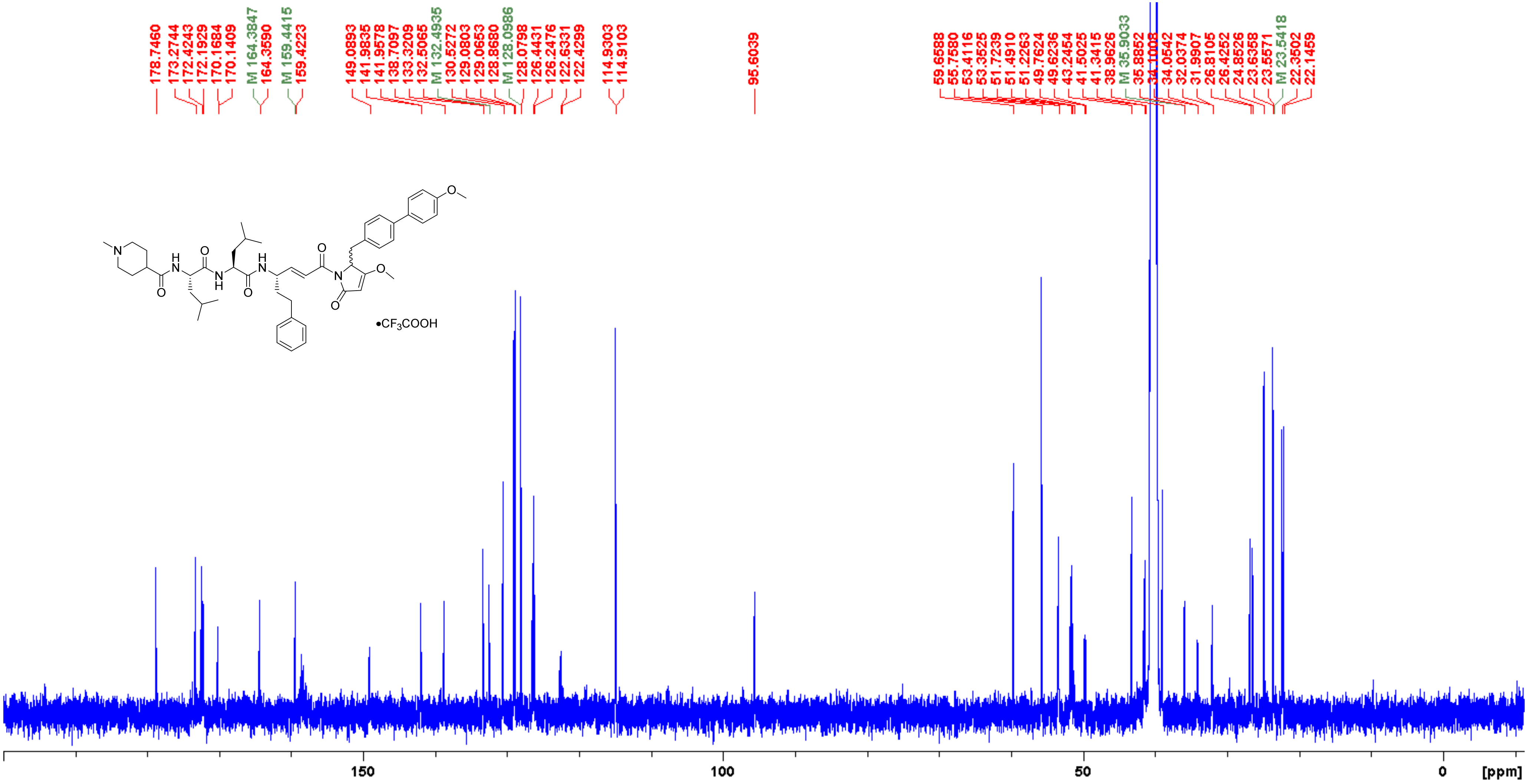

**Analogue 29** COSY (500 MHz, DMSO-$d_6$)

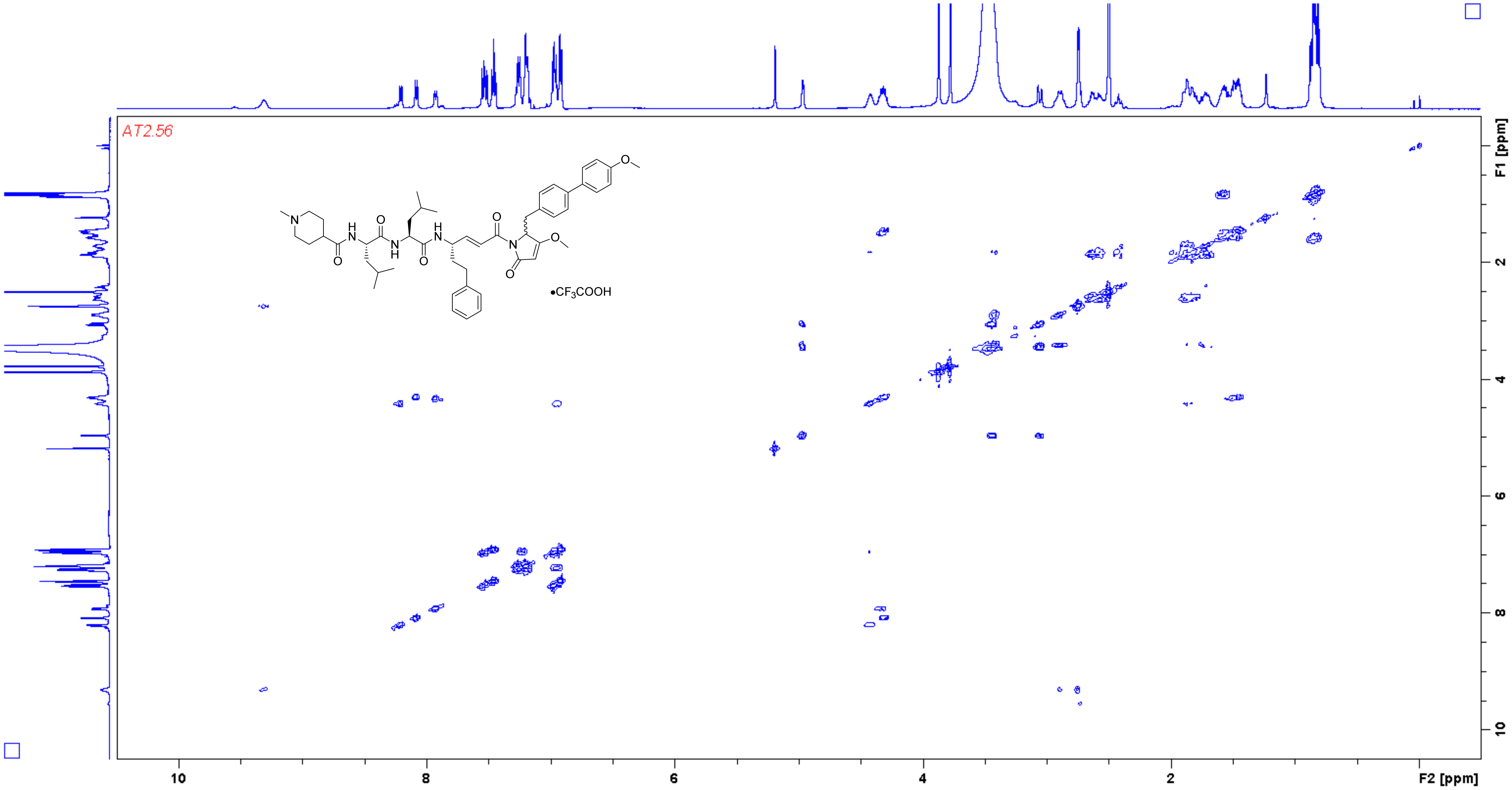

**Analogue 29** HSQC (500/125 MHz, DMSO-$d_6$)

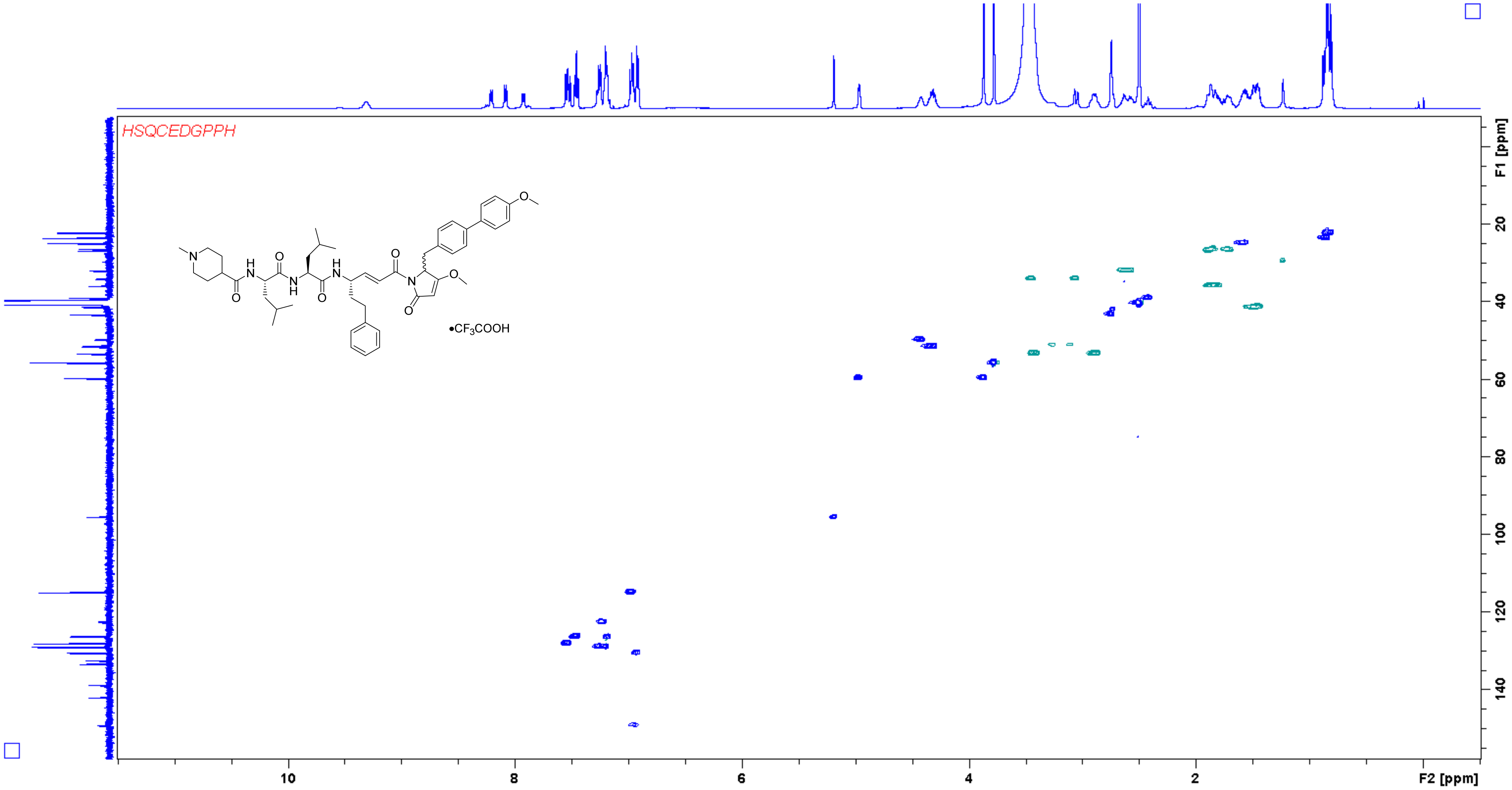

**Analogue 29** HMBC (500/125 MHz, DMSO-$d_6$)

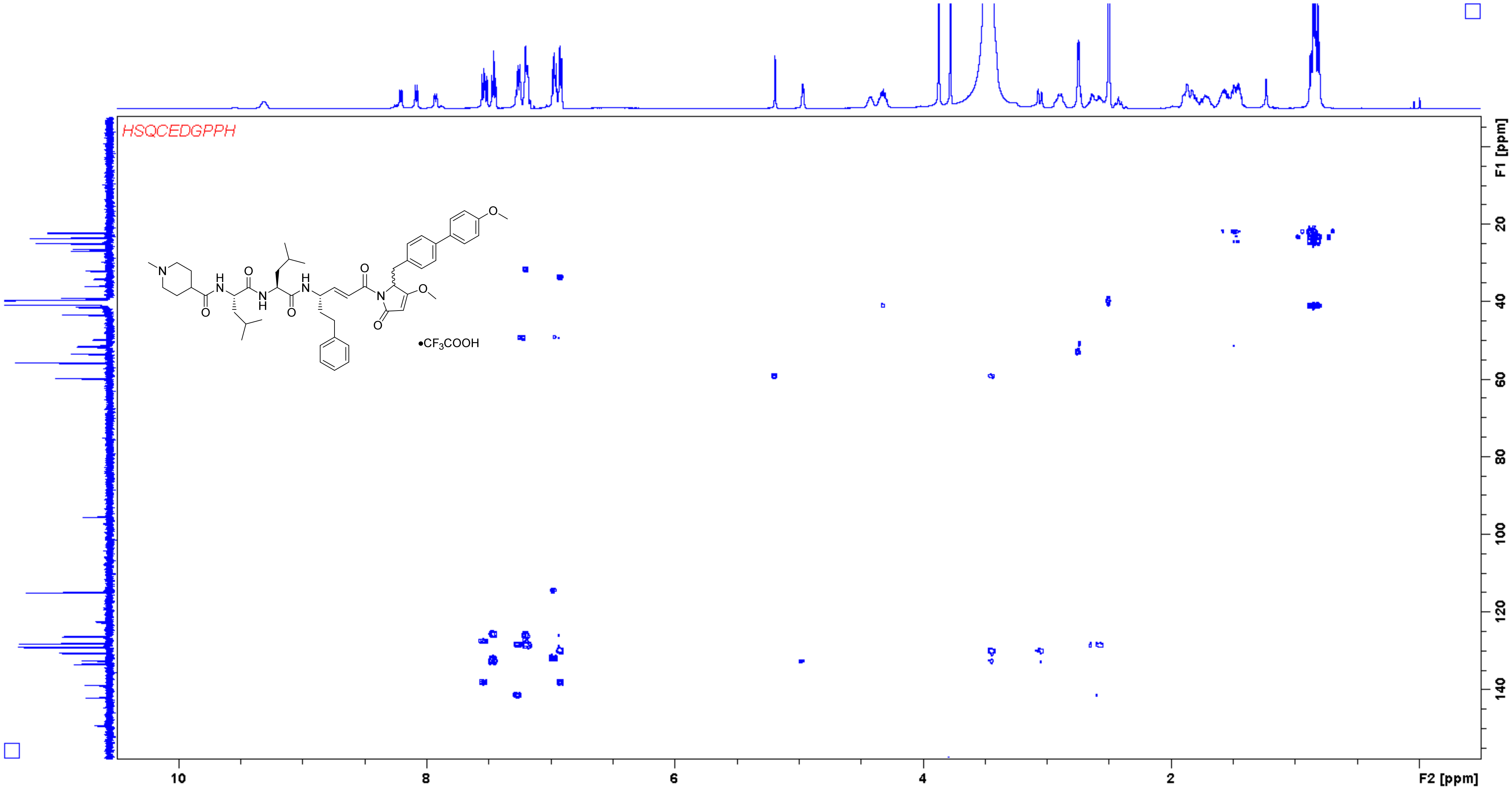

**Analogue 30** [1]H NMR (400 MHz, DMSO-$d_6$)

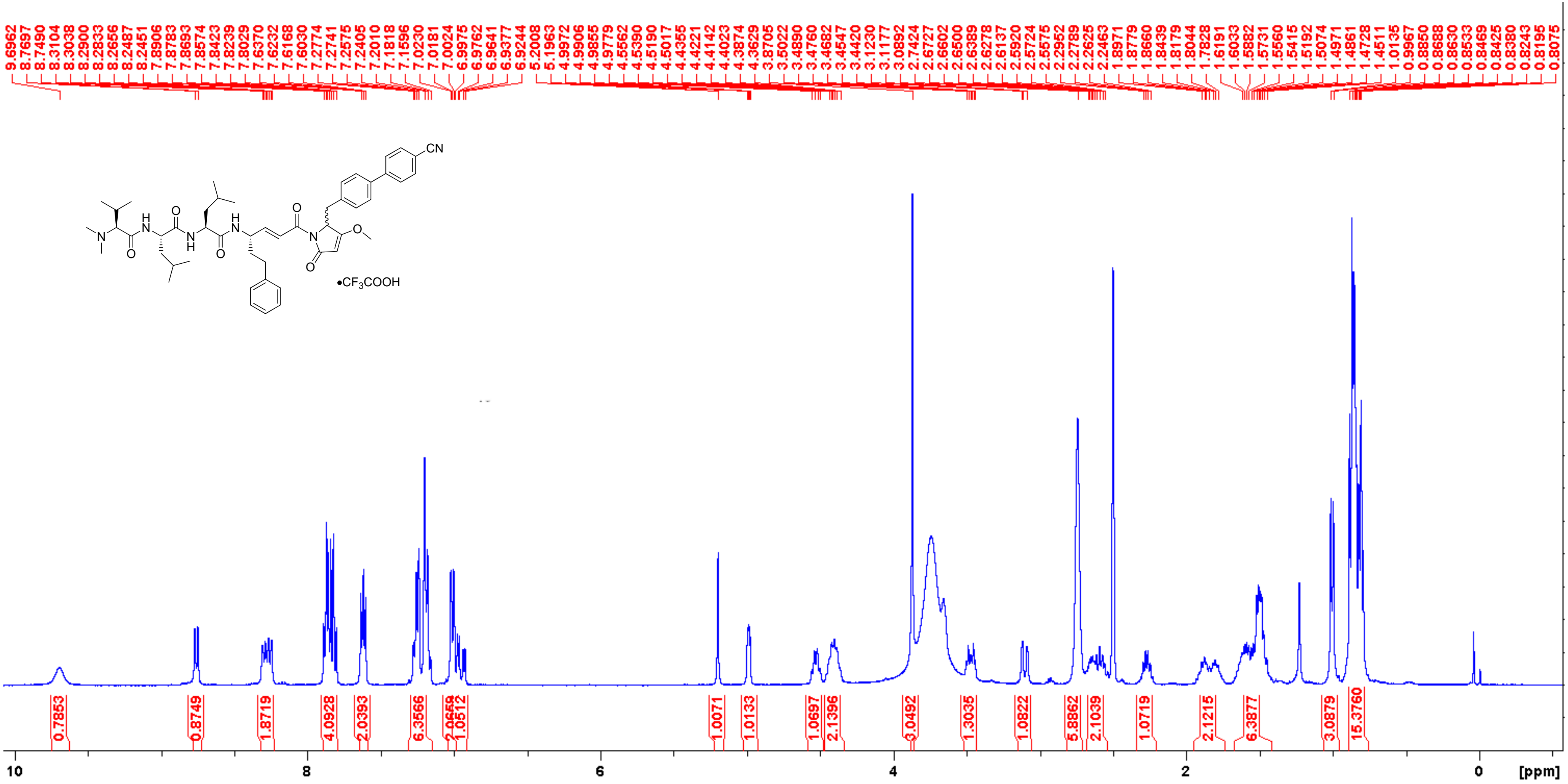

**Analogue 30** $^{13}$C NMR (100 MHz, DMSO-$d_6$)

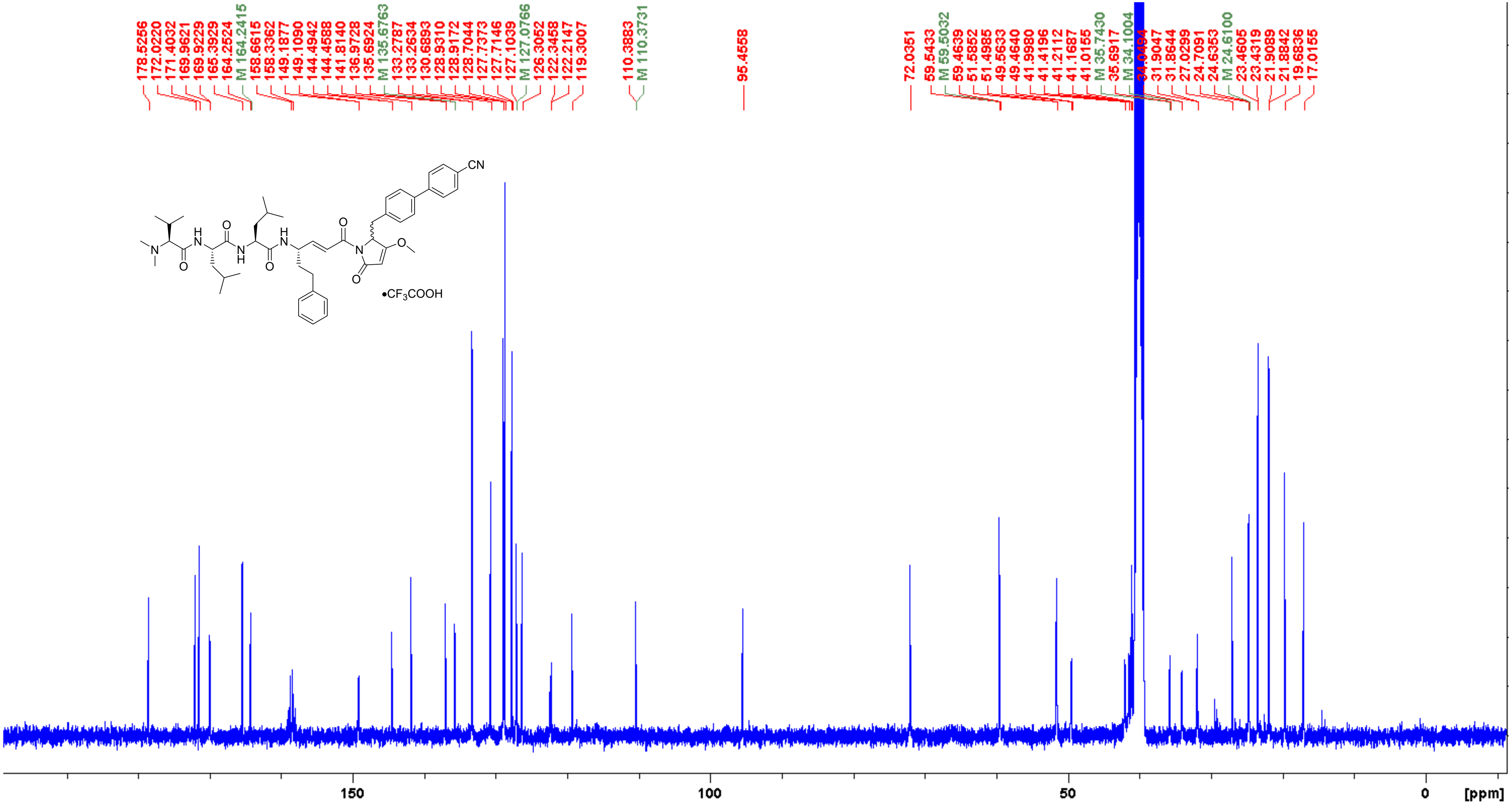

**Analogue 30** COSY (400 MHz, DMSO-$d_6$)

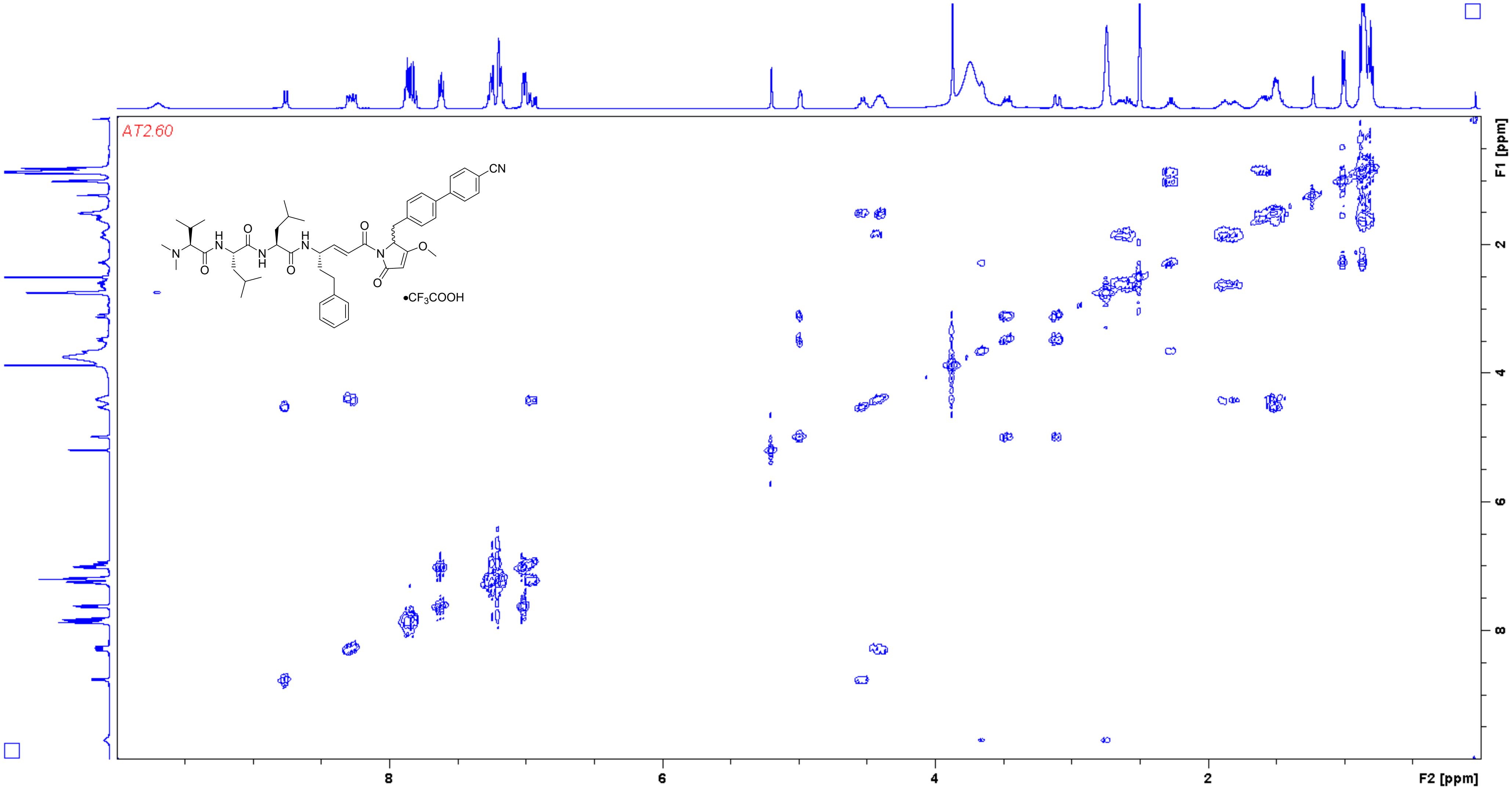

**Analogue 30** HSQC (400/100 MHz, DMSO-$d_6$)

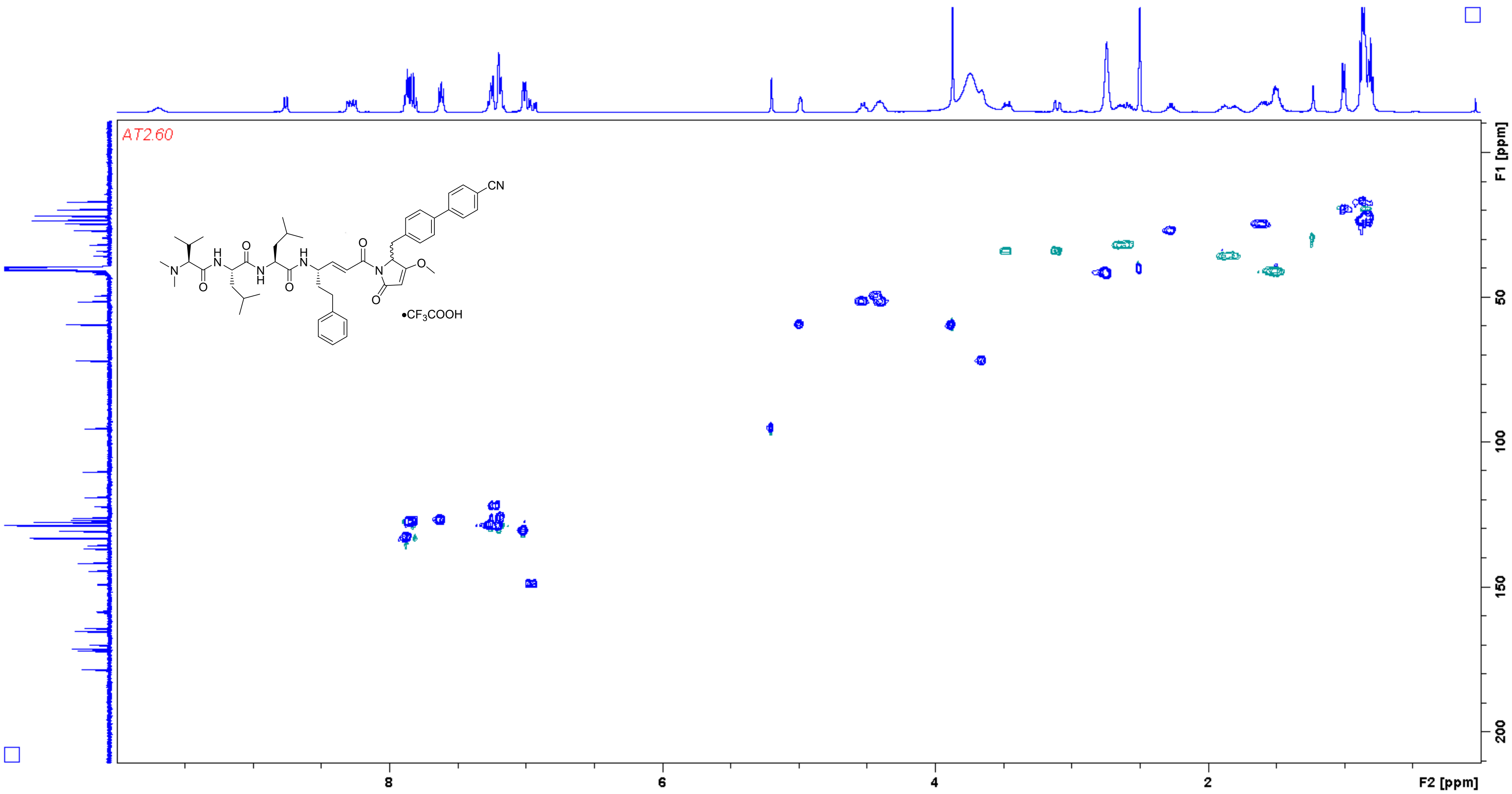

**Analogue 30** HMBC (400/100 MHz, DMSO-$d_6$)

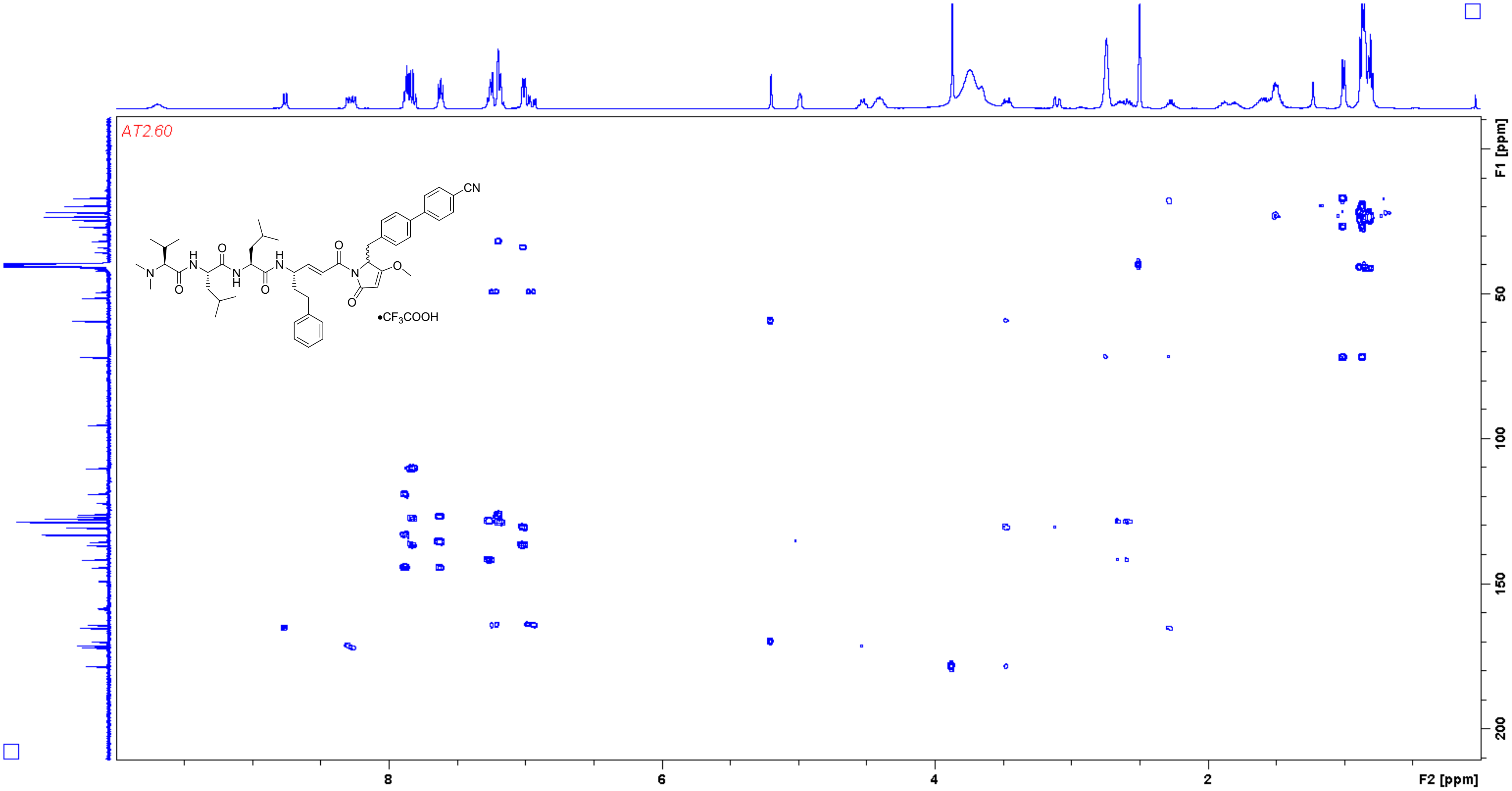

**Analogue 31** $^{1}H$ NMR (400 MHz, DMSO-$d_6$)

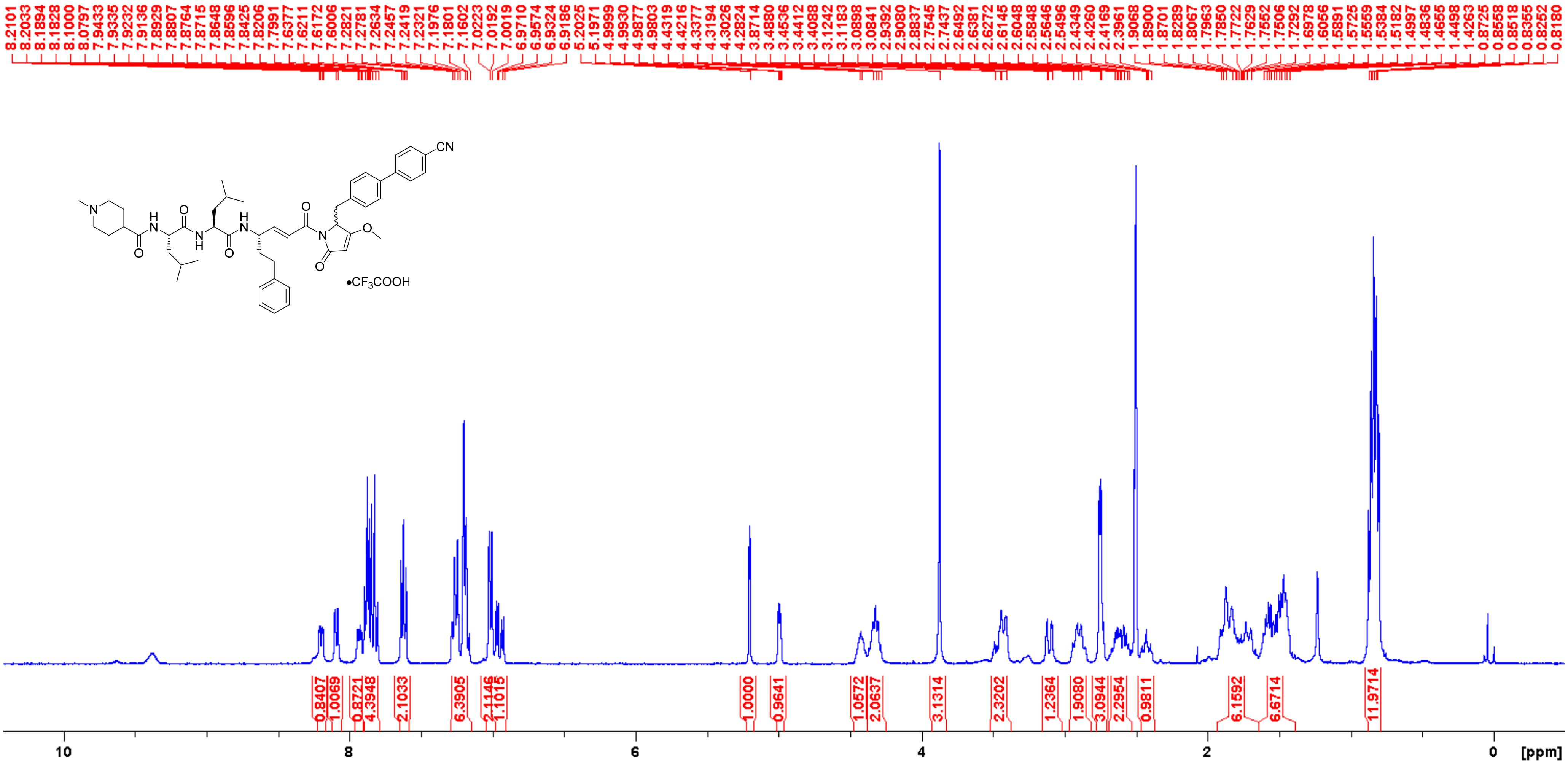

**Analogue 31** $^{13}$C NMR (100 MHz, DMSO-$d_6$)

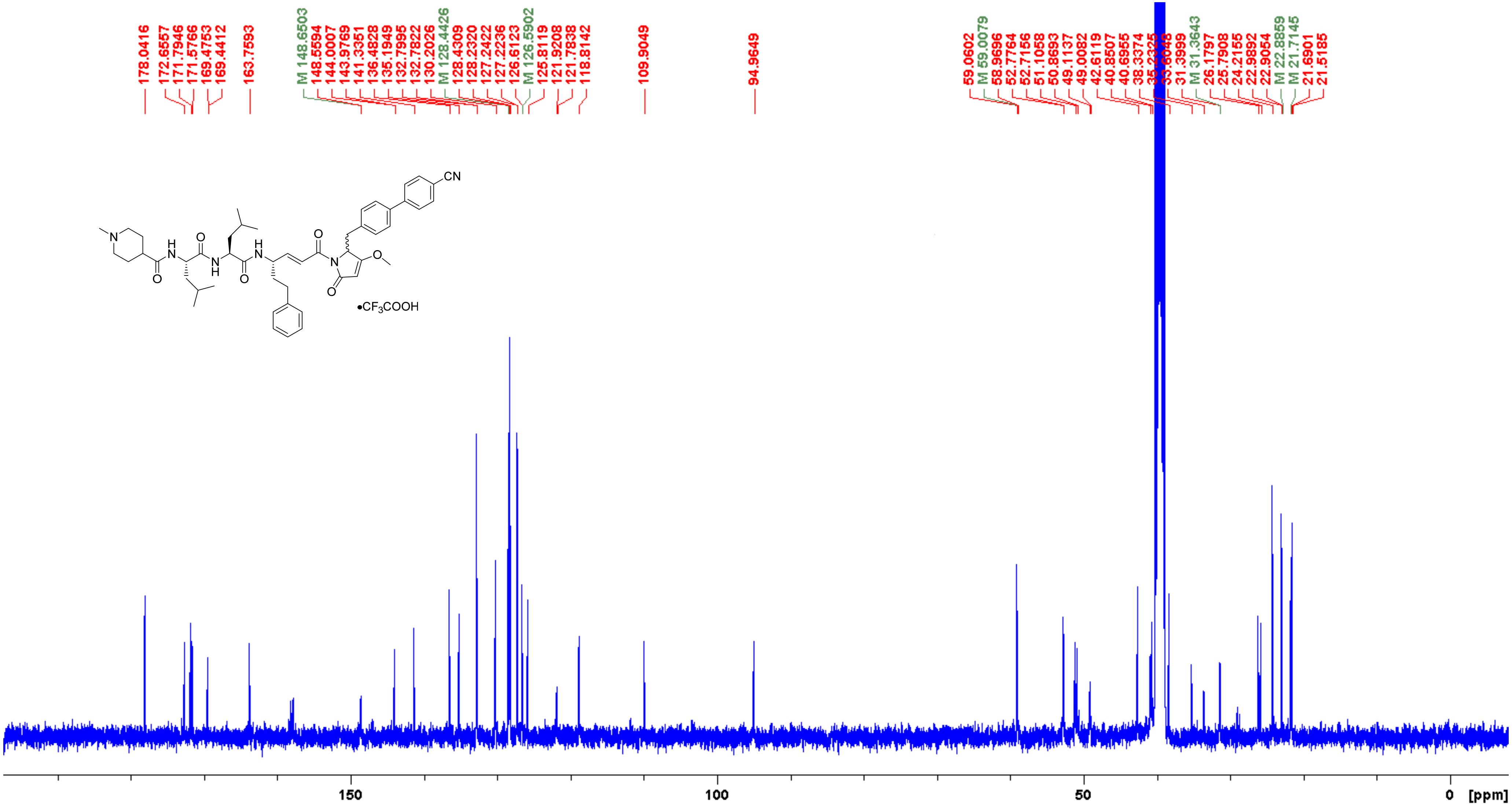

**Analogue 31** COSY (400 MHz, DMSO-$d_6$)

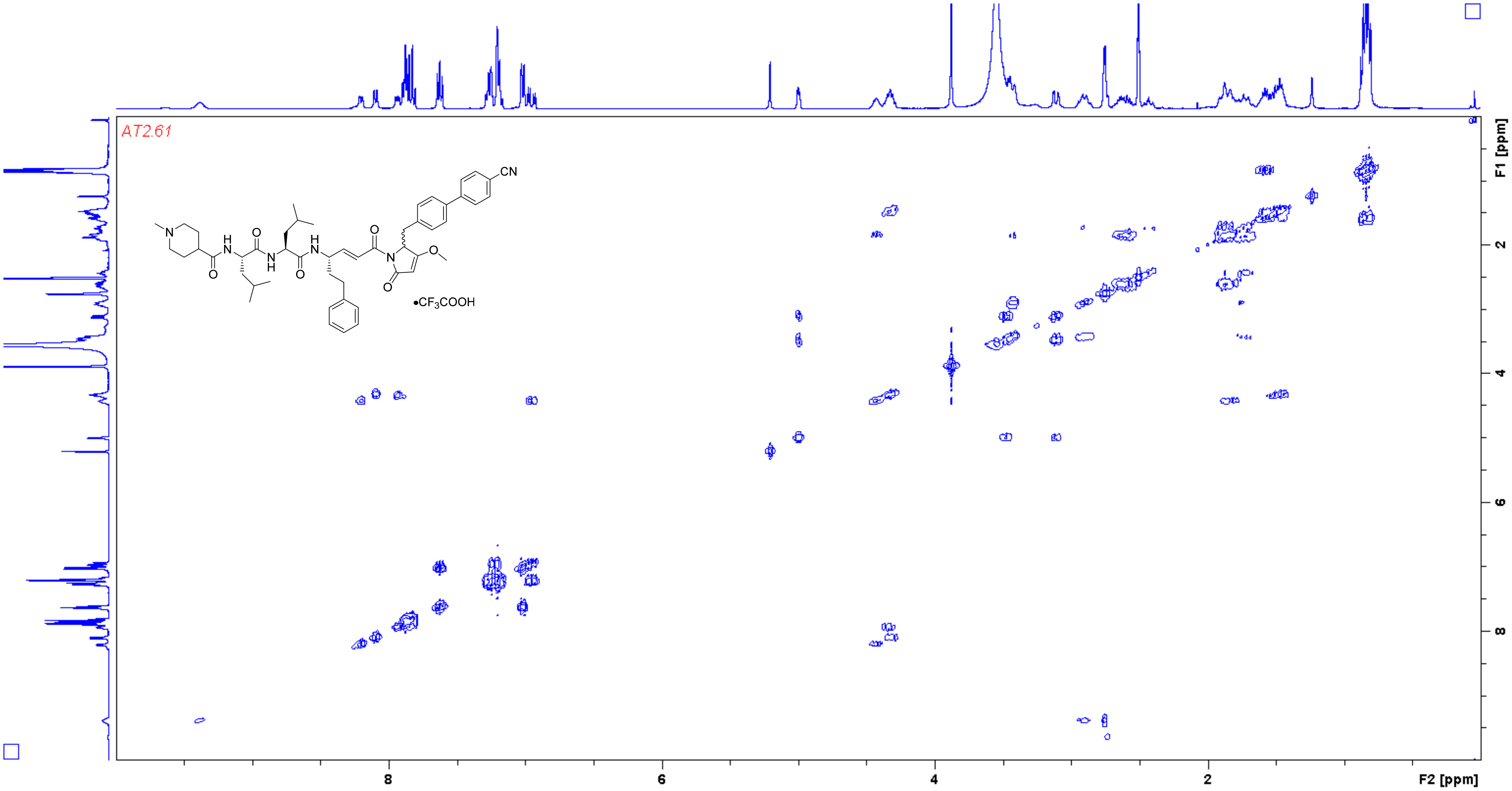

**Analogue 31** HSQC (400/100 MHz, DMSO-$d_6$)

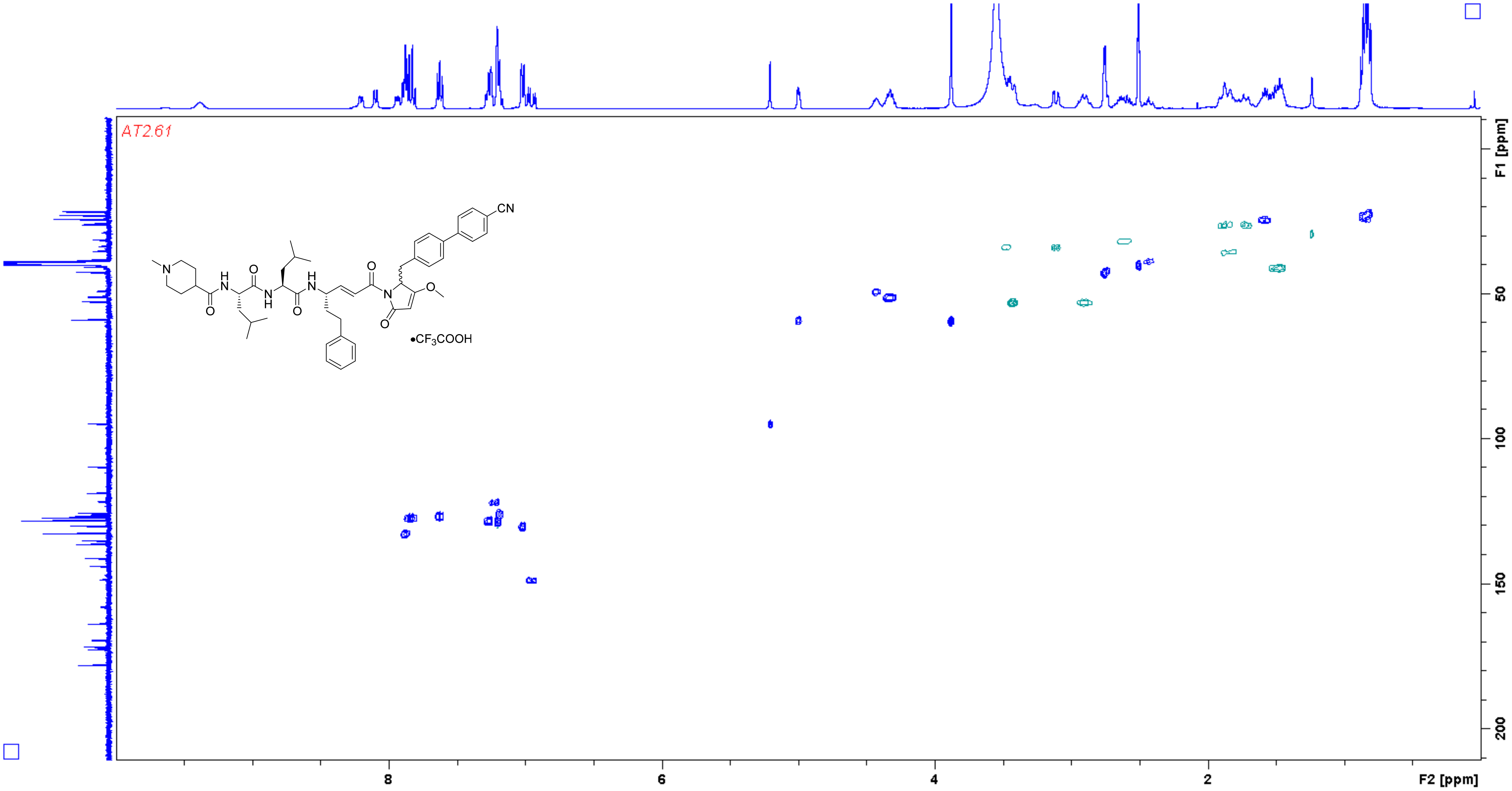

**Analogue 31** HMBC (400/100 MHz, DMSO-$d_6$)

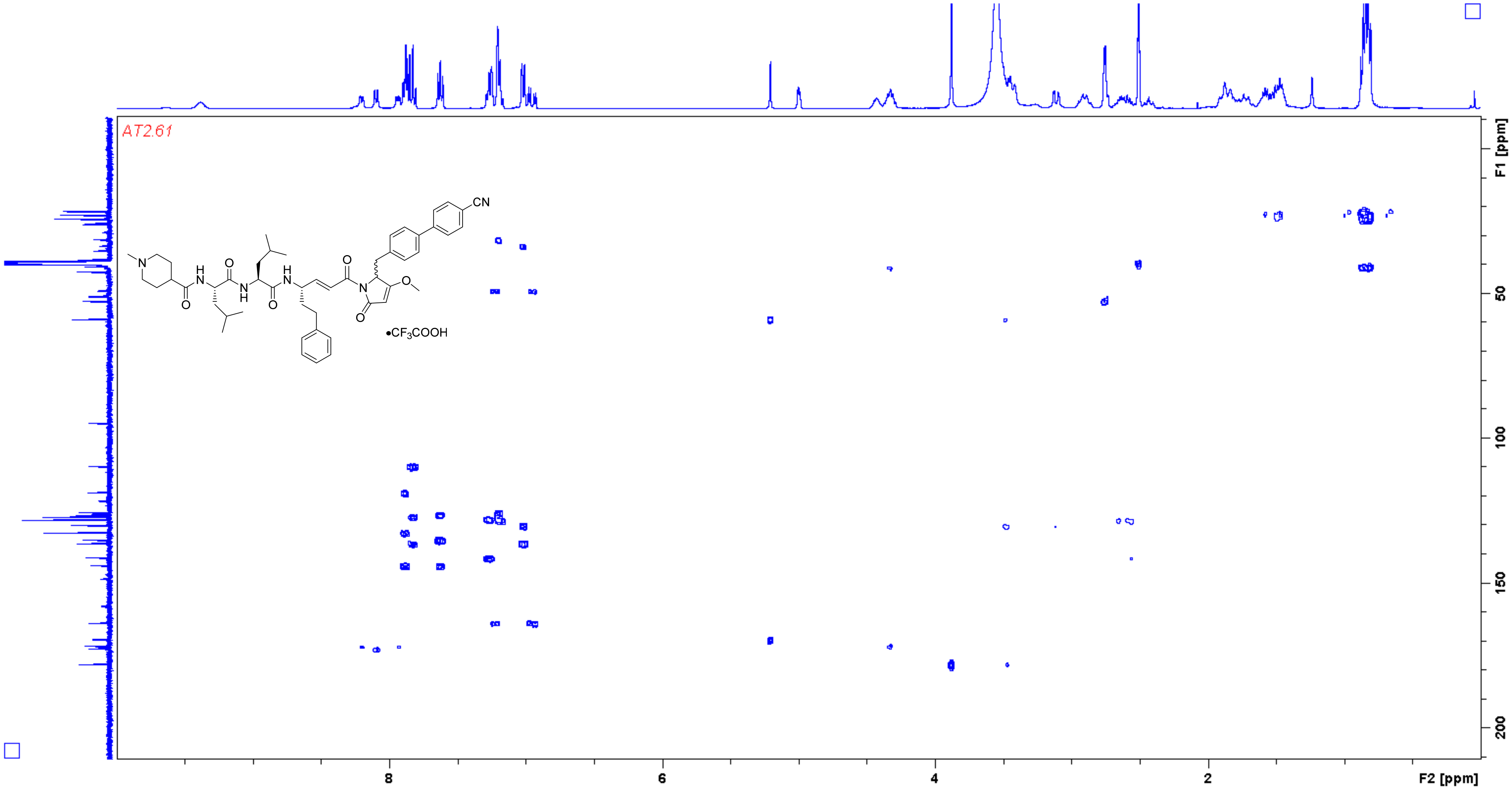

**Analogue 32** $^1$H NMR (400 MHz, DMSO-$d_6$)

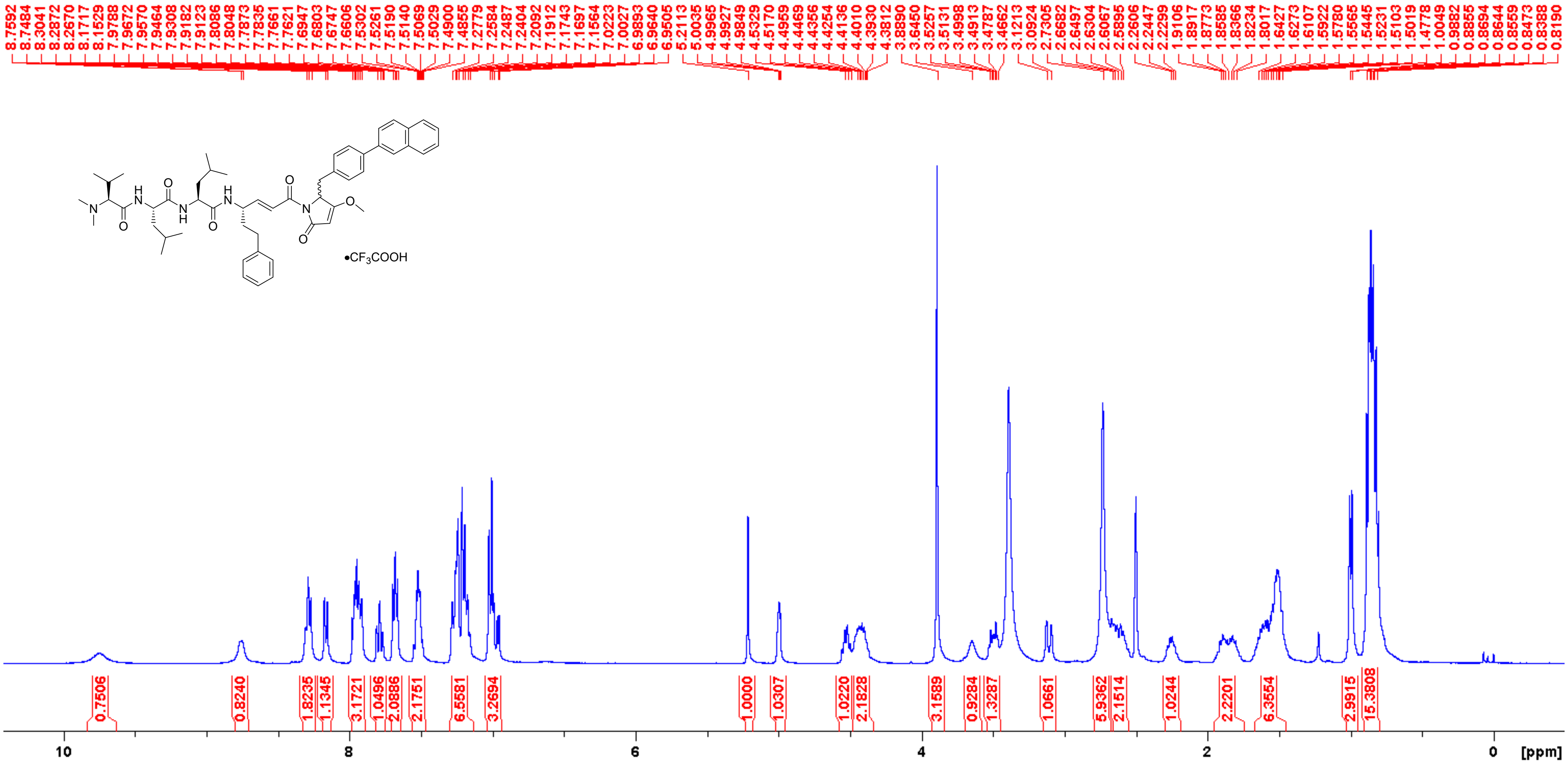

**Analogue 32** $^{13}$C NMR (100 MHz, DMSO-$d_6$)

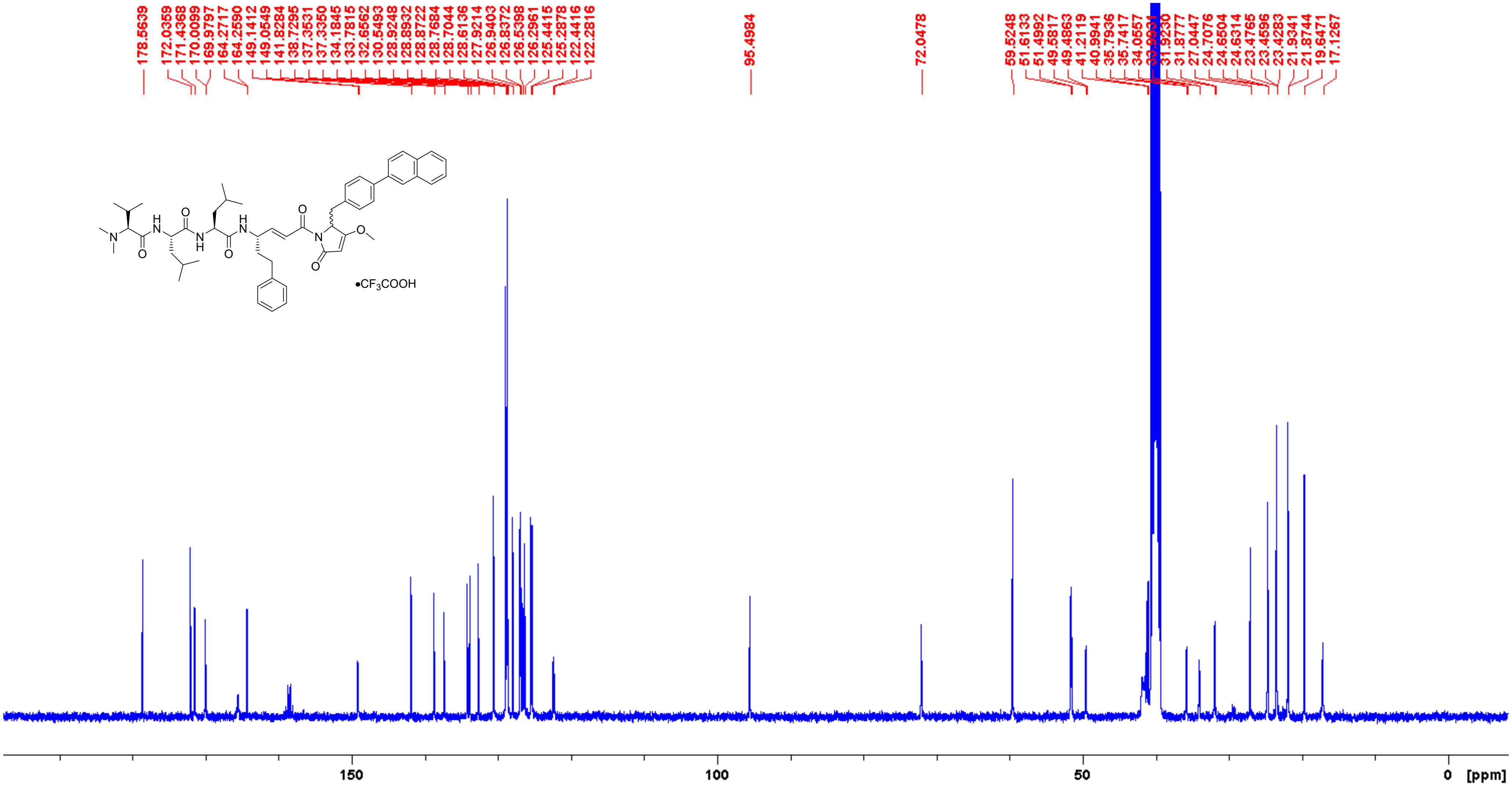

**Analogue 32** COSY (400 MHz, DMSO-$d_6$)

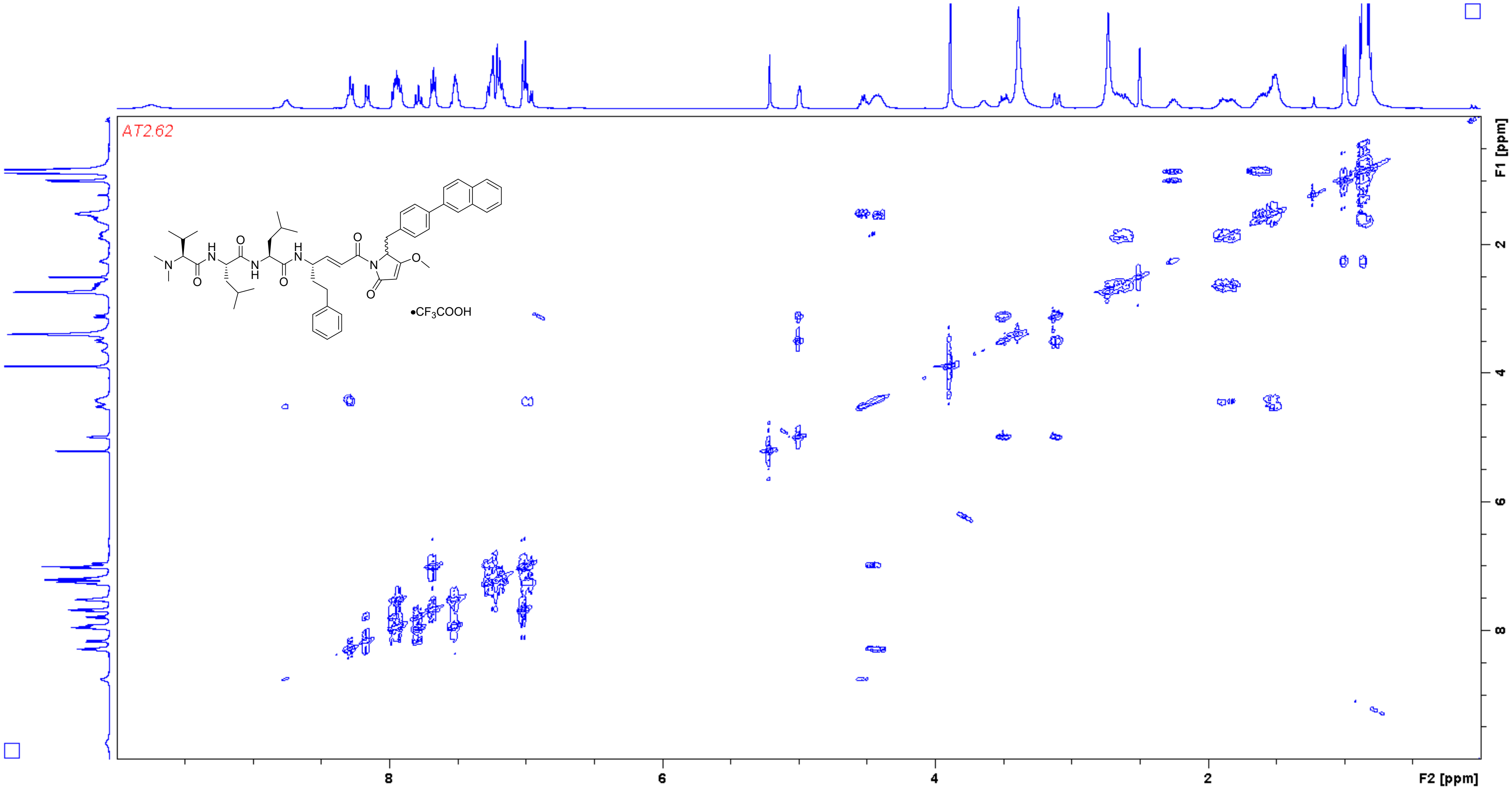

**Analogue 32** HSQC (400/100 MHz, DMSO-$d_6$)

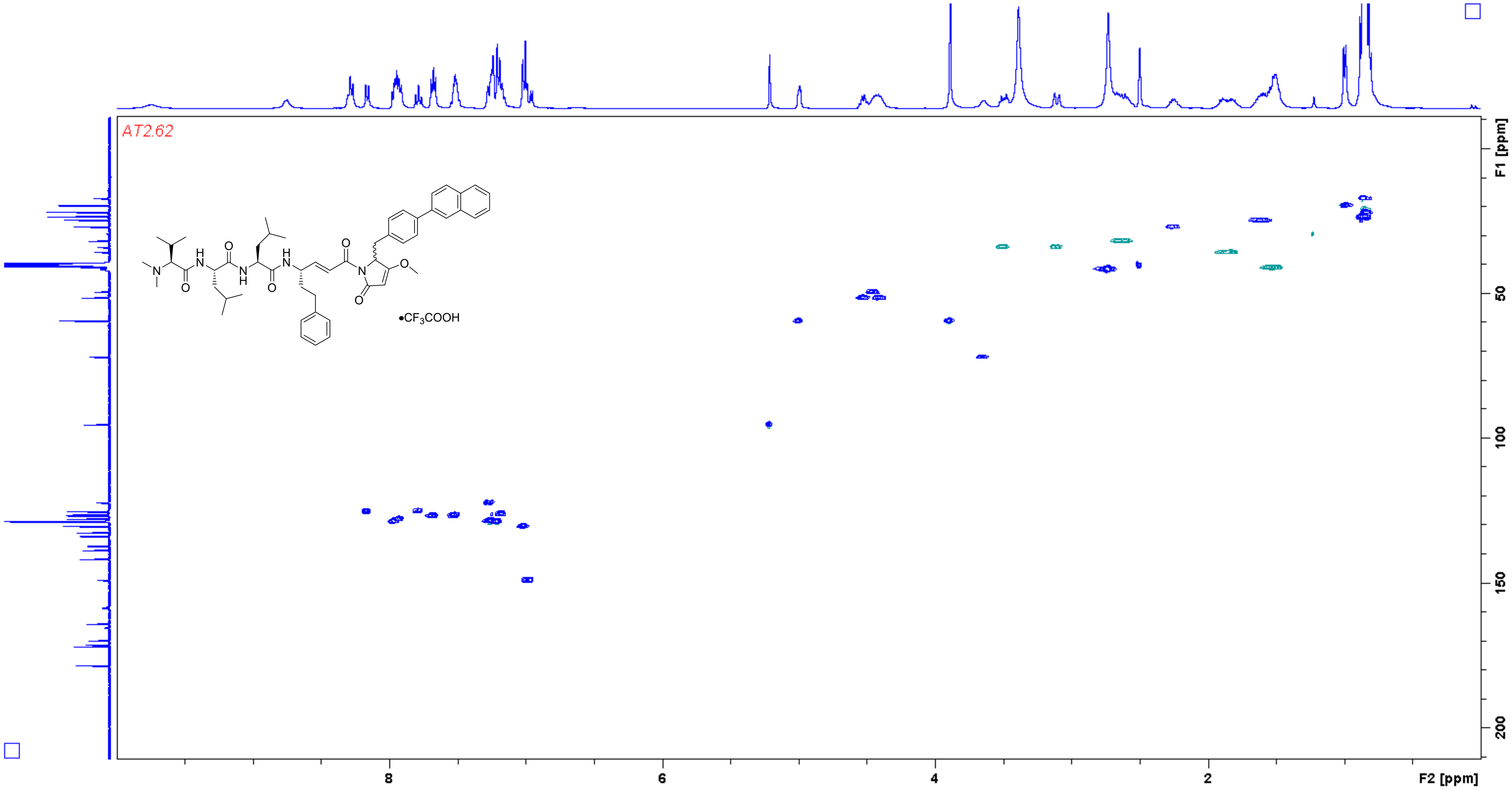

**Analogue 32** HMBC (400/100 MHz, DMSO-$d_6$)

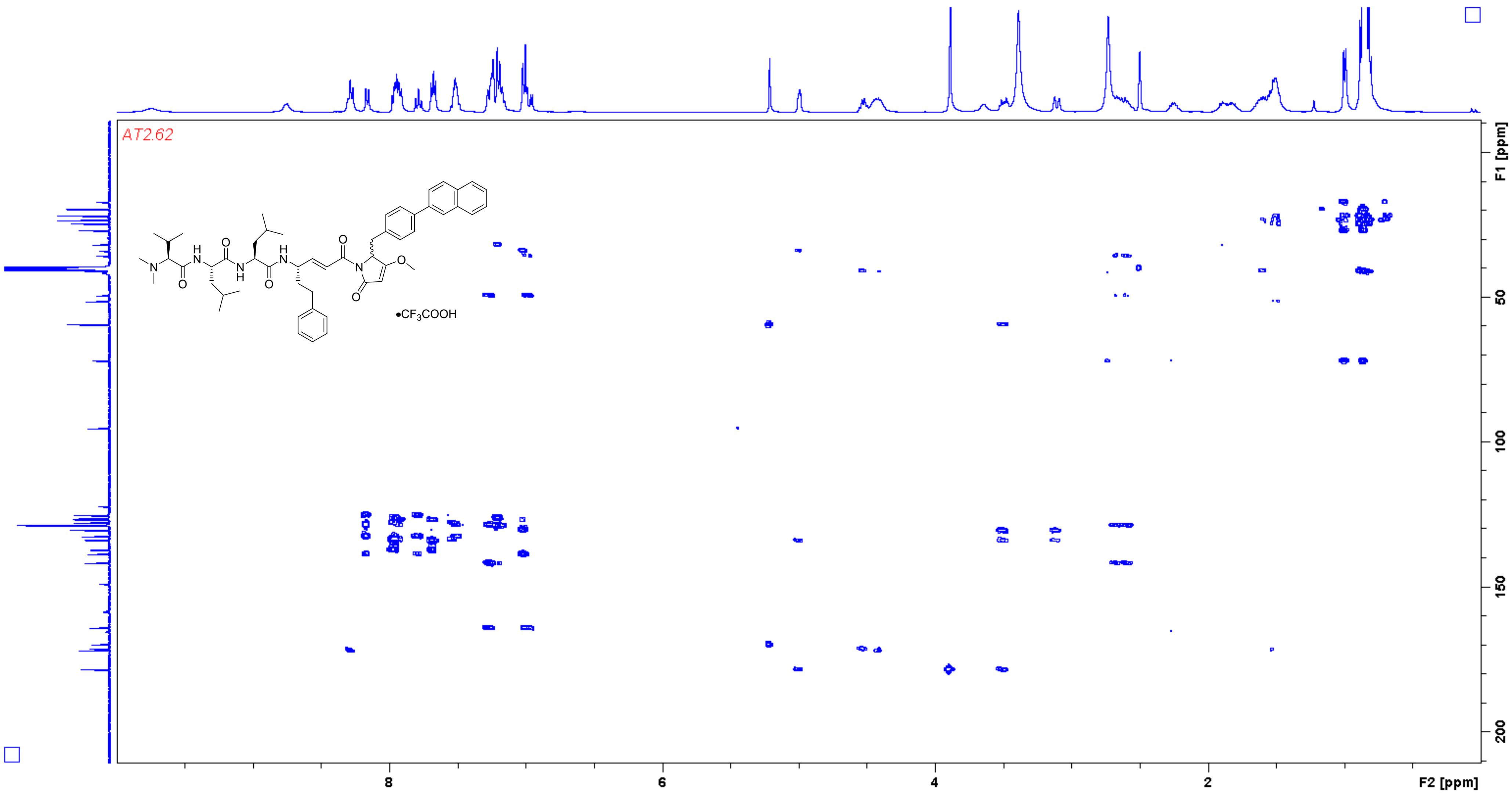

**Analogue 33** $^{1}$H NMR (400 MHz, DMSO-$d_6$)

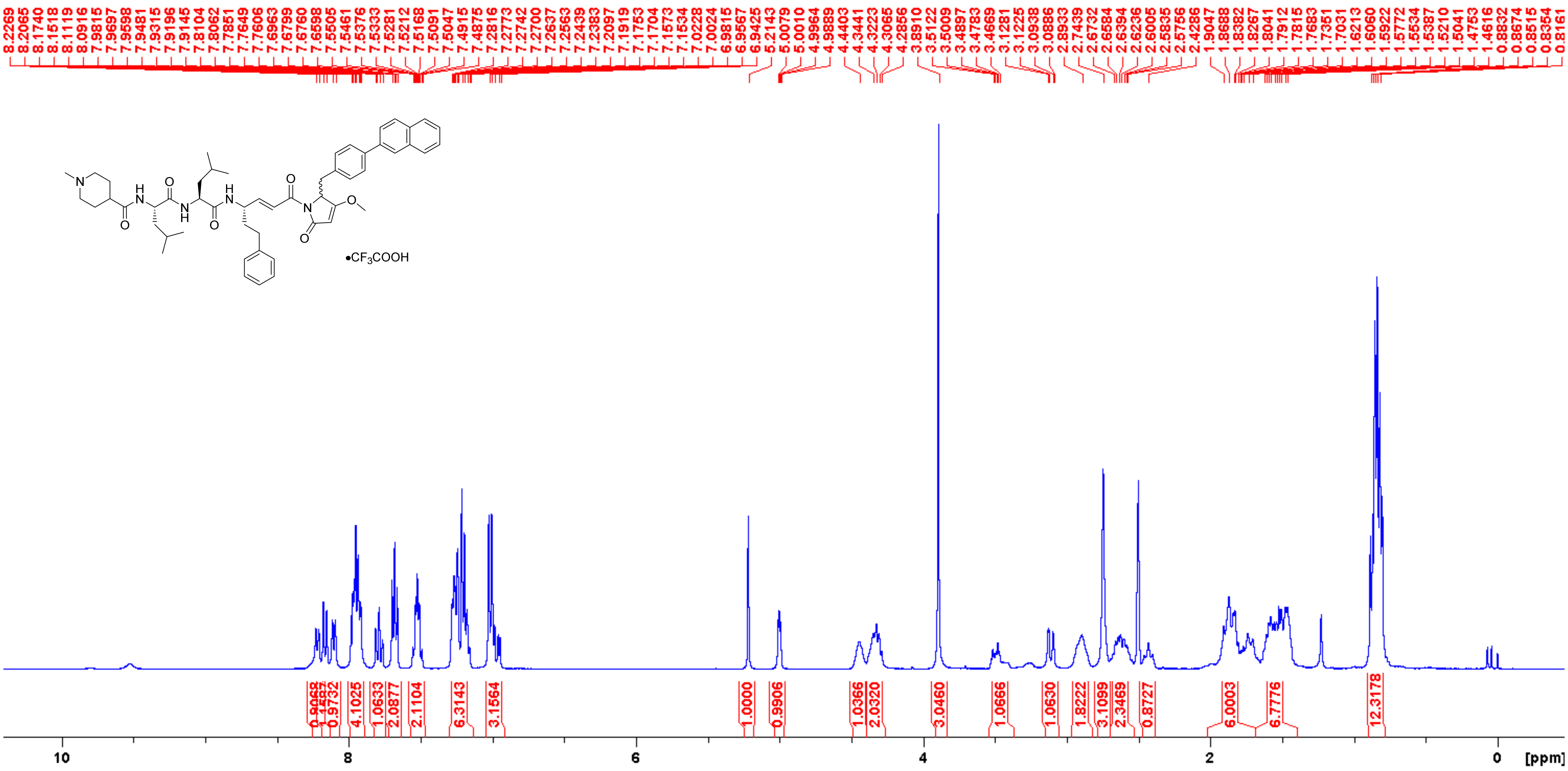

**Analogue 33** $^{13}C$ NMR (100 MHz, DMSO-$d_6$)

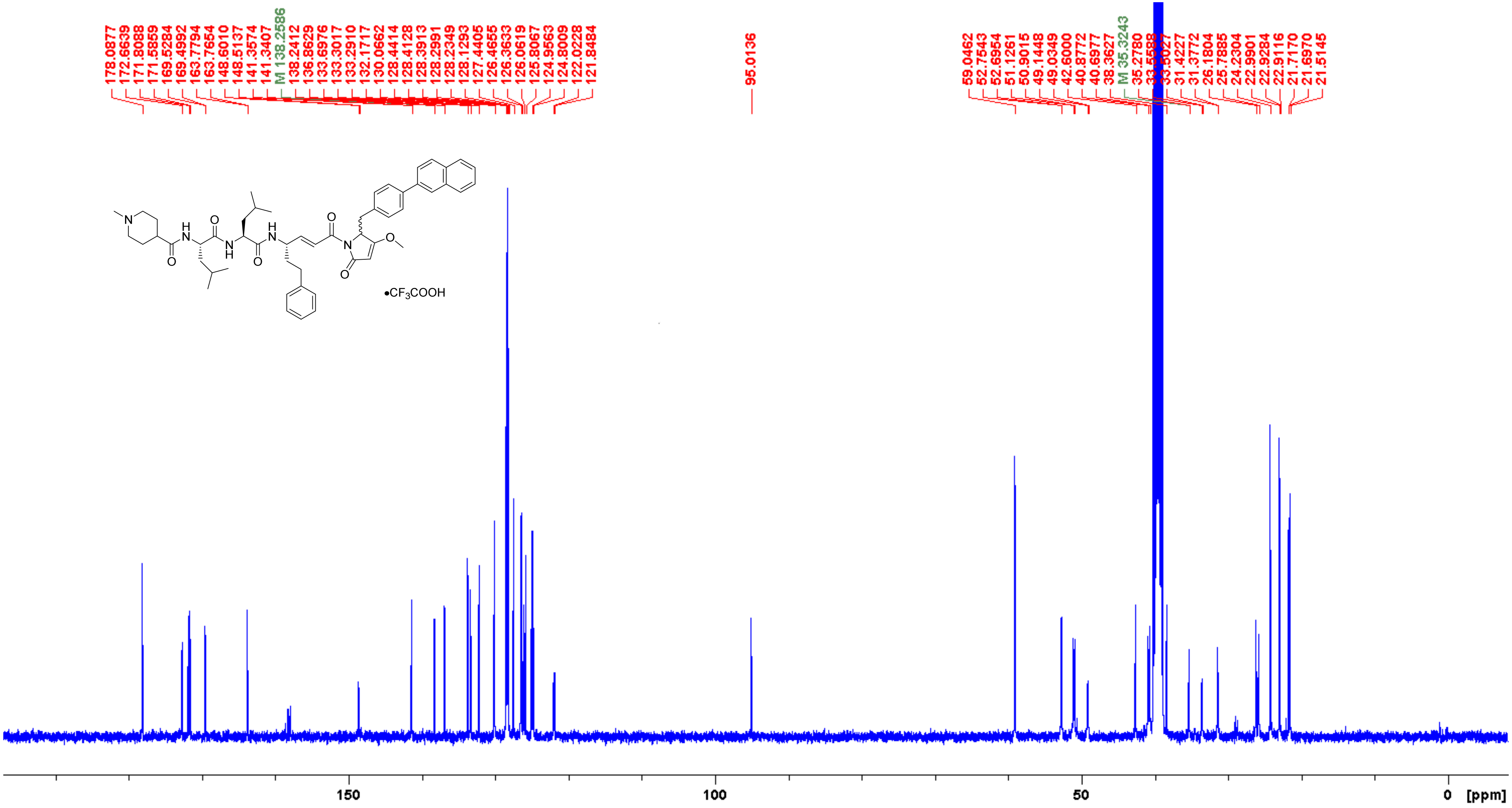

**Analogue 33** COSY (400 MHz, DMSO-$d_6$)

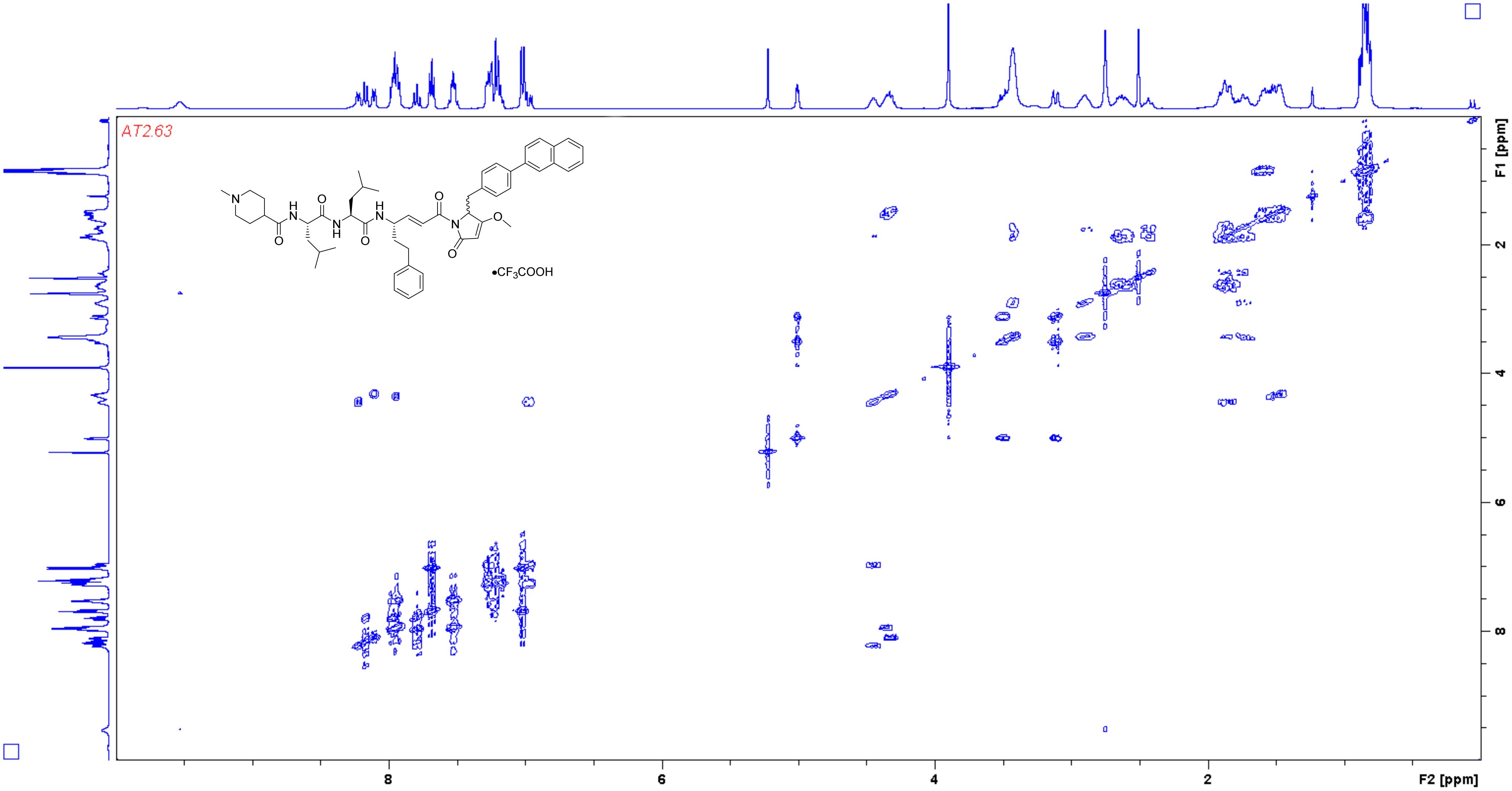

**Analogue 33** HSQC (400/100 MHz, DMSO-$d_6$)

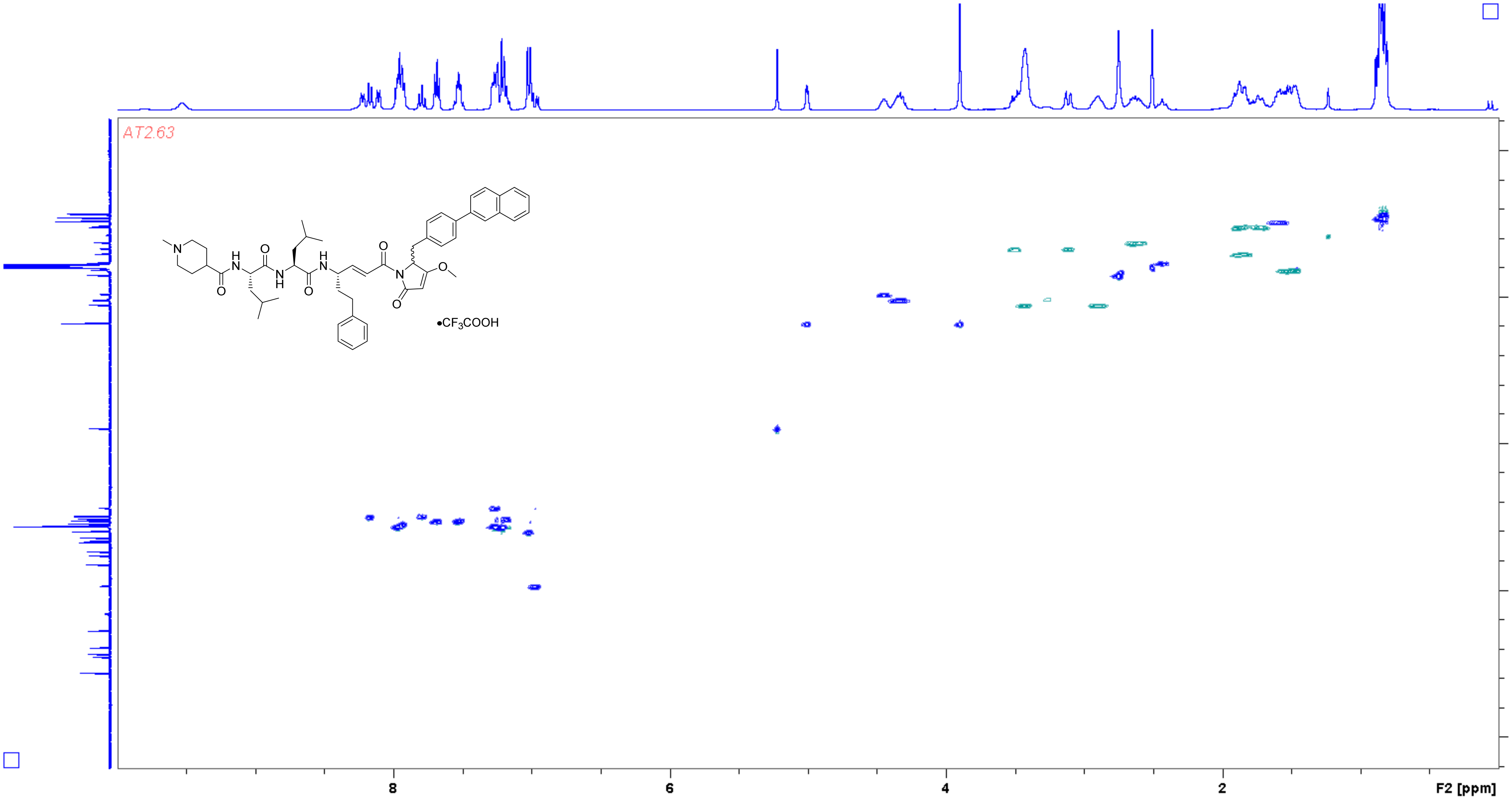

**Analogue 33** HMBC (400/100 MHz, DMSO-$d_6$)

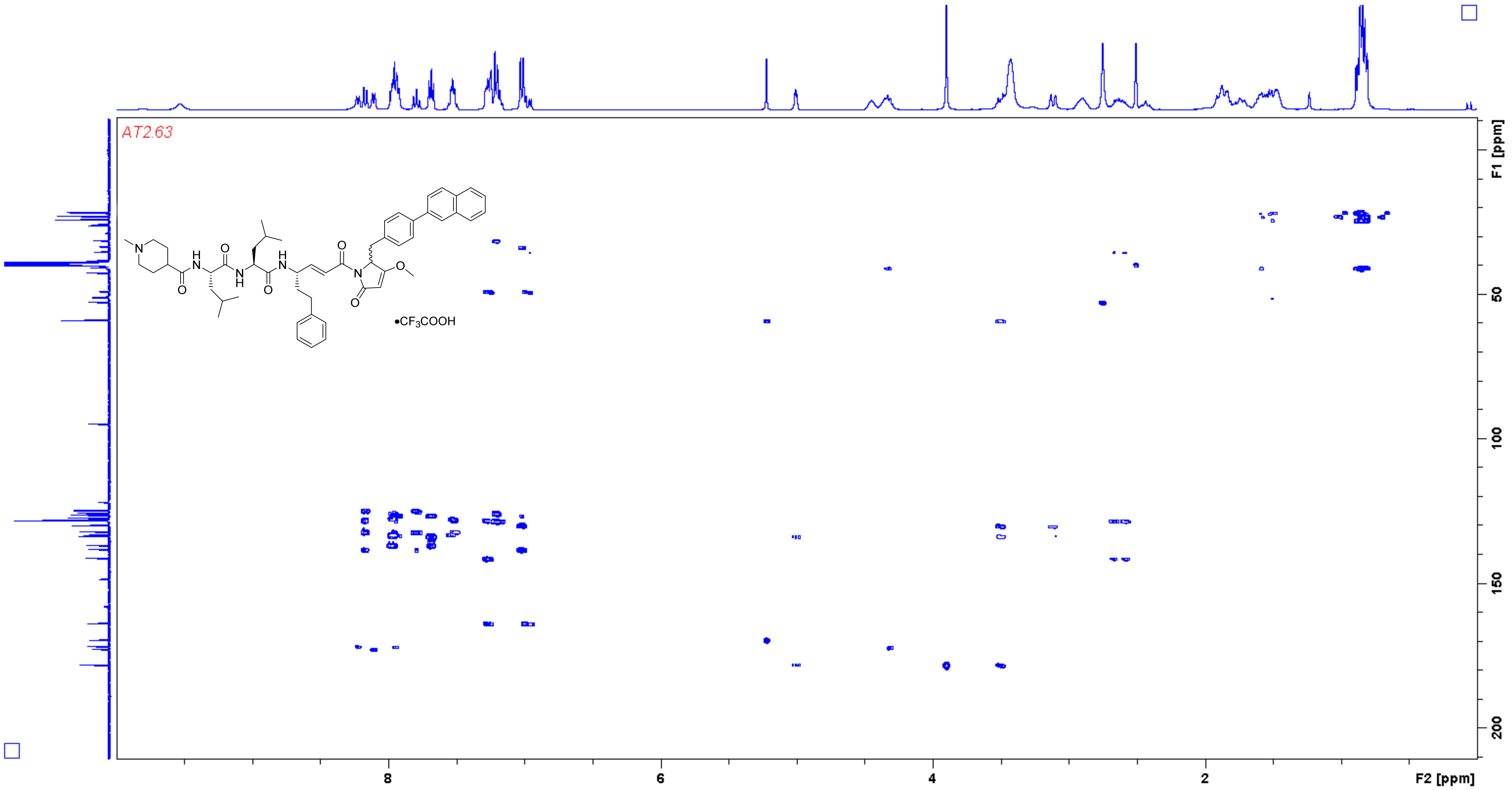
